# Tracing the spread of Celtic languages using ancient genomics

**DOI:** 10.1101/2025.02.28.640770

**Authors:** Hugh McColl, Guus Kroonen, Thomaz Pinotti, William Barrie, John Koch, Johan Ling, Jean-Paul Demoule, Kristian Kristiansen, Martin Sikora, Eske Willerslev

## Abstract

Celtic languages, including Irish, Scottish Gaelic, Welsh and Breton, are today restricted to the Northern European Atlantic seaboard. However, between three and two thousand years before present (BP), Celtic was widely spoken across most of Europe before being largely replaced by Germanic, Latin or Slavic^1–4^. Despite this rich history, how Celtic spread across the European continent remains contentious^5^. The debate is currently focused around three main models based on historical linguistics and archaeology: (1) a Late Bronze Age/Early Iron Age spread from Central Europe associated with the Hallstatt and La Tène Cultures^6–9;^ (2) a Late Neolithic/Early Bronze Age spread along the Atlantic seaboard linked to the Bell Beaker Culture^10–13;^ and (3) a Bronze Age spread from France, Iberia or Northern Italy^14–16^. Previous genomic investigations are centred around the arrival of Celtic to specific regions: Britain^17^, Iberia^18^ and Southwestern Germany^19^. Here, we utilise new genomic data from Bronze and Iron Age Europe to test how the population histories align with the three models of prehistoric spread of the Celtic languages. In line with the theory that Celtic spread from Central Europe during the Late Bronze Age to Early Iron Age, we find Urnfield-related ancestry – specifically linked to the Knovíz subgroup to have formed between 4 and 3.2 kyr BP, and subsequently expanded across much of Western Europe between 3.2 and 2.8 kyr BP. This ancestry further persisted into the Hallstatt Culture of France, Germany and Austria, impacting Britain by 2.8 kyr BP and Iberia by 2.5 kyr BP. Our findings thus agree with the model of Central European spread of the Celtic languages through consecutive expansions of the Urnfield, Hallstatt and La Tène Cultures rather than the competing models. These results demonstrate, yet again, the power of ancient population genomics in addressing long-standing debates in historical linguistics and archaeology.

## Main

Europe’s present-day linguistic landscape is dominated mainly by Romance, Germanic and Slavic languages, which expanded relatively recently during the Iron Age, mainly following the rise of the Roman Empire and the so-called Barbarian Invasions^1–4^. Prior to these events, Celtic languages constituted a dominant component of the European linguistic landscape^20,21^. Iron Age Celtic languages include Lepontic, documented in Northwestern Italy c. 2600–2100 BP, Celtiberian in Central-Northeastern Iberia c. 2200–1900 BP and Gaulish, extending across present-day France, Central Europe, Northern Italy, the Balkans and parts of Anatolia, c. 2500–1400 BP^21–23^. In the modern era, Celtic languages have retreated to the northwestern Atlantic fringe, where they persist as two groups: the Goidelic languages Irish, Manx and Scottish Gaelic, and the Brittonic languages Welsh, Cornish and Breton^24,25^.

To understand the prehistoric spread of the Celtic languages, a range of contradictory hypotheses have been formulated over the past century and still today the debate on the centre, the timing as well as the mechanisms of this dispersal remain highly contentious^26^. Previously, the late fifth-millennium BP Bell Beaker Complex (4750/4500–4000/3800 BP), which spanned much of Western and Central Europe, has been suggested as a possible archaeological context for the Celtic split and dispersal^11,27–29^. This interpretation is contradicted, however, by linguistic estimates dating the formation of Proto-Celtic, the common ancestor of all Celtic languages, to around the start of the Late Bronze Age (3200 BP)^30–33^. Thus, the current debate is focused around three main alternative models:

1. The Celtic-from-the-East model here describes the formation of Celtic in Eastern Europe and correlates its dispersal with the Urnfield Culture^34,35^ or the Hallstatt and La Tène archaeological horizons^6–9,36^. Between 2600 and 2400 BP, Hallstatt and La Tène contexts see the appearances of monumental mounds known as ‘princely burials’ that are often attributed to ‘Celtic’ elites. Genomically this period has been characterised by a period of homogenisation, suggesting high levels of connectivity across Central Europe^19^. The distribution of the Iron Age La Tène Culture shows a strong geographic correlation with that of Gaulish, ranging between France and the Balkans, and this is taken to indicate a secondary outmigration from within a larger Celtic-speaking area matching more closely the extents of the Urnfield and Hallstatt Cultures. This model thus implies linguistic and genetic continuity from Late Bronze Age Urnfield Culture to Iron Age Hallstatt and La Tène Culture.
2. The contrastive Celtic-from-the-West model instead places the formation of Celtic in Iberia and Southwestern France. Under this hypothesis, relatively undiversified Indo-European dialects entered Western Europe during the Bell Beaker period, and subsequently evolved into Celtic through the accumulation of changes diffusing along long-distance maritime networks culminating in the Atlantic Bronze Age Culture (3300–2700 BP)^10,13^. Supporting this model, the Bell Beaker period saw a north-to-south spread of Steppe-derived ancestry from the mid-fifth millennium BP, resulting in a near-complete genetic turnover in Britain and a significant impact also across the Iberian Peninsula^37,38^. This model then assumes long-term linguistic and genetic continuity from the Bell Beaker period to the Late Bronze Age/Iron Age transition.
3. A final, third model, Celtic-from-the-Centre challenges both the Hallstatt and La Tène Cultures and the Atlantic Bronze Age Culture as potential archaeological vectors for the Celtic dispersal. Notably, the presence of Celtiberian and Lepontic inscriptions outside the distributions of the Hallstatt and La Tène Cultures contradict a direct association of these archaeological phenomena with Celtic proper^39^. Instead, this model then assumes a Late Bronze Age origin in France and radiation to Britain, Spain and Italy during the third millennium BP^15^. A recent study on Britain has indeed identified a genetic transition, probably through admixture from France, between 3200 and 2800 BP, marking potential entry points for Celtic languages in Britain^17,40^.

To evaluate the above three contrastive hypotheses on the Celtic linguistic dispersal, we expand upon the results of ^41^, leveraging a filtered dataset of 4587 imputed ancient human genomes with dense sampling from Western, Southern and Eastern Central Europe where Celtic languages were spoken (Fig. 1; Supplementary Table S1.1).

**Fig. 1.**
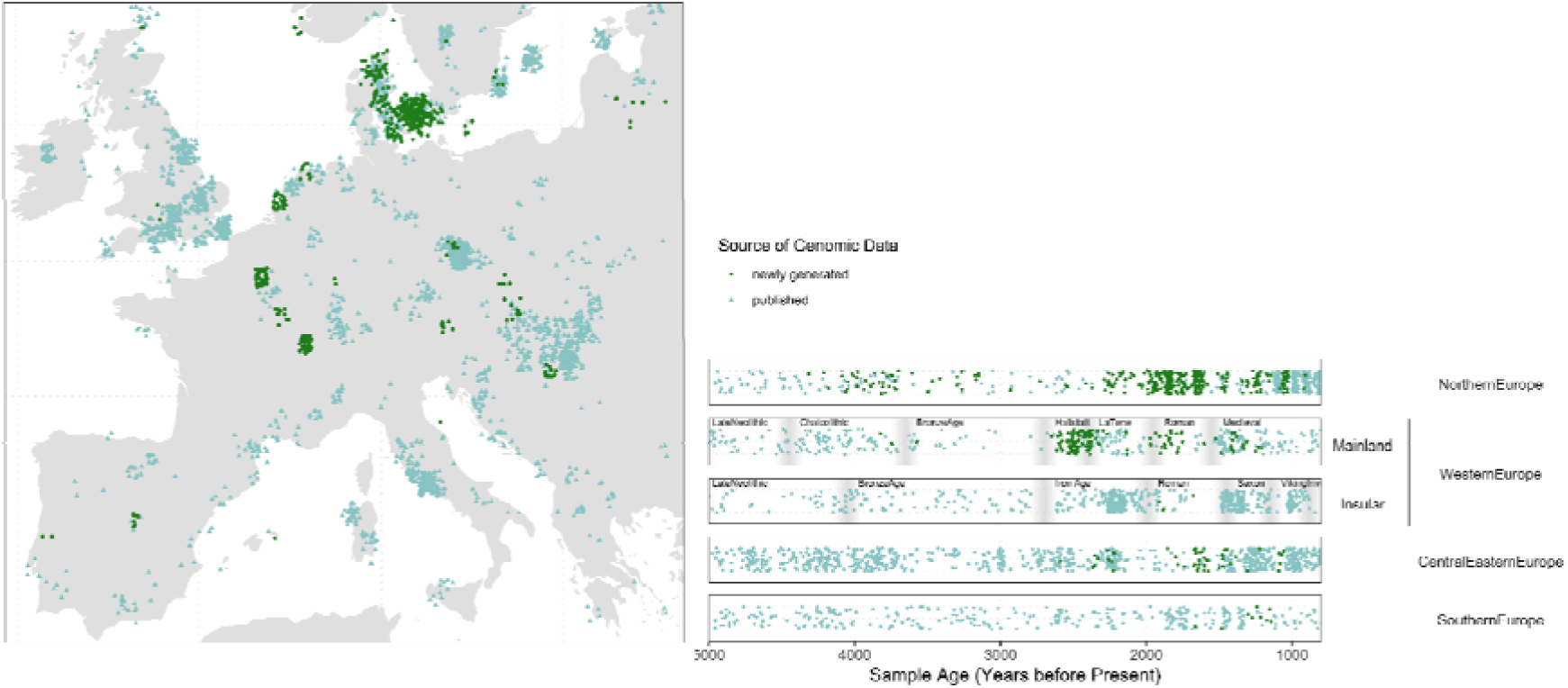
Geographic and temporal sampling of the subset of ancient individuals included in the final dataset. Newly generated (green) and published (light blue) ancient individuals from the Late Neolithic/Early Bronze Age to the Viking Age. Grey bars on the timeline represent the boundary between historical periods for Western Europe.

### Bell Beaker ancestry and Indo-European dialects

It is believed that the Indo-European dialect ancestral to the Celtic languages likely arrived in Europe along with the Yamnaya migrations from the Pontic Steppe^42,43^. In Western Europe, these migrations genetically contributed to people of the Bell Beaker Complex. By the Iron Age the region in which Celtic languages are spoken, and indeed the three proposed regions of origin of Celtic, fall spatially within the archaeological boundaries of the Bell Beaker Complex. It has therefore been argued that a subpopulation of the Bell Beaker Complex spoke an Indo-European dialect ancestral to Celtic. If correct, the formative region of Proto-Celtic should be located within the genetic boundaries of Bell Beaker-related ancestry. To test this, we performed IBD mixture modelling (methods), a supervised form of modelling which allows for ‘target’ individuals to be modelled as combinations of ‘source’ groups, allowing genetic continuity or migrations to be detected. In brief, a palette is created for every ancient individual in the dataset, representing the proportion of IBD (identity-by-descent) segments of DNA shared with all clusters in the dataset. The palette of each ‘target’ individual is then modelled as the best-fitting combination of the palettes of the ‘source’ groups, using non-negative least squares (ref).

To establish whether the three suggested formative regions for Proto-Celtic carry Bell Beaker-related ancestry, we first repeated the IBD mixture modelling analyses and ordinary global spatio-temporal kriging^41^. This revealed a strongly significant association linking individuals from Bell Beaker archaeological contexts and later Celtic-speaking regions. We find Bell Beaker-related ancestry to be present across Southern, Western and Central Europe (Supplementary Fig. S1.9; Supplementary Fig. S1.14) in the Proto-Celtic period (c. 3200 BP). This result is consistent with a Bell Beaker-mediated entry of the dialect ancestral to Proto-Celtic.

To link the emergence of Celtic with the spread of the Indo-European language family as a whole, we look to the incorporation of Steppe ancestry in the formation of Bell Beaker-related populations. Bell Beaker-related ancestry is a mixture of three distinct sources: a Yamnaya-related source from the Pontic Steppe, and two Neolithic Farmer-related sources from within Europe^44^. From the spatio-temporal kriging results, we find the Bell Beaker-related ancestry to be modelled in its highest proportions first around the Netherlands and on the British Isles, between 4400 and 4000 BP (Fig 2B; Supplementary Fig. S1.21). By this time, we already find Bell Beaker-related ancestry as far south as Iberia and Italy, albeit in small proportions (Supplementary Note S1) relative to the Neolithic Farmer-related ancestry (Fig. 2C). Between ∼4300 and ∼2600 BP, we find that Bell Beaker-related ancestry expands (Fig. 2B, E) and European Farmer-related ancestry contracts correspondingly (Fig. 2C, F). In contrast, Corded Ware-related ancestry, which in the Late Iron Age becomes associated with Germanic languages^41^, remains stable (Fig. 2A, D). We therefore conclude that the Steppe ancestry present in Bell Beaker populations likely spread south from the Netherlands along with several Indo-European dialects, some of which ultimately contributed to the formation of Celtic.

**Fig. 2.**
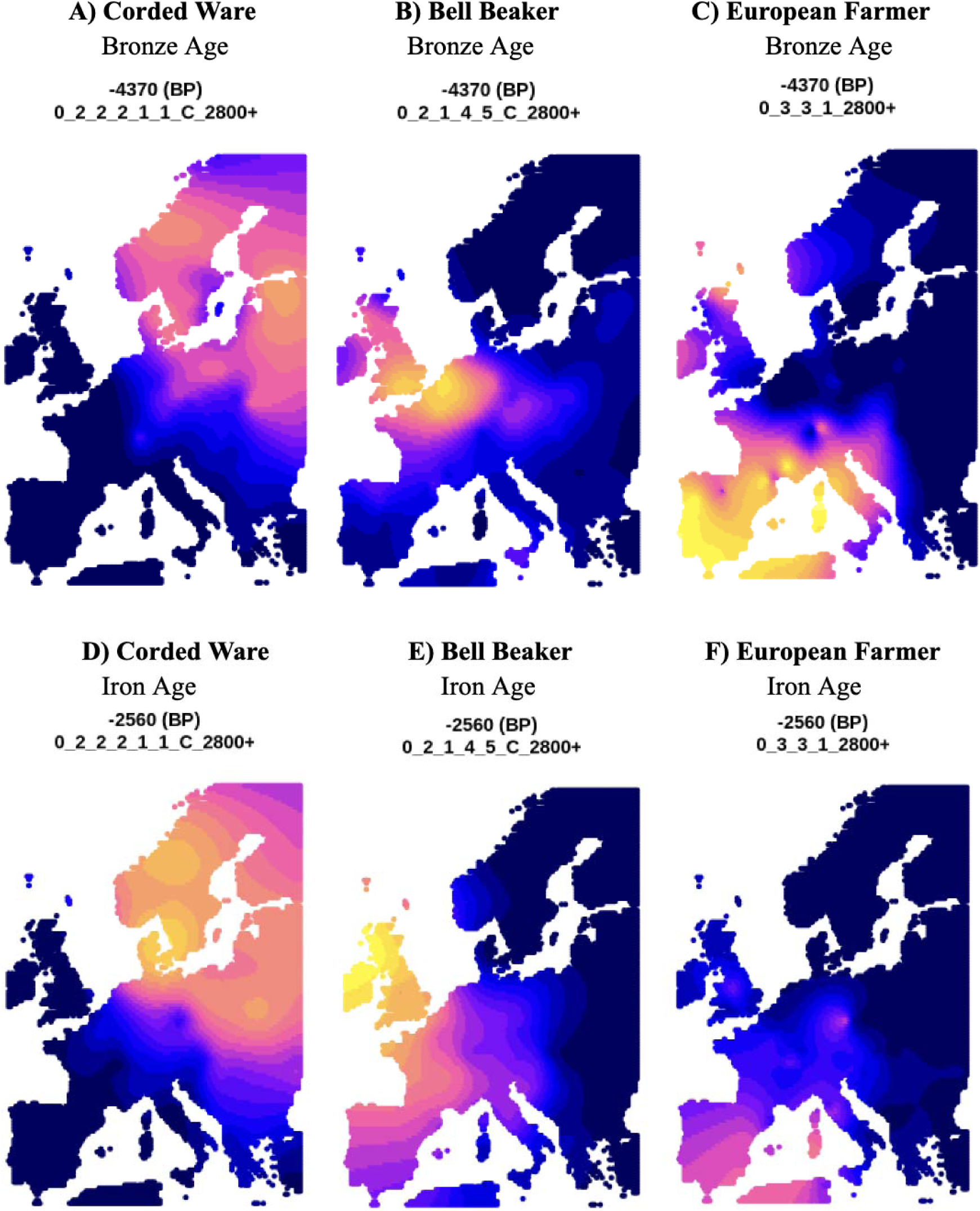
Spatio-temporal kriging results showing the distribution of Corded Ware-(A,D), Bell Beaker-(B,E) and European Farmer-related (C,F) ancestry at ∼4400 BP (upper panel) and ∼2600 BP (lower panel). Brighter colours indicate a higher proportion of ancestry modelled. Kriging was performed on the IBD mixture modelling Set C2 results.

These results are consistent with Bell Beaker-related migrations facilitating the spread of the Indo-European dialect ancestral to Celtic, as the main source of Steppe-related ancestry in the three formative regions for Celtic is Bell Beaker-related. However, to assess whether later migrations occurred consistent with any of the three regions of origin, additional resolution is required. We therefore turn to the period 4000–3000 BP.

### Middle Bronze Age population dynamics

Over a thousand years separate the presumed arrival of Indo-European dialects to Western Europe (c. 4300 BP) and the formation of Proto-Celtic (c. 3200 BP). However, how the population structure changed over this period remains unclear. To be able to evaluate the three hypotheses on Celtic origins, it is therefore first necessary to understand the demographic processes affecting Bell Beaker-derived groups of the second millennium. Importantly, detecting migrations during this time has proved difficult, due to the close genetic relation between all individuals post-dating the arrival of Steppe ancestry.

One method to infer migrations relies on detecting changes in the relative proportions of Steppe and Farmer ancestry in the millennia after the admixture initial event. For example, an increase in Farmer ancestry must result from a migration from a region in which populations carry more Farmer ancestry. Here, we expand on the use of Farmer ancestry as a proxy for migration by identifying the specific source of European Farmer ancestry. This allows us to infer the region from which later migrations occurred.

To detect in which regions admixture occurred between Bell Beaker individuals and Neolithic Farmer populations, we performed IBD mixture modelling (Supplementary Note S1), including source individuals from IBD clusters associated with Neolithic British-Irish Isles (0_3_1_2800+), France and Iberia (0_3_2_4_4_1_2800+), Italy (0_3_2_5_2_2_2_2800+) and the Czech Republic (0_3_3_1_2800+), encompassing proposed homelands for Celtic. In addition, we included a series of Neolithic-related outgroups, Bronze Age Anatolia (0_3_4_1_1_2800+), Polish Globular Amphora Culture (0_3_3_3_C_2800+) and early Anatolian Farmers (Early Anatolian 0_3_4_3_3_2_2_2800+). Relatively dense sampling now exists from across the region of interest during much of the Bronze and Iron Ages. However very few genomes exist from the Late Bronze Age as a result of the widespread practice of cremation associated with the Urnfield Culture. We therefore first focus on the peripheries of the region of Bell Beaker-related ancestry, England and the Czech Republic, where dense sampling continues throughout the Bronze and Iron Age, and France, where the previously sparse Iron Age sampling has been greatly improved.

To visualise these changes through time, we first binned samples into three Copper/Bronze Age bins (4800–4000 BP, 4000–3200 BP and 3200–2800 BP) and two Iron Age bins (2800– 2470 BP and 2470–2000 BP) and plotted the proportion of ancestry modelled for each individual in (Fig. 3.).

**Fig. 3.**
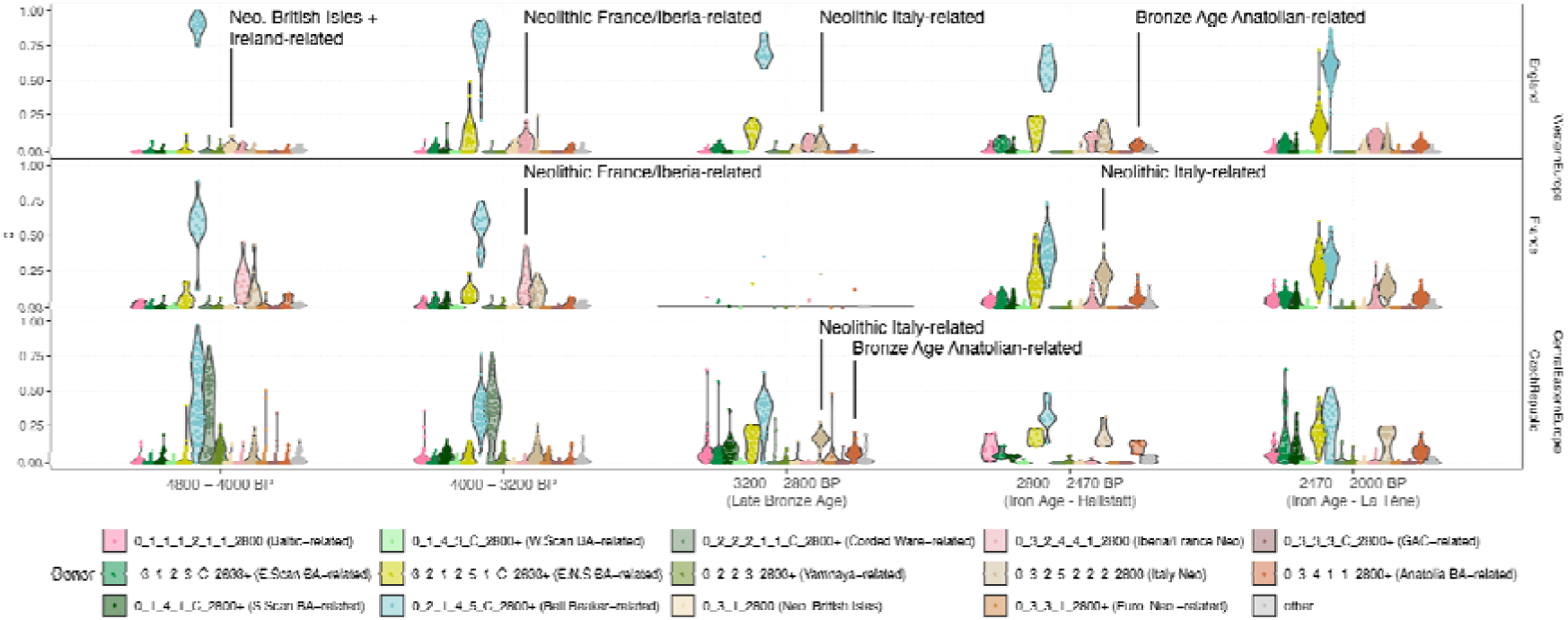
IBD mixture modelling results (Set C3) for Bronze and Iron Age England, France and the Czech Republic. The shifts in relevant Neolithic Farmer-related ancestries have been annotated. The full set of IBD mixture modelling results can be found in Supplementary Fig. S1.1.

For England, we find a series of transitions in the prominent Farmer ancestries present for each time slice (Fig. 3). Initially, between 4800 and 4000 BP, we find that individuals are modelled with a high proportion of Bell Beaker-related ancestry, and the tendency to have a slightly higher proportion of local British-Irish Isles Neolithic ancestry, relative to the other Neolithic-related ancestries. By 4000–3200 BP, the highest Farmer-related ancestry is French/Iberian Neolithic-related rather than the local Neolithic ancestry, consistent with recent studies suggesting migrations from the mainland^17,40^. This migration, specifically the Iberian connection, is further supported by evidence that the UK received copper from Iberia during this phase (3350/3250–750 BP)^16^. However, in the Late Bronze Age, we see a shift, in which the proportion of Italian Neolithic ancestry has increased to similar proportions to that of French/Iberian Neolithic. In the Iron Age, similar patterns are seen, with the additional appearance of Bronze Age Anatolian-related ancestry.

In France, we see a similar transition (Fig. 3). During the Early and Middle Bronze Age, more local French/Iberian-than Italian Neolithic-related ancestry tends to be present. By the Iron Age, the relative proportions have swapped, so Italian Neolithic-related ancestry is the highest, accompanied by Bronze Age Anatolian ancestry. Due to the lack of samples from the Late Bronze Age, the time of this transition cannot be directly measured. However, the increased proportion of Italian Neolithic-related ancestry during the Late Bronze Age on the British Isles suggests it was present in France by this time. We also in France an increase in ‘East North Sea’ ancestry going from 2800–2470 BP to 2470–2000 BP, corresponding to the Hallstatt and La Tène periods in France. This ancestry is Bell Beaker-related and found in high proportions throughout the Bronze and Iron Ages in the Netherlands^41^. The increase in France thus indicates the arrival of more northern ancestry by the La Tène period.

Further east, in the Czech Republic, we see the increase of Italian Neolithic-related and Bronze Age Anatolian-related ancestry between 3200 and 2800 BP (Fig. 3). We also note that we detect no evidence of French/Iberian Neolithic ancestry. By splitting further into the cultural phases for the region, we find that this ancestry profile in the Czech Republic occurred by 3300 BP, in individuals associated with the Tumulus Culture and continuing into the Knovíz and Hallstatt periods (Extended Data Fig. 1).

To understand how these Neolithic Farmer-related ancestries spread during the Bronze and Iron Age more broadly, we performed spatio-temporal kriging on the IBD mixture modelling results for these ancestries (Extended Data Fig. 2). For the three European Farmer-related ancestries, we see the proportion of ancestry modelled decreasing through time. However, while we see a general range reduction of the British and French/Iberian Neolithic-related ancestries, we find an increase in the geographical range of the Italian Neolithic-related and Bronze Age Anatolian-related ancestries throughout these periods.

We interpret the consistency of Bell Beaker-related ancestry and shifts in Neolithic Farmer ancestry to result from multiple admixture events between Bell Beaker-related populations and local farmers across Europe, who subsequently migrated within the region prior to the formation of Proto-Celtic.

### Migrations post-dating the formation of Proto-Celtic

The spread of Celtic under all three hypotheses occurs between the formation of Proto-Celtic from 3200 BP and the time of its historical distribution, around 2500–1500 BP. To identify the centre of spread of the descendent Celtic languages, we attempted to detect migrations impacting historically Celtic-speaking regions from c. 3200 BP.

To model the migrations directly, we included Bronze Age sources from the regions in which the distinct Farmer ancestries associated with the migrations were found. We initially chose Bronze Age individuals from Southwest and Southeast Europe, to track general movements. From Southwest Europe, we included individuals from France and Iberia, who are modelled primarily as Bell Beaker- and the local French/Iberian Bronze Age-related ancestry (0_3_2_2_2800+) and are relevant to the Middle Bronze Age migrations to the British-Irish Isles. From Southeast Europe, we included individuals from Hungary/Serbia (0_3_4_2_2_C_2800+), who are relevant to the appearance of Italian Neolithic and Bronze Age Anatolian-related ancestry in Western Europe by the Iron Age. Consistent with the results found from using the Farmer-related ancestries as a proxy, we find Bronze Age French/Iberian ancestry appearing in England during the Middle Bronze Age, and the Hungarian/Serbian Bronze Age reaching widespread distributions during the Iron Age (Fig. 4). In the Czech Republic, we find almost all individuals being modelled with a large proportion of Hungarian/Serbian ancestry during the Late Bronze Age.

**Fig. 4.**
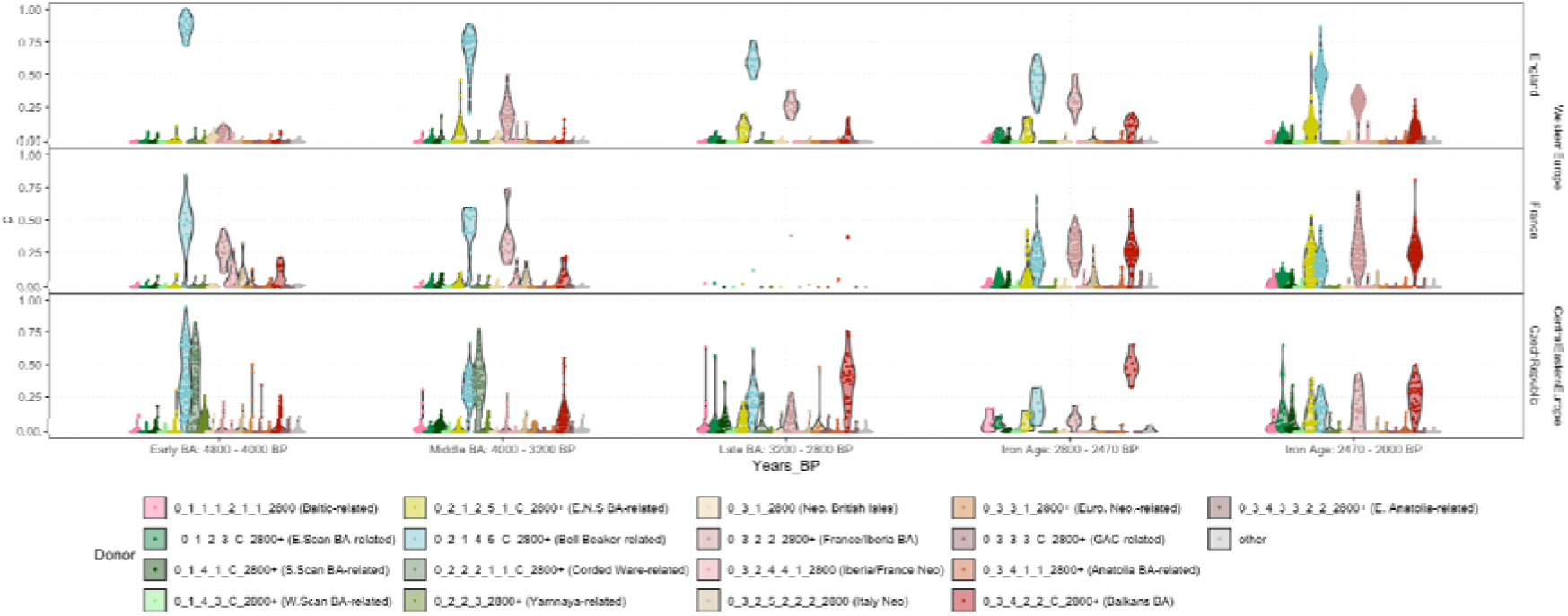
IBD mixture modelling results for England, France and the Czech Republic during the Bronze and Iron Ages, highlighting the Southeast (Hungarian/Serbian) and Southwest (France Iberian) European Bronze Age ancestries.

To understand how these Bronze Age-related ancestries spread during the Bronze and Iron Age more broadly, we performed spatio-temporal kriging on the IBD mixture modelling results for these ancestries (Extended Data Fig. 4). The full set of IBD mixture modelling results can be found in Supplementary Fig. S1.1.

Next, we included individuals from the Late Bronze Age from the Czech Republic, associated with the Urnfield subgroup of the Knovíz Culture, as a source, who were modelled above with high proportions of Hungarian/Serbian ancestry (Extended Data Fig. 1). In the early Iron Age (2800–2470 BP), we find this ancestry modelled across Western, Southern and Eastern Central Europe in varying proportions, complemented by more local sources (Fig. 5; Extended Data Fig. 3.). In England, the highest proportion tends to be modelled as Bell Beaker-related, followed by Knovíz-related and French/Iberian Bronze Age-related. In contrast, on the mainland, the proportion of ancestry modelled as Bell Beaker-related tends to be low or absent, with the Steppe ancestry in these individuals better modelled by the other Bronze Age sources, i.e. Bronze Age Knovíz-related, French/Iberia and Hungary/Serbia.

**Fig. 5.**
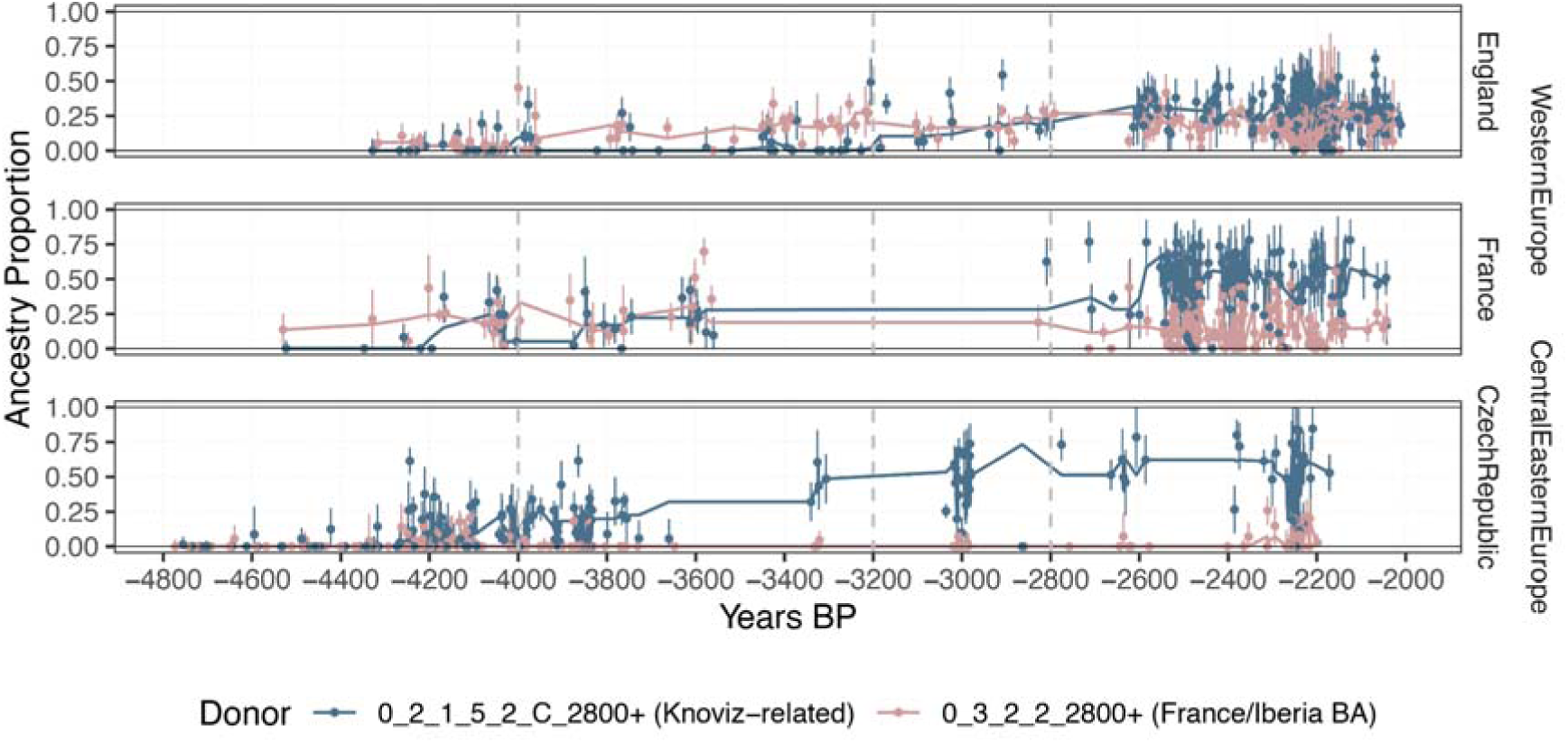
IBD mixture modelling results for England, France and the Czech Republic during the Bronze and Iron Age, highlighting the spread of Knovíz-related ancestry, relative to French/Iberian Bronze Age ancestry. The trendline shows the rolling median with a window of 5.

Compared to England, the impact of Knovíz-related ancestry is particularly high in France, Germany and the Czech Republic. In Austria, we note considerable diversity at the eponymous Hallstatt site, with individuals modelled with particularly high proportions of either Knovíz or Hungarian/Serbia Bronze Age-related ancestry, more similar to Hungary, Slovenia and Slovakia, where the Hungarian/Serbian Bronze Age-related ancestry is modelled in high proportions.

The results here indicate a migration from Eastern Central Europe, occurring between 3200– 2800 BP. The genetic impacts diminish the further west that they are detected, but can be measured as far west as Iberia and the British Isles. The Knovíz-related source that models this migration is found in the Czech Republic from 3300 BP. However, we note that this population is distinct from the earlier Únětice people from this region before 3800 BP (Supplementary Note S1; Extended Data Fig. 1) and has links further south and east, evident from the appearance of Italian Neolithic Farmer-related and Bronze Age Hungary/Serbia-related ancestry. Thus, we observe that the population of the Únětice Culture does not significantly contribute to the ancestry of the successive Hallstatt Culture.

Additionally, we find some level of support to these migrations from Y-chromosome haplogroups. While R-P312 is closely linked spatiotemporally with Bell Beaker ancestry, its subhaplogroups exhibit great geographical structure, particularly in the Bronze Age, with R-L21 dominant in Britain and Ireland^17,45^, R-DF27 in Iberia and France^18,38,46^ and R-U152 in Central Europe and Northern Italy^37,43,47,48^ (Supplementary Note S4). This distribution provides a framework for understanding later population movements. Consistent with IBD mixture modelling results, we see R-DF27 appearing in England during the Middle Bronze Age^17,49^, likely from France/Iberia, and R-U152 reaching both England and France at the end of the Bronze Age^17,49,50^. In France, most occurrences are associated with Hallstatt and Gaulish contexts, further reinforcing this connection.

To confirm the occurrence of these migrations using more conventional methods, we ran a series of tests using qpAdm (Supplementary Note S2) and chromopainter (Supplementary Note S3). We find that the results are not contradictory, and are broadly consistent with the IBD mixture modelling results. We therefore conclude that in England, migrations from at least three distinct populations occurred. Following the first arrival of Steppe ancestry with the Bell Beaker Complex, we find Middle and Late Bronze Age groups can be modelled only with additional ancestry from Mainland Southern and Western Europe. In contrast the Iron Age population requires ancestry from more Eastern Central Europe to be well modelled, likely occurring at the end of the Bronze Age. In France, a similar result is found; in the Iron Age, populations can only be modelled when including a Bronze Age source from Eastern Central Europe, indicating that here too, migrations occurred from the east. The only migration at a time consistent with the three hypotheses has origins in Eastern Central Europe.

## Discussion

We have tested three competing hypotheses on the spread of the Celtic languages. We find the Knovíz-related expansion to impact the regions in which Celtic languages are attested during the Iron Age^51^. The timing, origins and direction of this expansion are consistent with the Celtic-from-the-East model. Importantly, the results do not support the Celtic-from-the-West or Celtic-from-the-Centre models, as this would require a language diffusion in a direction opposite to that of the genetic ancestry.

The spread of this genetic ancestry occurring at this time is supported by archaeological evidence (Supplementary Note S6), which indicates a westward expansion of Urnfield Culture (3300–2750 BP) from Eastern Central Europe into France and Northeastern Spain^52,53^ Several factors have been proposed as contributing to the expansion of the Urnfield Culture, including advancements in metalworking, new agricultural techniques and the introduction of millet, together fostering economic stability and demographic surplus^51,54,55^. Our findings support the expansion of the Urnfield Culture as representing the introduction not only of a new cultural and ritual practice, but also of a new population, thus providing resolution to a longstanding archaeological debate^56^.

The demographic impact of the Urnfield Culture allows us to provide new detail to the divergence of the Celtic languages. We detect continuity of Knovíz ancestry across the Urnfield, Hallstatt and La Tène Cultures, spanning a ∼1000-year period. However, how the expansion of each of these three cultural groups is related to specific Celtic languages (Supplementary Note S5) remains unclear. The interpretation of our findings suggests the following new model.

We associate the first expansion, that of the Urnfield Culture, with the spread of archaic continental Celtic varieties. This is corroborated archaeologically by the Lepontic inscriptions of the Alpine Golasecca Culture (2900–2400 BP), which itself developed from the local Canegrate variant of the Urnfield Culture. The link between Celtiberian and the appearance of the Urnfield Culture in Iberia^18,57,58^ is supported by our detection of Knovíz ancestry in this region by 2500 BP, in addition to preexisting Bell Beaker ancestry. Additional ancient genomes from Iberia would allow for the arrival of Knovíz-related ancestry during the Urnfield period to be tested directly.

In contrast the transition to Hallstatt can be seen primarily as an internal transformation among the existing population without major migrations. The collapse of the Late Urnfield Culture, replaced by the Hallstatt Culture, coincided with a major climatic cooling event called the Little Ice Age (2800 BP) as well as the introduction of Iron technology. Within the core Hallstatt region, this transition is more suggestive of a reorganisation of settlements, rather than migrations. However, recent archaeological evidence documents the westward expansion of Hallstatt C from the Rhine into England and Ireland^59,60^. This Hallstatt expansion to Britain can potentially be associated with dialects ancestral to Insular Celtic. However, at present, Iron Age genomes from Ireland do not exist, preventing us from identifying the exact processes behind the emergence of Irish.

The third and final expansion, of the La Tène Culture, has often been linked to the expansion of the Gaulish language, based on classical sources and inscriptional evidence^61,62^. The archaeologically attested formation of La Tène Culture in between the Moselle and the Upper Rhine and the subsequent spread towards Eastern and Southern Europe, including the Balkans, occurred as a result of internal revolts in resistance to the power and wealth of the Hallstatt centres^51^. Indeed, we see a subtle shift after 2470 BP in France, in which the more northern East North Sea ancestry is present in higher proportions. This is consistent with previous studies showing the arrival of more Northern European ancestry into Central and Western Europe at this time^19^. This result corresponds to the late spread of Gaulish occurring from within the broader Celtic region from around 2470 BP.

Finally, our results additionally allow us to link the emergence of Celtic in Central Europe to the broader dispersal of the Indo-European language family from the Pontic-Caspian Steppe. The migration that brought Steppe ancestry into the Bell Beaker region is believed to have introduced multiple Indo-European dialects, including dialects ancestral to Celtic (Supplementary Note S5). One archaeological hypothesis on the formation of the Bell Beaker Culture is that it occurred along the Rhine, incorporating cultural and genetic influences from east and west^63,64^. This region lies at the western peripheries of the Corded Ware Complex.

However it remains to be demonstrated whether the Steppe ancestry arrived to the Lower Rhine via Corded Ware populations in northern Europe, or directly to the Upper Rhine from Yamnaya populations in Southeastern Europe. The presence of Globular Amphora ancestry in Corded Ware and Bell Beaker populations^41^, as opposed to its absence in Yamnaya populations, points to the northern route for Steppe ancestry of Bell Beaker populations.

Providing further support for a northern route, we find the earliest individuals with high proportions of Bell Beaker-related Steppe ancestry around the Netherlands. We conclude that these processes likely played a decisive role in the linguistic formation of the Celtic branch out of the Indo-European language family after its spread to Europe.

In summary, our findings support that the later Celtic languages formed in, and spread from, Bronze Age Eastern Central Europe in association with the westward expansion of the archaeological Urnfield Culture, continuing into Hallstatt and La Tène Cultures^6,34,65^. We additionally show this process to have taken place within the broader Bell Beaker region, providing a link between the arrival of Indo-European languages in Europe and the Celtic languages spoken historically and still today. Thus, the present investigation of the emergence and early spread of the historically known Celtic languages demonstrates the power of past population genomics in addressing a long-standing debate in historical linguistics.

## Methods

### Dataset preparation

The 578 new ancient individuals included in the analyses here are part of a larger dataset of 712 ancient shotgun sequenced genomes from ^41^. Details on the sampling, data generation, quality checks, site description and detailed metadata can be found in ^41^. In brief, ancient samples were visually identified as suitable for processing. Samples were drilled, pre-digested ^66^, extracted ^43^, built into libraries ^67,68^ and sequenced on Illumina Hiseq 4000 and Novaseq 6000 platforms. Sequenced reads were mapped against the human reference genome (hg19), and authenticated based on the length of reads, damage patterns, endogenous rates and presence of contamination. All samples with a genomic coverage greater than 0.1X were imputed, and close relatives identified using ngsRelate (v2)^69^ were excluded for IBD-based analyses.

Genomes were imputed using Glimpse^70^ against the 1000 Genomes (1000G) v5 phase 3 dataset^71^. Shared IBD segments were called using IBDseq ^72^ with default parameters. Segments with a LOD score <3 and hotspot regions^44^ with excess sharing were excluded. A detailed description can be found in ^41^.

### IBD mixture modelling

To identify fine-scale structure, we applied IBD mixture modelling^44^. The use of non-negative least square modelling has been routinely applied using chromopainter chunks shared between ancient source and target sets^73^. However recent studies have shown that using shared IBD segments between ancient individuals is effective at recovering finescale detail^41,44^. We relied on the dataset and IBD clustering from ^41^. The cluster IDs for all individuals included can be found in Supplementary Table S1.1. The 4587 ancient individuals in the dataset were determined to be suitable for IBD mixture modelling based on a series of filtering steps for genomic coverage (0.1X for shotgun sequenced data, 1X for 1240K capture data), average genotype probability for the 1240K captures sites (0.9) and the removal of close relatives. The final dataset included 690,211 sites, being those that intersected with the sites 1000 Genomes panel and the 1240K sites, that pass the 1000 Genomes mappability mask and had an imputation INFO score greater than 0.5.

For each of the 4587 ancient individuals in the dataset, a palette was created, representing the proportion of sharing with all other clusters, and with the other individuals in their own cluster. The proportions were normalised by the number of individuals in each cluster. To account for the fact that the sharing within the individuals own cluster has n-1 individuals, sharing with all other clusters excludes one individual at random. As a result, clusters with a single individual do not contribute to other individuals palettes. All segments greater than 1cM, with a LOD score greater than 3, that did not originate from a hotspot of IBD sharing^44^ were included in the palettes. IBD mixture modelling was then undertaken using these palettes, by modelling the target individual using non-negative least squares^74^, as described in^44^. Standard errors of the ancestry proportions were calculated using a weighted block jackknife, leaving out each chromosome.

A strength of this method is that target palettes are modelled as combinations of potential source palettes, meaning information from all other individuals in the dataset is incorporated into the modelling, rather than relying solely on the sharing of segments between the source and target individuals. When applied to densely sampled regions like Holocene Western Eurasia, sources and targets separated by 10,000 years continue to be modelled reliably^41^.

### Spatio-temporal kriging

To understand the trends in ancestry through time, we performed spatiotemporal kriging^75^ to interpolate IBD mixture modelling results for points in time and space where genetic data is not available. For the ancestry of particular relevance for Set 2 (Corded Ware-, Bell Beaker- and European Farmer-related) and Set C3 (Neolithic British Isles/Ireland-, Neolithic France/Iberia-, Neolithic Italy- and Bronze Age Anatolia-related), we fit spatiotemporal variograms via a metric covariance model^76^.

### qpAdm analyses

qpAdm analyses were undertaken on the 690,211 sites of the dataset described above, using admixtools2^77^. Populations were placed into three sets: left_fixed, right_fixed or rotational (Supplementary Table S2.4). The ‘left_fixed’ population was included as a potential source in all qpAdm analyses. The ‘right_fixed’ population was included as outgroups in all analyses. In 1 population models, all ‘rotational’ populations were included as outgroups. For 2 population models, one ‘rotational’ population was placed together with the ‘left_fixed’ population as potential sources, and all others as outgroups, in all possible combinations. For 3 population models, two ‘rotational’ populations were placed together with the ‘left_fixed’ population as potential sources, and all others as outgroups, in all possible combinations.

## Data availability

Sequence data for the new 578 ancient genomes from the accompanying study^41^ can be found in the ENA under accession: PRJEB87274. The public 4009 ancient genomes included in the dataset are listed in Supplementary Table S1.1, with additional details provided in Supplementary Note S4 of the accompanying paper^41^. All maps were generated in R (v4.2.3), using ‘map_data’ from ggplot2 (v3.5.0.9000)

## Methods References

References 66 - 77.

## Acknowledgements

GK, JK, JL and MS are supported by Riksbankens Jubileumsfond (grant M 21-0018 for the 6-year programme MARITIME ENCOUNTERS: a counterpoint to the dominant terrestrial narrative of European prehistory). KK is supported by Riksbankens Jubileumsfond (M16-0455:1, Rise II). The Lundbeck Foundation GeoGenetics Centre is supported by grants from the Lundbeck Foundation (R302-2018-2155, R155-2013-16338), the Novo Nordisk Foundation (NNF18SA0035006), the Wellcome Trust (WT214300), Carlsberg Foundation (CF18-0024), the Danish National Research Foundation (DNRF94, DNRF174), the University of Copenhagen (KU2016 programme) and Ferring Pharmaceuticals A/S.

## Author contributions

EW initiated the study. HMC, GK, EW led the study. HMC, GK, MS, EW conceptualised the study. KK, MS, EW supervised the research. GK, JK, JL, KK, MS, EW acquired funding for research. HMC, TP, WB, MS were involved in data analysis. HMC, GK, JK, JL, KK, EW drafted the main text. HMC, GK, TP, WB, JK, JL, KK, MS drafted supplementary notes and materials. HMC, GK, TP, WB, JK, JL, JPD, KK, MS, EW involved in reviewing and editing drafts.

## Ethics declarations

The authors declare no competing interests

## Additional Information

Supplementary Information is available for this paper. Reprints and permissions information is available at www.nature.com/reprints.

## Extended Data Figures

**Extended Data Fig. 1.**
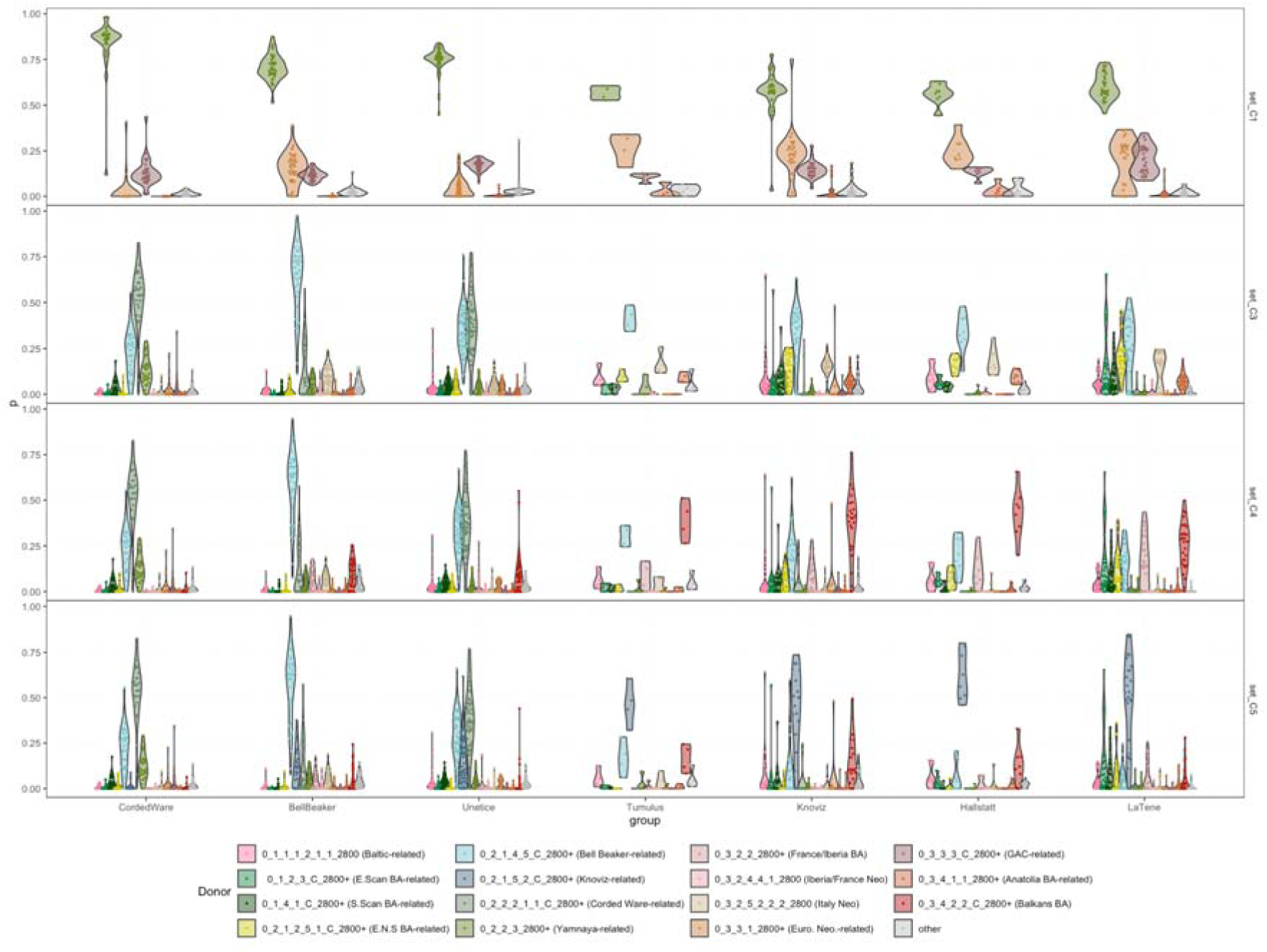
IBD mixture modelling results for samples associated with archaeological cultures/complexes from the Czech Republic for different archaeological periods. Cultures are ordered from oldest (left) to youngest (right). The top panel (Set C1) highlights decreasing Yamnaya ancestry (olive green) and varying Farmer ancestry through time. The second panel (set 3) highlights transitions between Corded Ware (grey-green) and Bell Beaker (light blue) ancestry, and the various Farmer-related ancestries. The third panel (Set C4) highlights the appearance of the Balkans Bronze Age (red) ancestry from the Tumulus Culture onwards. The lower panel (Set C5) models the Knovíz-related ancestry (dark blue) directly.

**Extended Data Fig. 2.**
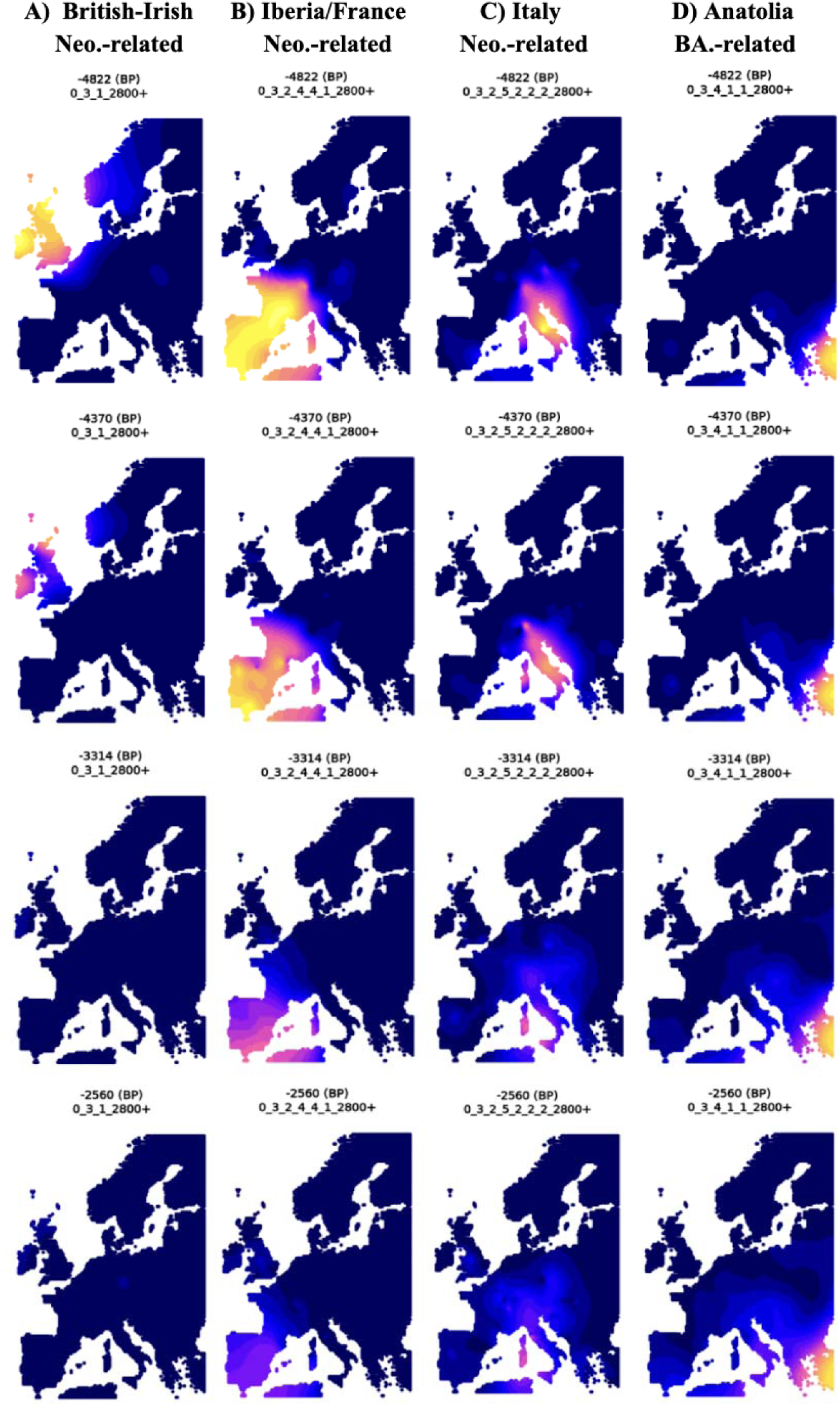
Spatio-temporal kriging from IBD mixture modelling for European Farmer-related source groups from (A) British Isles, (B) France/Iberia, (C) Italy and (D) Bronze Age Anatolia. Kriging results from around the time of arrival of Steppe ancestry (4822 BP), the start of the Bell Beaker migrations (4300 BP), the Middle/Late Bronze Age (3314 BP) and the Iron Age (2560 BP) are shown (top to bottom). Kriging was performed on the IBD mixture modelling Set C3 results.

**Extended Data Fig. 3.**
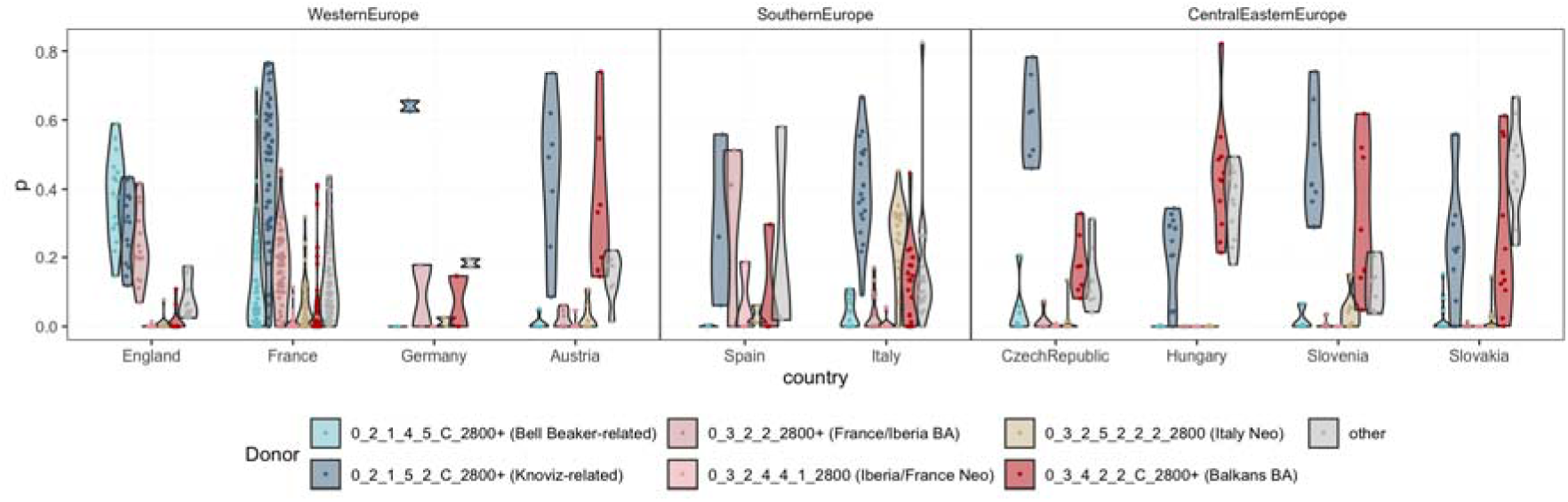
Ancestry proportions from IBD mixture modelling Set C5 for the Early Iron Age (2800–2470 BP), showing the relevant ancestry sources.

**Extended Data Fig. 4.**
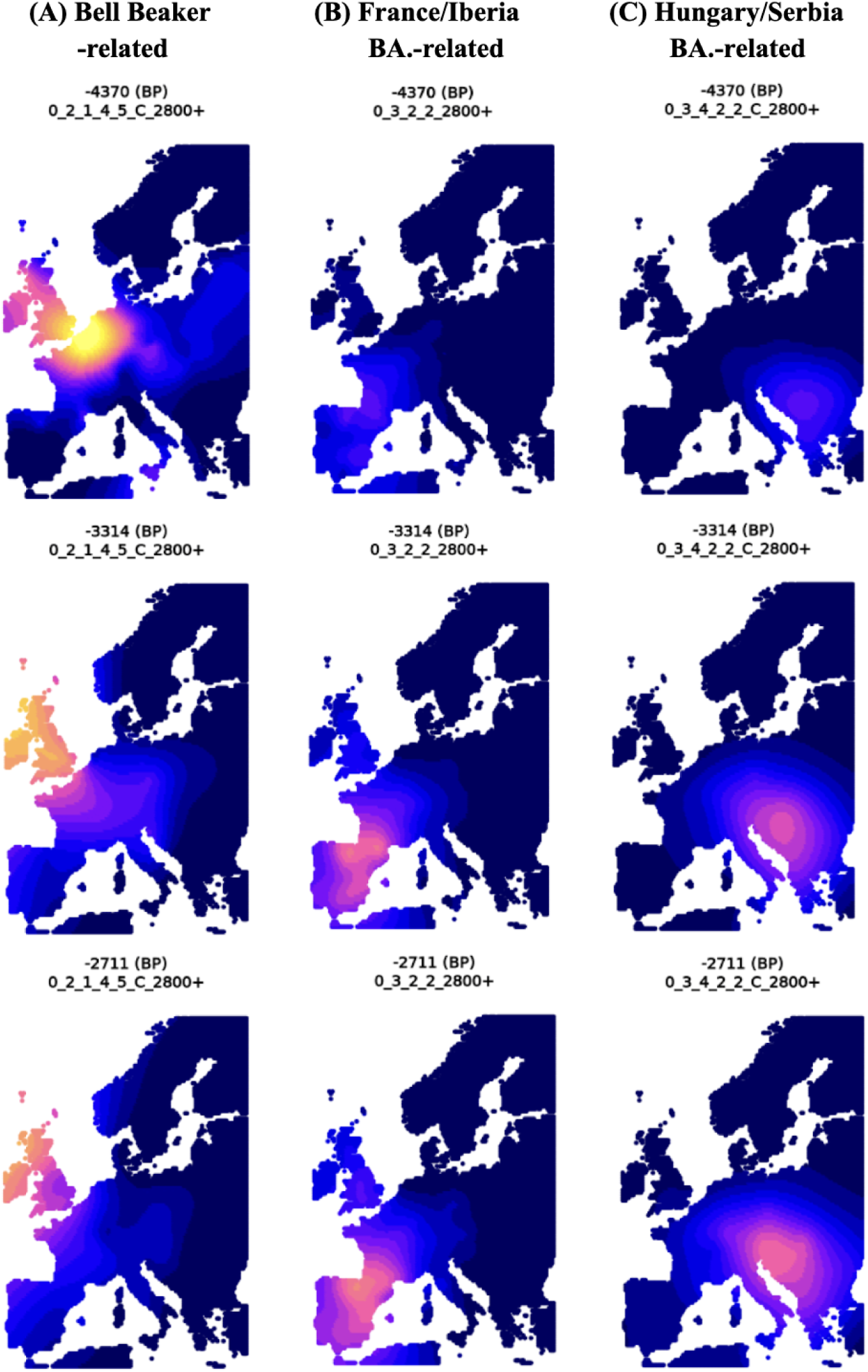
Spatio-temporal kriging results for (A) Bell Beaker-related, (B) Bronze Age France/Iberia-related and (C) Hungary/Serbia-related sources from IBD mixture modelling Set C4.

**Extended Data Fig. 5.**
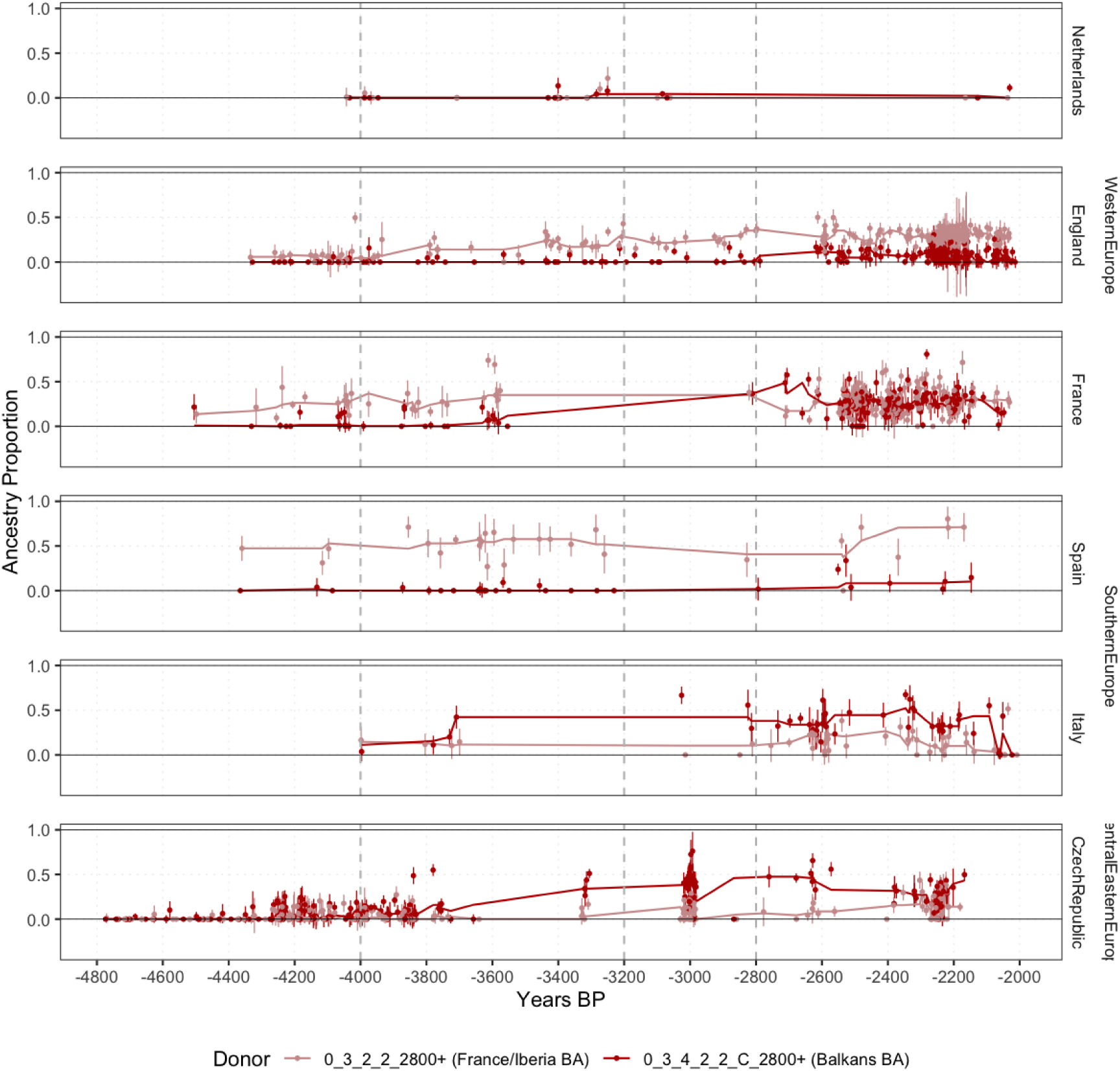
IBD mixture modelling results (Set C4) for the Netherlands, England, France, Spain, Italy and the Czech Republic during the Bronze and Iron Age, highlighting the spread of Southeastern (Hungary/Serbia) and Southwestern (France/Iberia) European Bronze Age ancestry. The trendline shows the rolling median with a window of 5.

**Extended Data Fig. 6.**
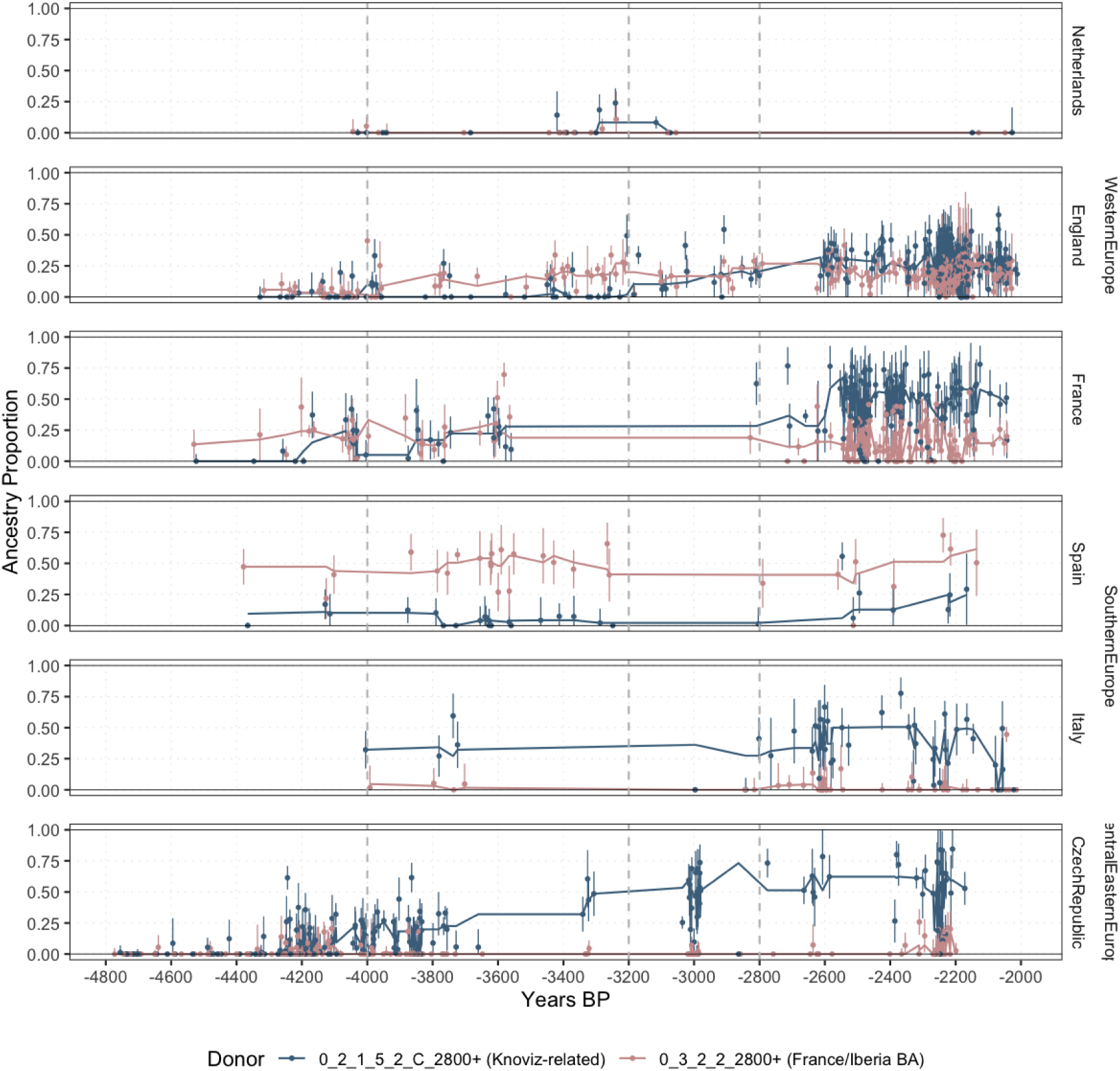
IBD mixture modelling results (Set C5) for the Netherlands, England, France, Spain, Italy and the Czech Republic during the Bronze and Iron Ages, highlighting the spread of Knovíz-related ancestry relative to French/Iberian Bronze Age ancestry. The trendline shows the rolling median with a window of 5.

