## Supplementary figures and images for "Tracing the spread of Celtic languages using ancient genomics"

### Supplementary Fig. S1.6

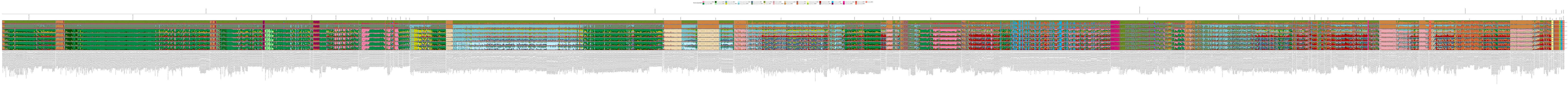

### Supplementary Fig. S1.7

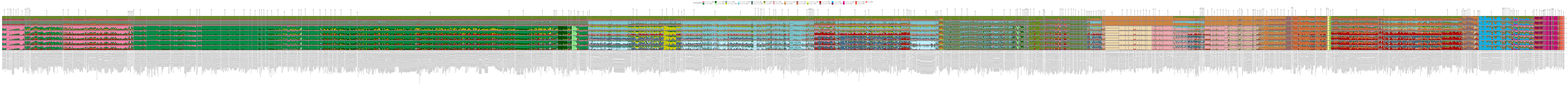
