## Supplementary Notes S1-S6 for "Tracing the spread of Celtic languages using ancient genomics"

#### Supplementary Note S1. IBD Mixture Modelling Results.

Hugh McColl[1,2] and Martin Sikora[1]

[1]Lundbeck Foundation GeoGenetics Center, Globe Institute, University of Copenhagen, Copenhagen, Denmark

[2]Department of Historical Studies, University of Gothenburg, Gothenburg, Sweden

IBD Mixture modelling was undertaken as described in the methods, on the set of 4587 individuals (Supplementary Table S1.1). Additional detail, including comparisons to more traditional methods, reliability of the calling of short IBD segments, the effect of temporal distance between sources and targets, and of the number of source individuals in a group are explored in [41,44]

In this study we use six sets of sources, referred to as Set C1-C7:

Set C1. This set corresponds to set 3 from [41]. This source set highlights the three distinct sources of Farmer ancestry in Bronze Age European - Globular Amphora, European Farmers, and Bronze Age Anatolians. The temporal and spatial position of the source individuals in this cluster can be visualised in Supplementary Fig. S1.1.

Set C2. This set corresponds to set 4 from [41]. This set highlights the distinction between Bell Beaker- and Corded Ware-related ancestry. The temporal and spatial position of the source individuals in this cluster can be visualised in Supplementary Fig. S1.1. and Supplementary Fig. S1.2.

Set C3. This set corresponds to set 5 from [41], but also includes a series of Bronze Age sources from Northern Europe, and here includes Farmer-related sources from the British Isles, France/Iberia and Italy. The temporal and spatial position of the source individuals in this cluster can be visualised in Supplementary Fig. S1.1, Supplementary Fig. S1.2 and Supplementary Fig. S1.3.

Set C4. This set includes the same individuals as Set C3, but also includes representatives of Bronze Age Balkans (Hungary/Serbia) and Bronze Age Southwest Europe (France/Iberia). The temporal and spatial position of the source individuals in this cluster can be visualised in Supplementary Fig. S1.1, Supplementary Fig. S1.2, Supplementary Fig. S1.3 and Supplementary Fig. S1.4.

Set C5. This set includes the same individuals as Set C4, but also includes representatives of the Knovíz Culture. The temporal and spatial position of the source individuals in this cluster can be visualised in Supplementary Fig. S1.1, Supplementary Fig. S1.2, Supplementary Fig. S1.3 and Supplementary Fig. S1.4.

Set C6. This set includes the same individuals as Set C5, but also includes individuals from Bronze Age England (who are used as a ‘left’ population in the qpAdm analyses). The temporal and spatial position of the source individuals in this cluster can be visualised in Supplementary Fig. S1.1, Supplementary Fig. S1.2, Supplementary Fig. S1.3 and Supplementary Fig. S1.4.

The colours used to refer to the source groups for IBD mixture modelling can be seen in Fig. S1.5. The samples included in each cluster are detailed in Supplementary Table S1.2.

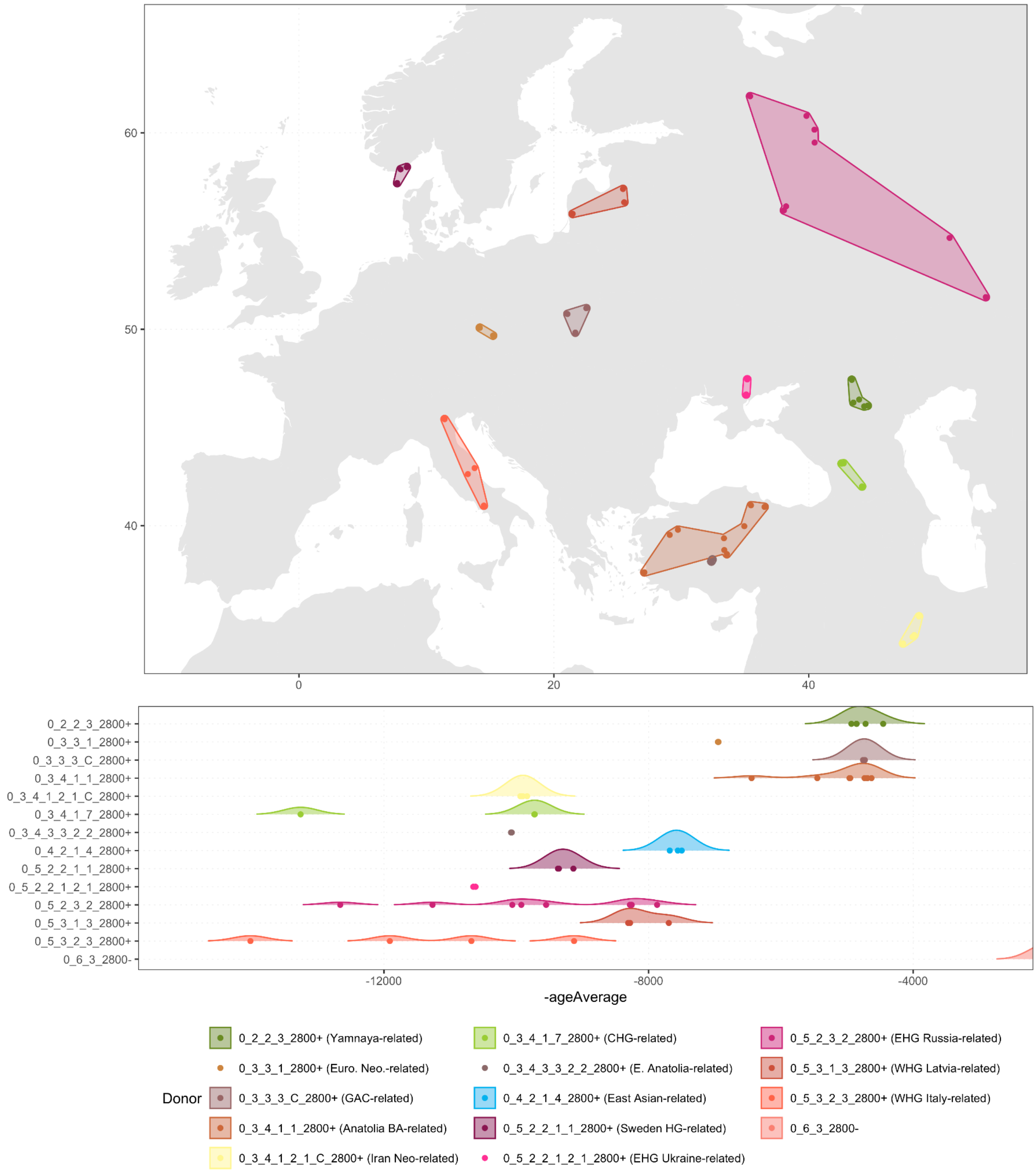

**Supplementary Fig. S1.1. Spatial and temporal distribution of all samples from IBD mixture modelling Source Set C1.** These samples are also included in Sets C2-C6. The East Asian and African sources are not shown.

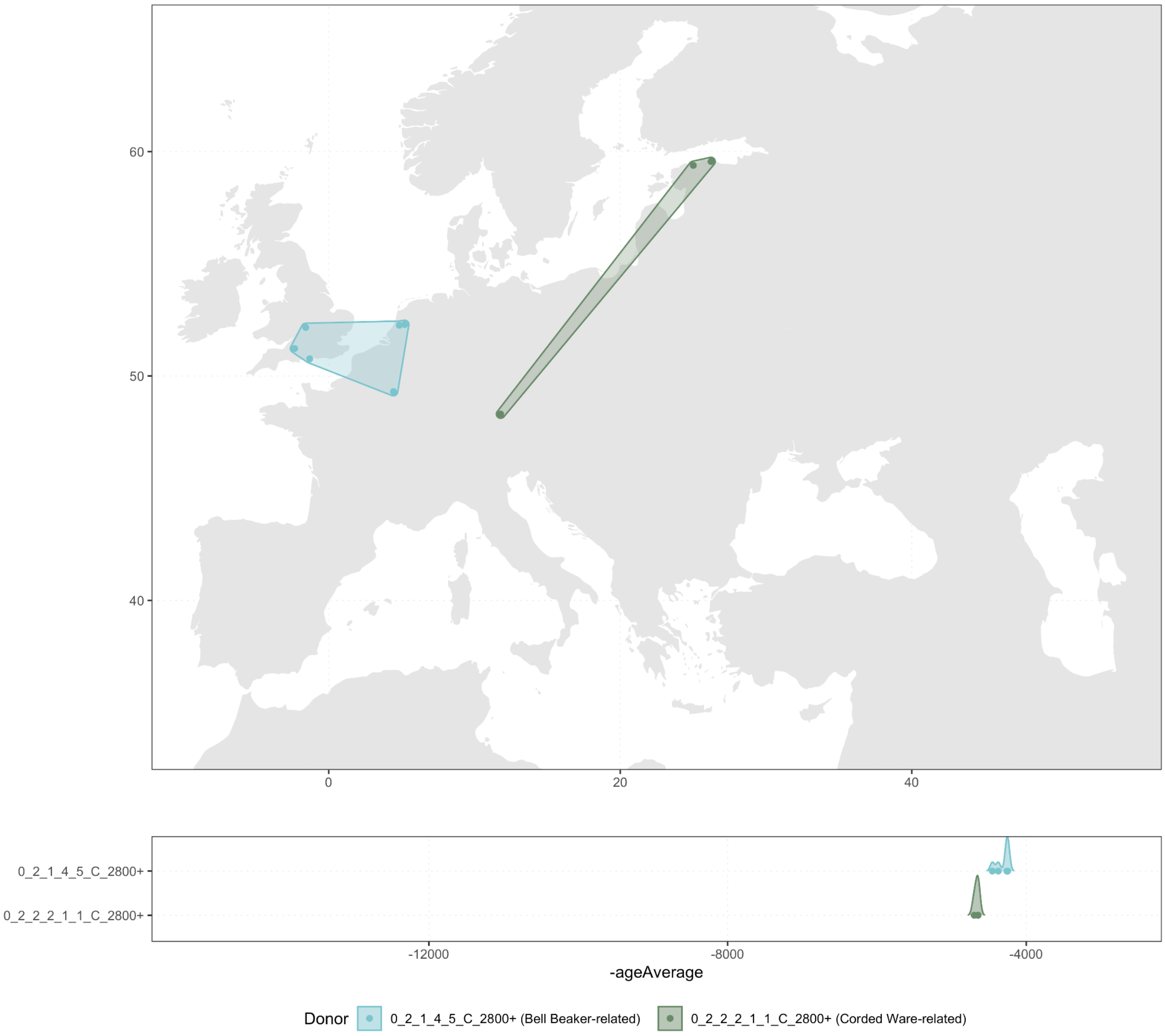


**Supplementary Fig. S1.2. Spatial and temporal distribution of additional samples from IBD mixture modelling Source Set C2.** These samples are also included in Sets C3-C6. Samples from Supplementary Fig. S1.x3 are also included in Set C2.

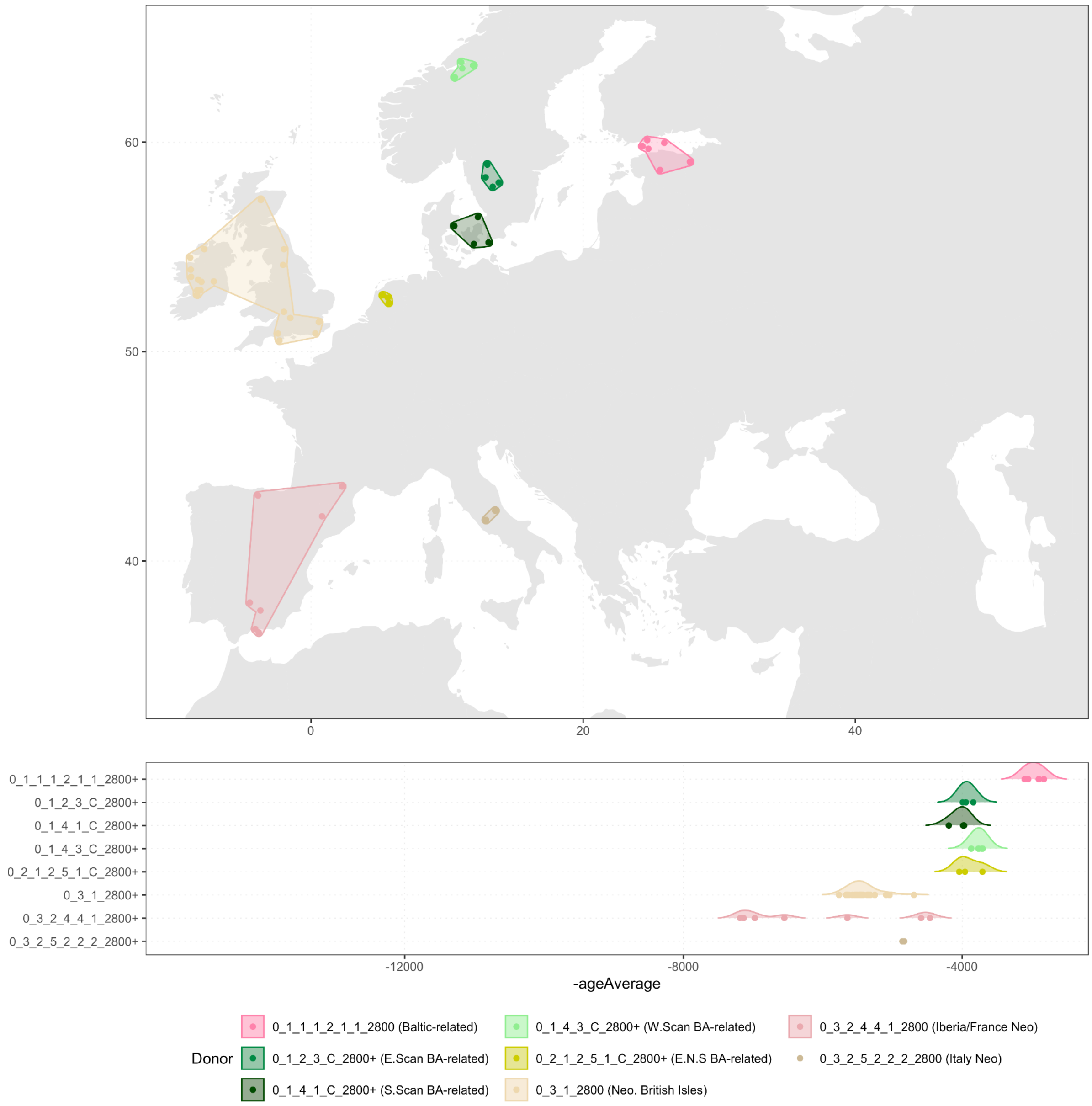


**Supplementary Fig. S1.3. Spatial and temporal distribution of additional samples from IBD mixture modelling Source Set C3.** These samples are also included in Sets C4-C6. Samples from Supplementary Fig. S1.x3 and Supplementary Fig. S1.x4 are also included in Set C3.

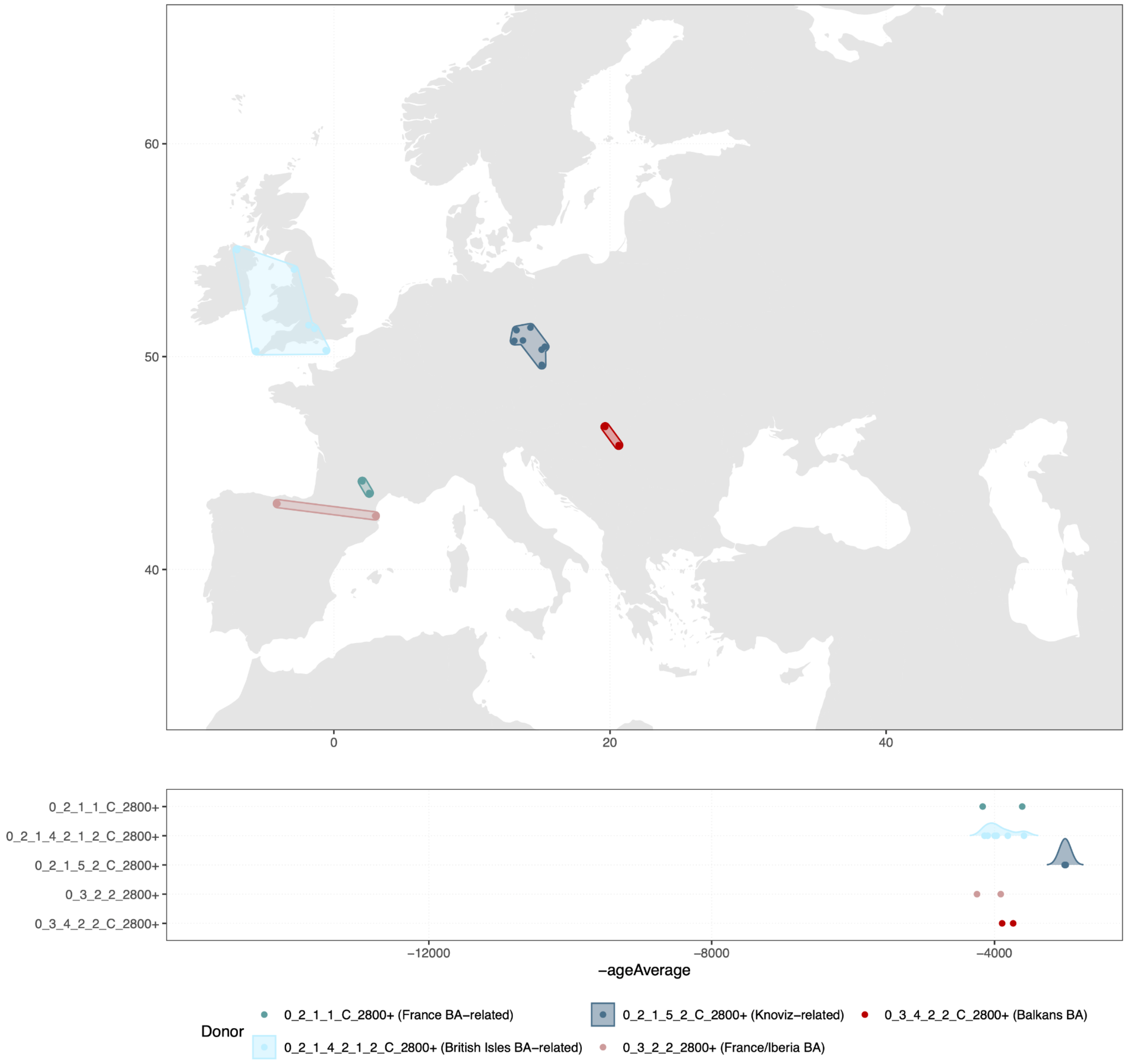


**Supplementary Fig. S1.4. Spatial and temporal distribution of additional samples from IBD mixture modelling Source Sets C4-7.** Samples from Supplementary Fig. S1.x3, Supplementary Fig. S1.x4 and Supplementary Fig. S1.x5 are also included in sets C4, C5 and C6.

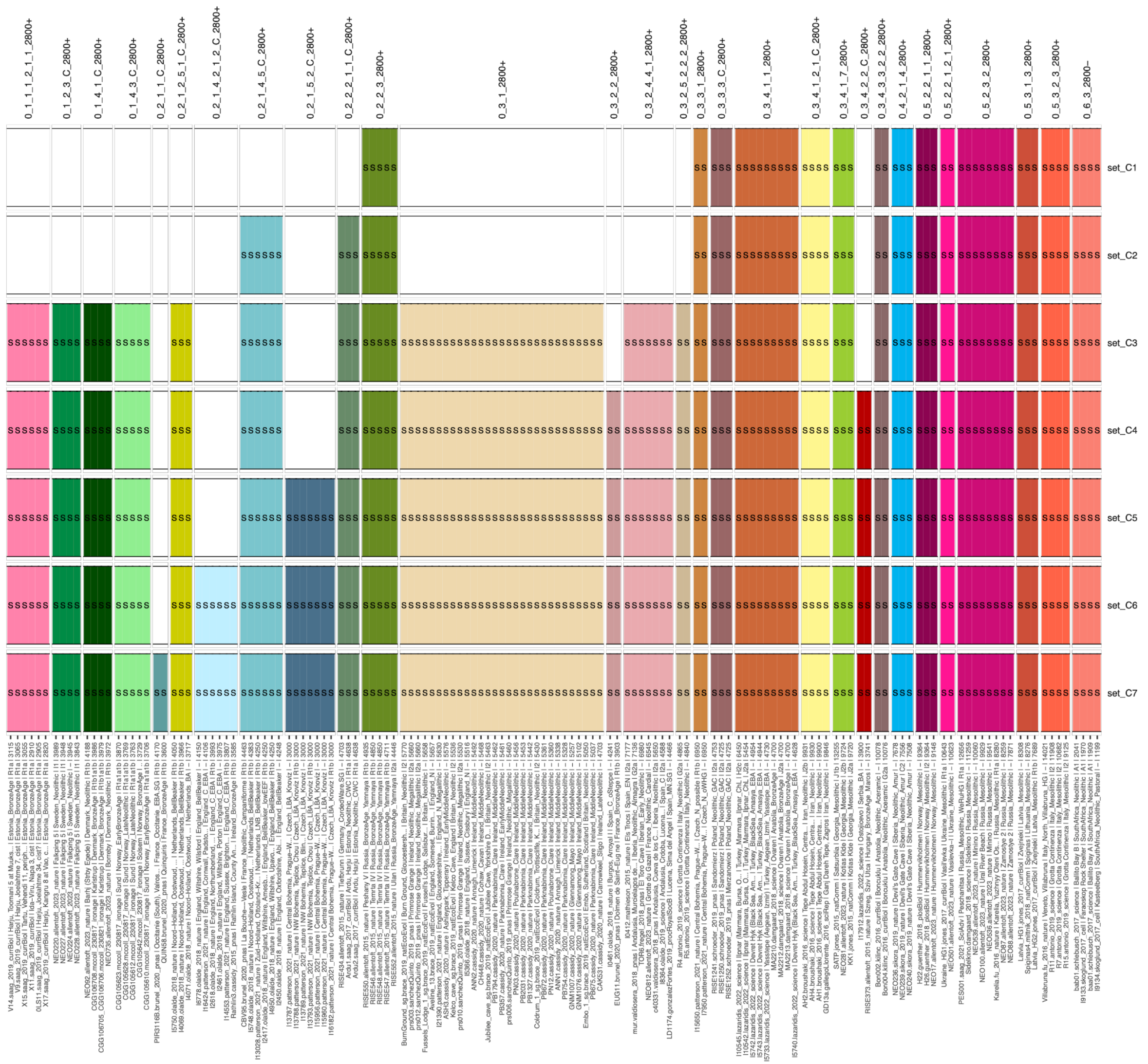


**Fig. S1.5. IBD mixture modelling sources (Sets C1-C6).**

As targets for the IBD mixture modelling, we included all ancient individuals from Europe between 5600 - 600 BP. The full IBD mixture modelling results can be found in Table S1.3, and plotted faceted by country (Fig. S1.6) and by cluster (Fig. S1.7). The sources for each set and associated colours can be found in Fig.1.5.

LARGE FIGURE INCLUDED SEPARATELY

**Fig. S1.6. IBD mixture modelling results (Sets C1-C7) for all European samples from 5600–600 BP, faceted by region and country.**

LARGE FIGURE INCLUDED SEPARATELY

**Fig. S1.7. IBD mixture modelling results (Sets C1-C7) for all European samples from 5600–600 BP, faceted by cluster.**

For the main regions of relevance to this study (The Netherlands, England, Scotland, France, Spain, Italy and the Czech Republic), we plotted the IBD mixture modelling results as violin plots for a series of time transects: Early BA (4800-4000 BP), Middle BA (4000-3200), Late BA (3200-2800 BP), Iron Age (2800-2470 BP), Iron Age (2470-2000 BP). These can be seen in Supplementary Fig. S1.8 (Set C1), Supplementary Fig S1.9 (Set C2), Supplementary Fig S1.10 (Set C3), Supplementary Fig S1.11 (Set C4), Supplementary Fig S1.12 (Set C5), Supplementary Fig. S1.13 (Set C6). Only samples modelled with at least 2% Yamnaya ancestry in Set C1 were included in these plots.

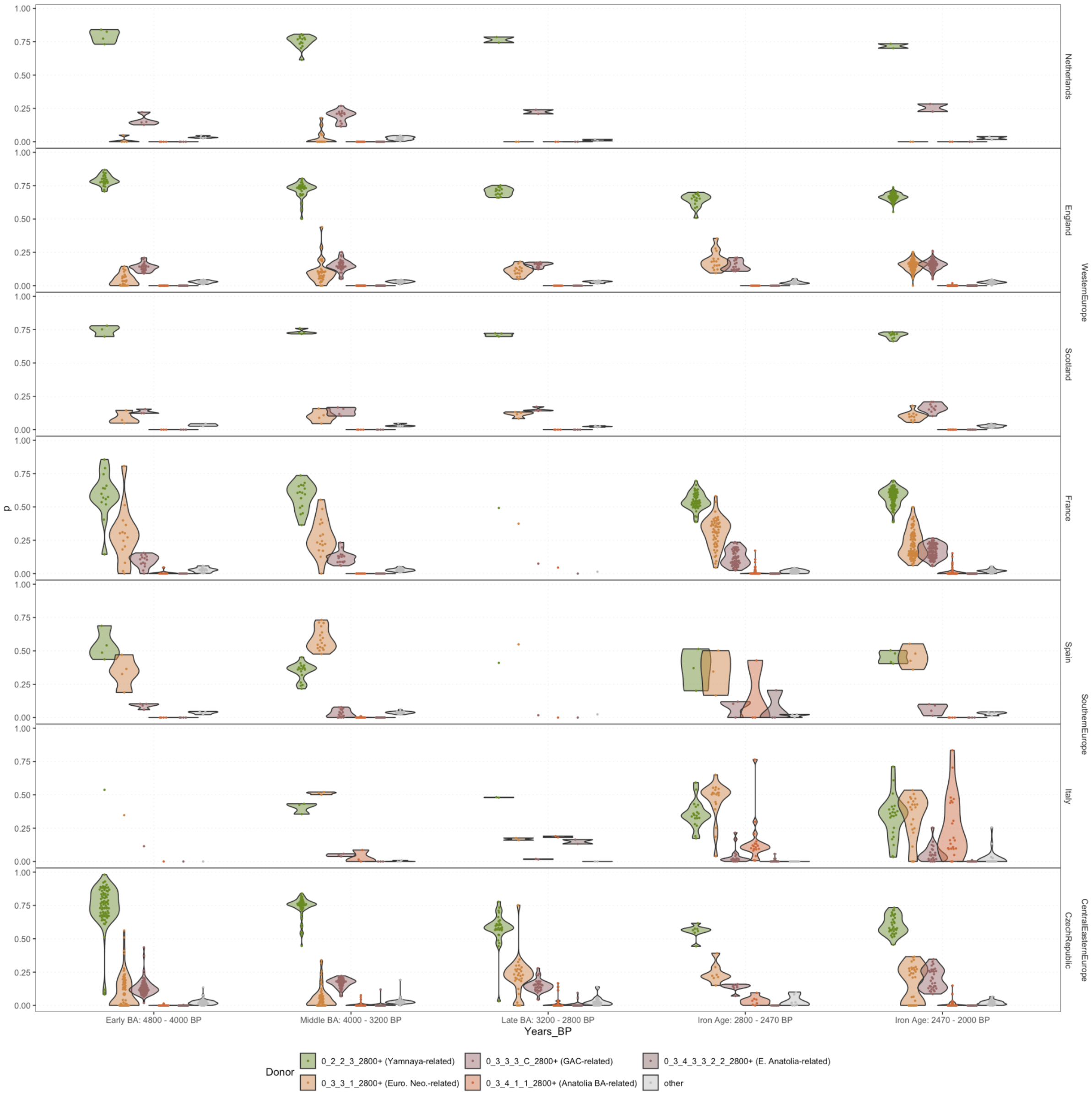


**Supplementary Fig. S1.8. IBD mixture modelling results using Set C1, highlighting the variation in Yamnaya and various Neolithic Farmer-related ancestries.** Only individuals with more than 2% Yamnaya ancestry modelled using Set C1 are shown. The full set of IBD mixture modelling results can be found in Supplementary Fig. S1.1.

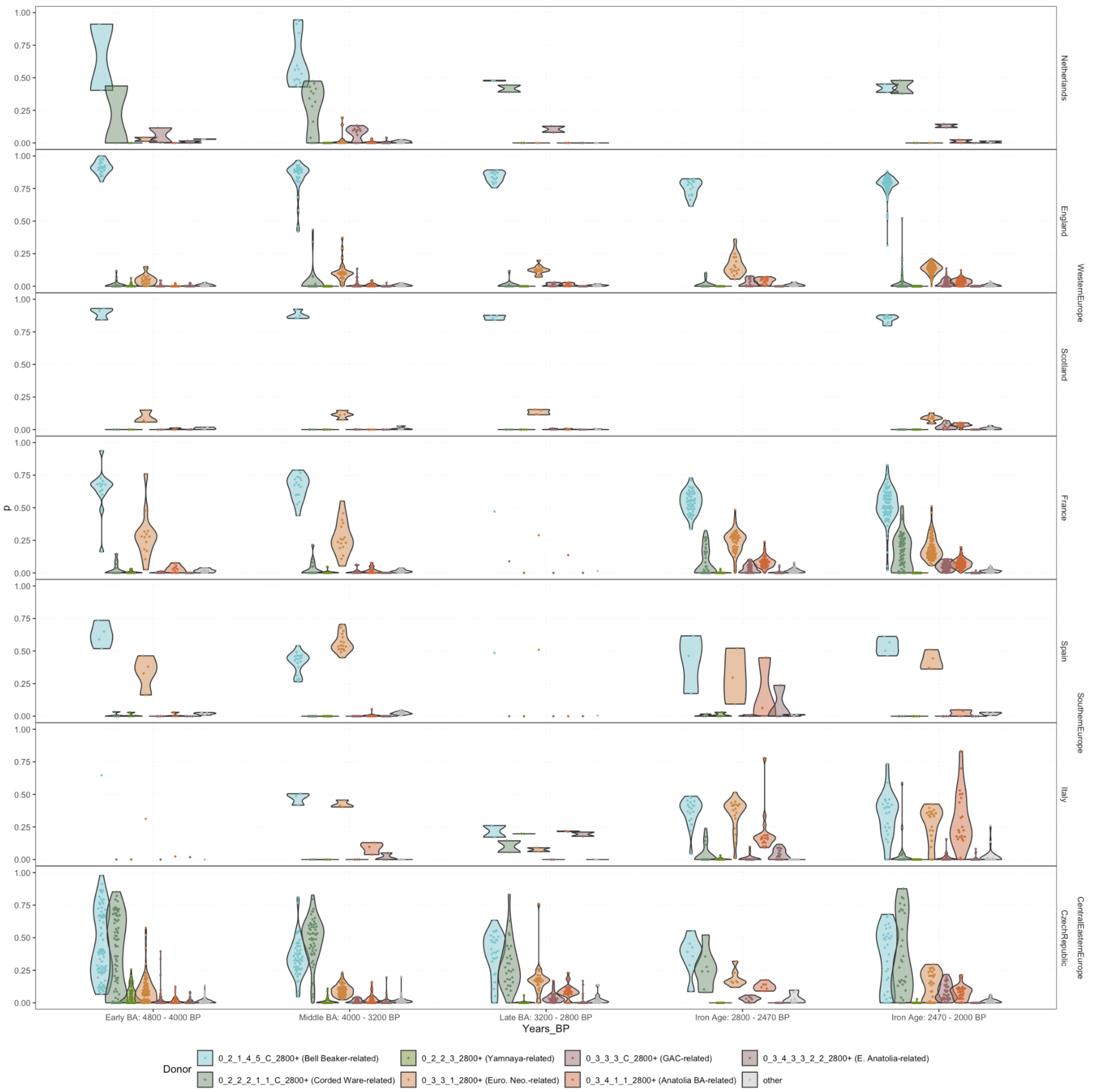


**Supplementary Fig. S1.9. IBD mixture modelling results using Set C2, highlighting the variation in Corded Ware and Bell Beaker-related ancestries.** Only individuals with more than 2% Yamnaya ancestry modelled using Set C1 are shown. The full set of IBD mixture modelling results can be found in Supplementary Fig. S1.1.

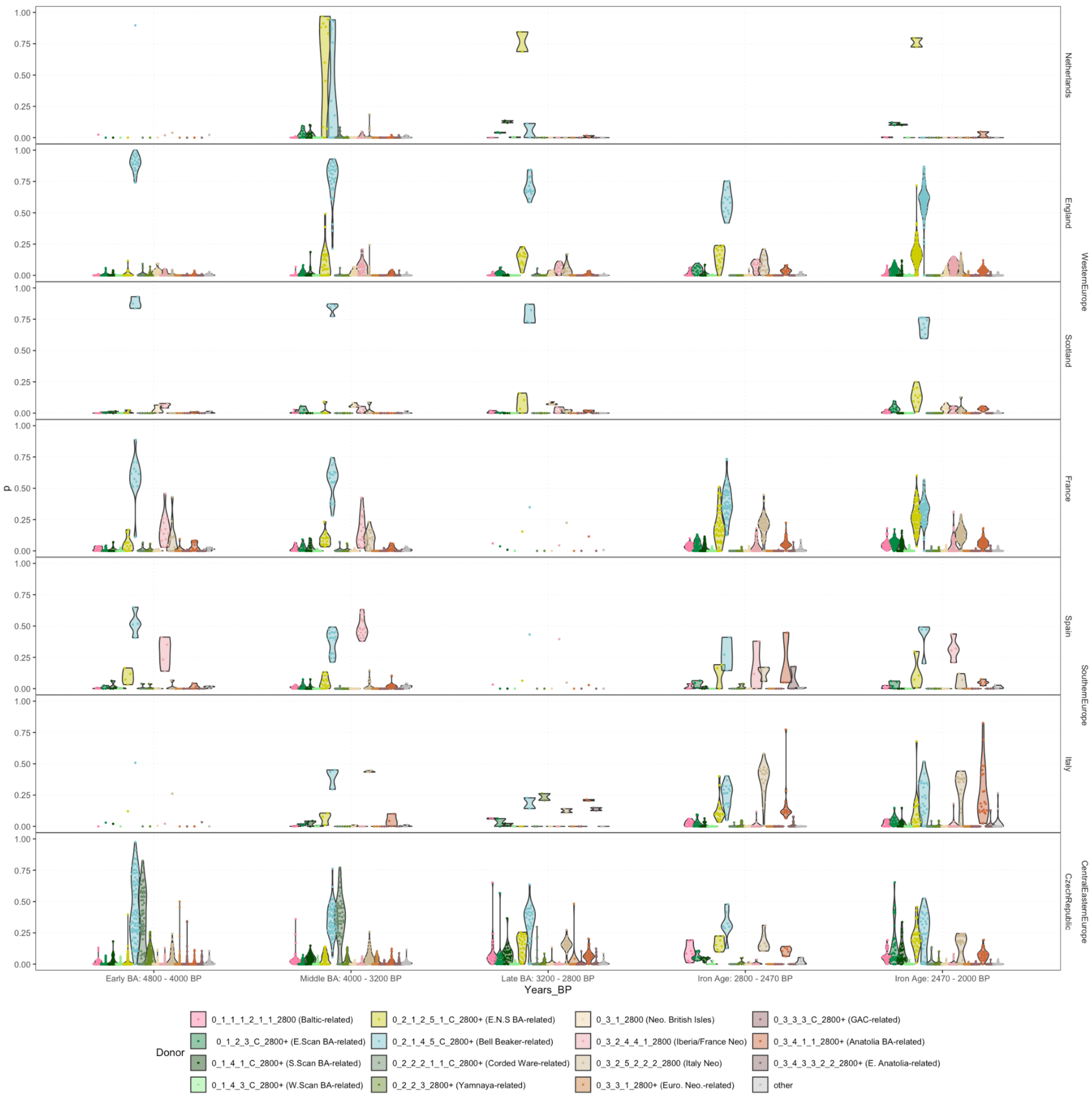


**Supplementary Fig. S1.10. IBD mixture modelling results using Set C3, highlighting the variation in Neolithic Farmer-related ancestries from the British-Irish Isles, Ibera/France and Italy.** Only individuals with more than 2% Yamnaya ancestry modelled using Set C1 are shown. The full set of IBD mixture modelling results can be found in Supplementary Fig. S1.1.

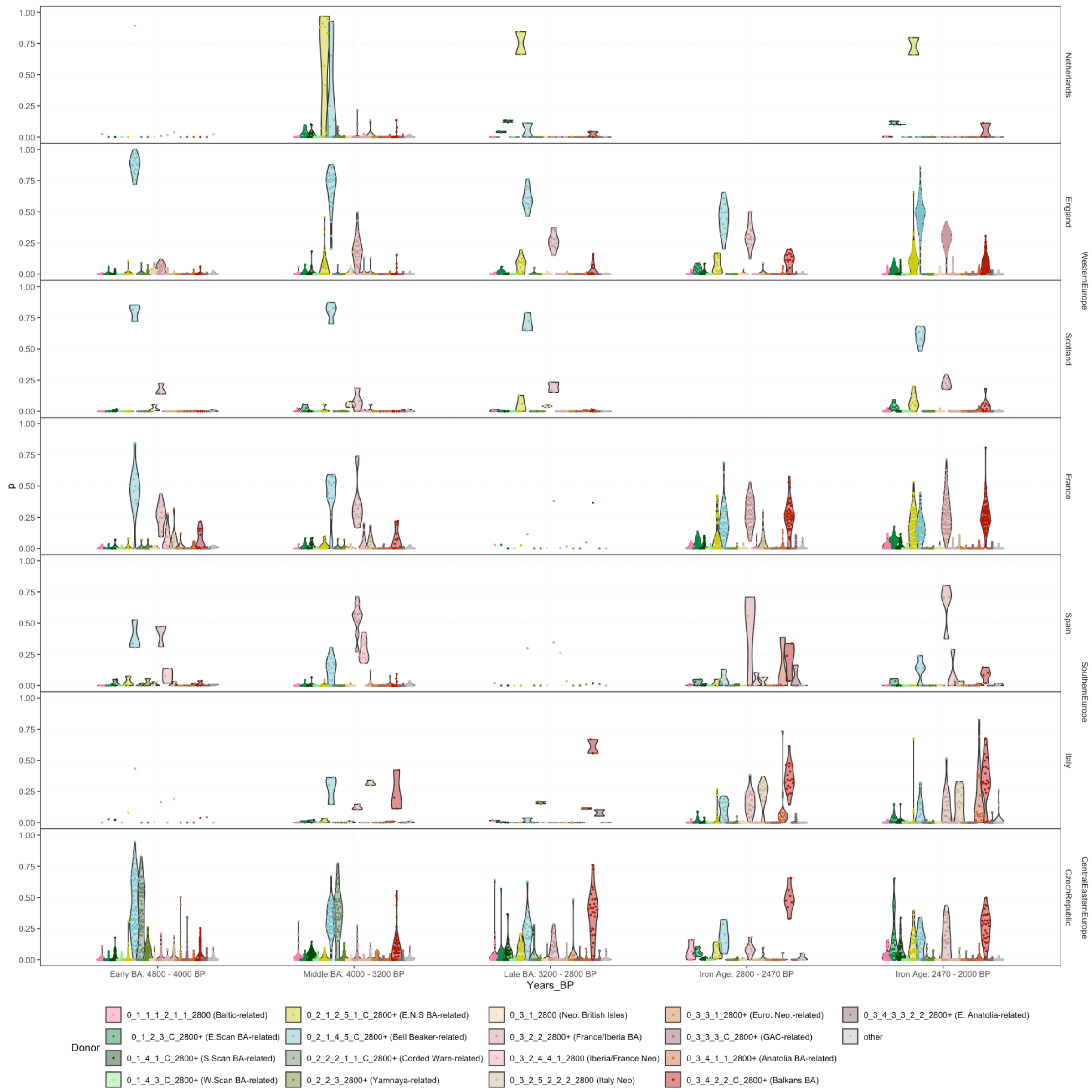


**Supplementary Fig. S1.11. IBD mixture modelling results using Set C4, highlighting the variation in Bronze Age ancestry from the Balkans (Hungary/Serbia) and Southwest Europe (France/Iberia).** Only individuals with more than 2% Yamnaya ancestry modelled using Set C1 are shown. The full set of IBD mixture modelling results can be found in Supplementary Fig. S1.1.

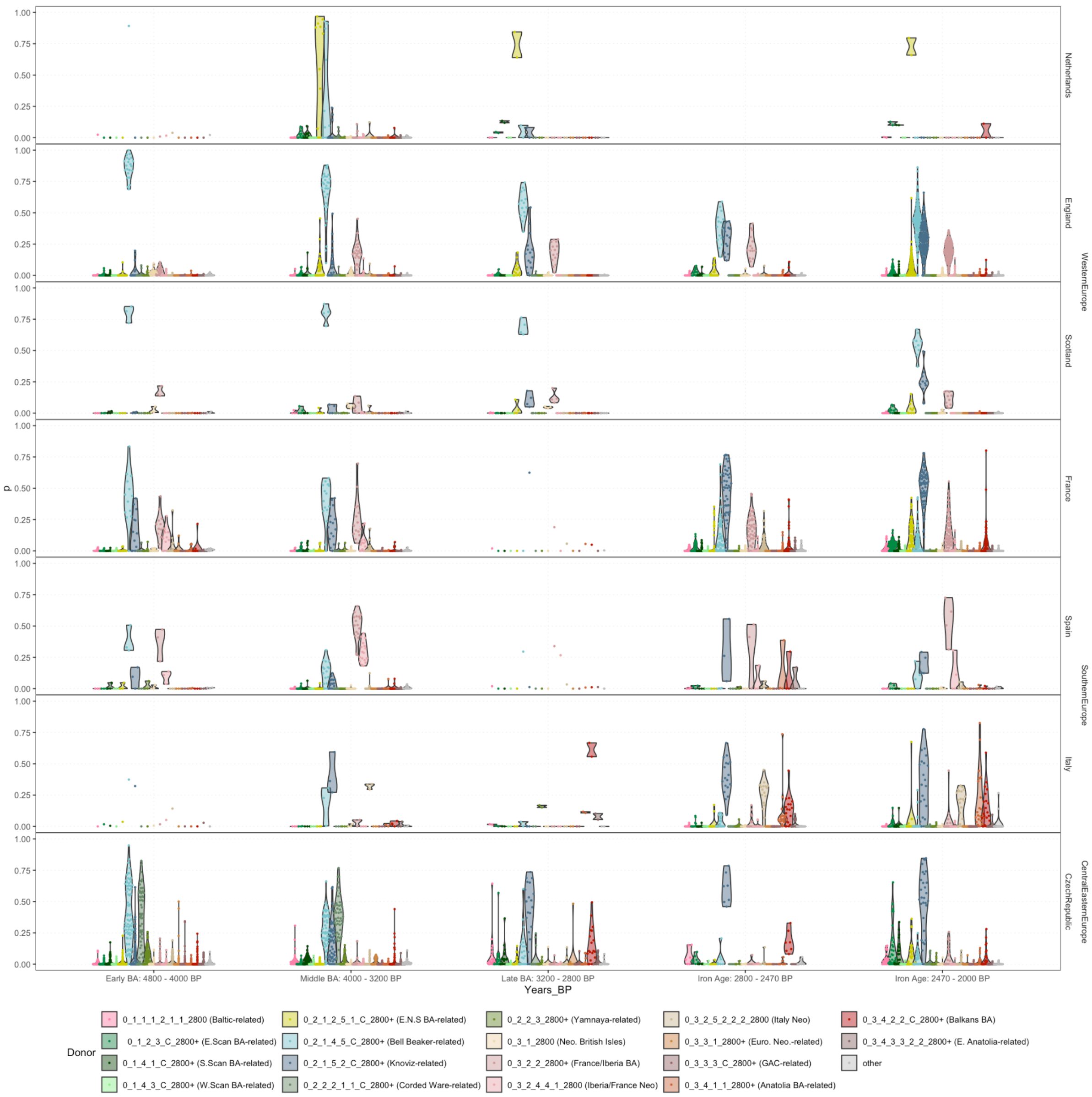


**Supplementary Fig. S1.12. IBD mixture modelling results using Set C5, highlighting the Knovíz-related ancestry.** Only individuals with more than 2% Yamnaya ancestry modelled using Set C1 are shown. The full set of IBD mixture modelling results can be found in Supplementary Fig. S1.1.

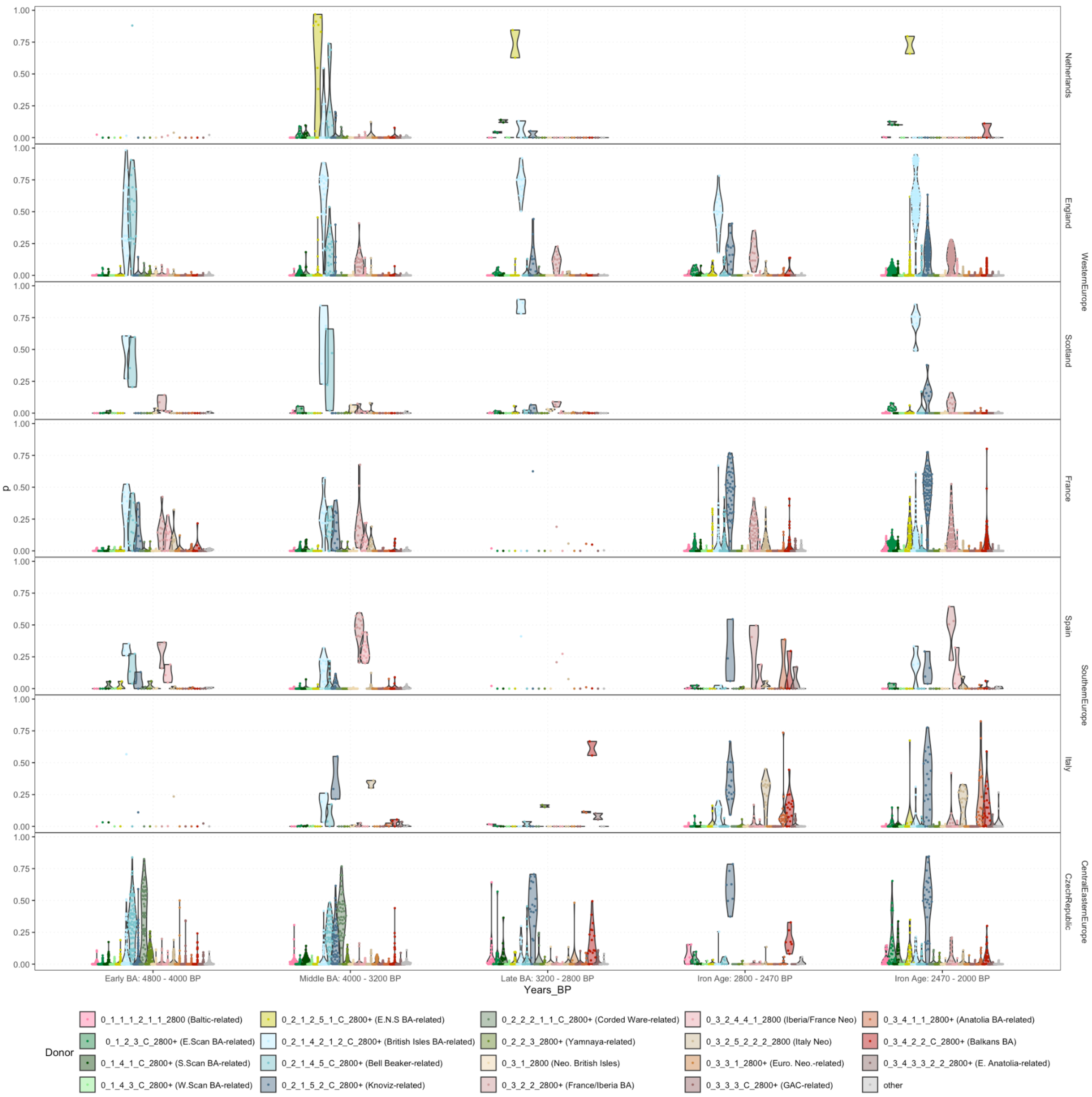


**Supplementary Fig. S1.13. IBD mixture modelling results using Set C6, highlighting the British Isles-related ancestry used in the qpAdm analyses.** Only individuals with more than 2% Yamnaya ancestry modelled using Set C1 are shown. The full set of IBD mixture modelling results can be found in Supplementary Fig. S1.1.

For key sources from sets C2, C3 and C4, we performed spatio-temporal kriging[75] on the IBD mixture modelling, as described in methods, using the 'Europe-Wide' parameters from [41]. The results are shown in Figs. S1.14-S1.23.

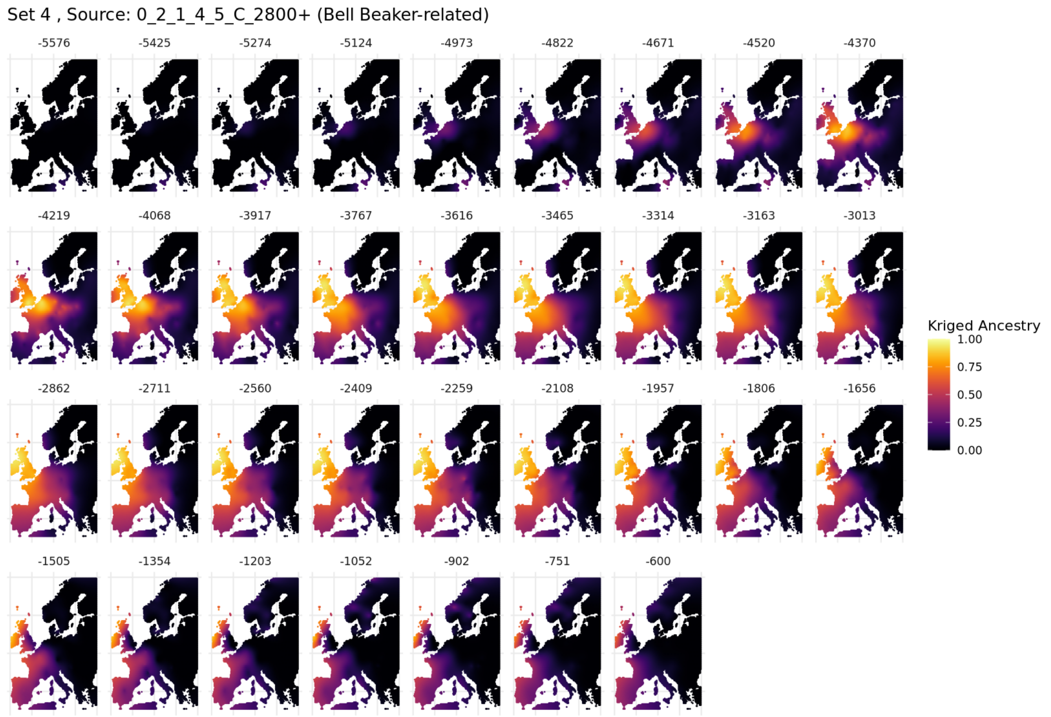


**Supplementary Fig. S1.14. Spatiotemporal kriging results for the Bell Beaker-related source from Set C2 using 'Europe-Wide' parameters.**

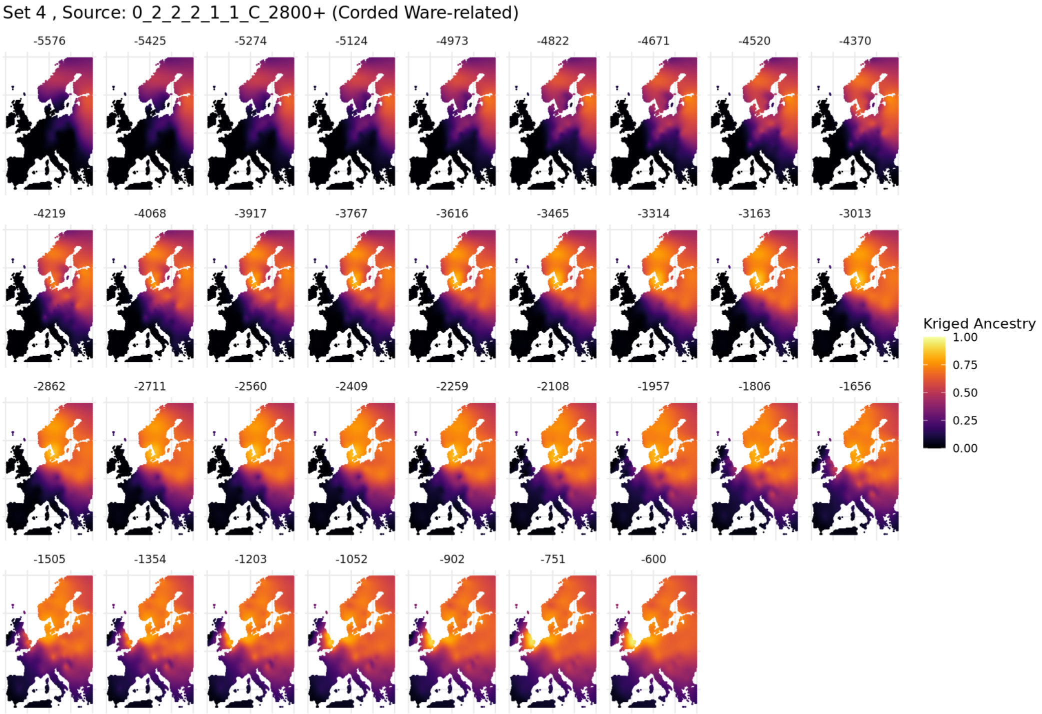


**Supplementary Fig. S1.15. Spatiotemporal kriging results for the Corded Ware-related source from Set C2 using 'Europe-Wide' parameters.**

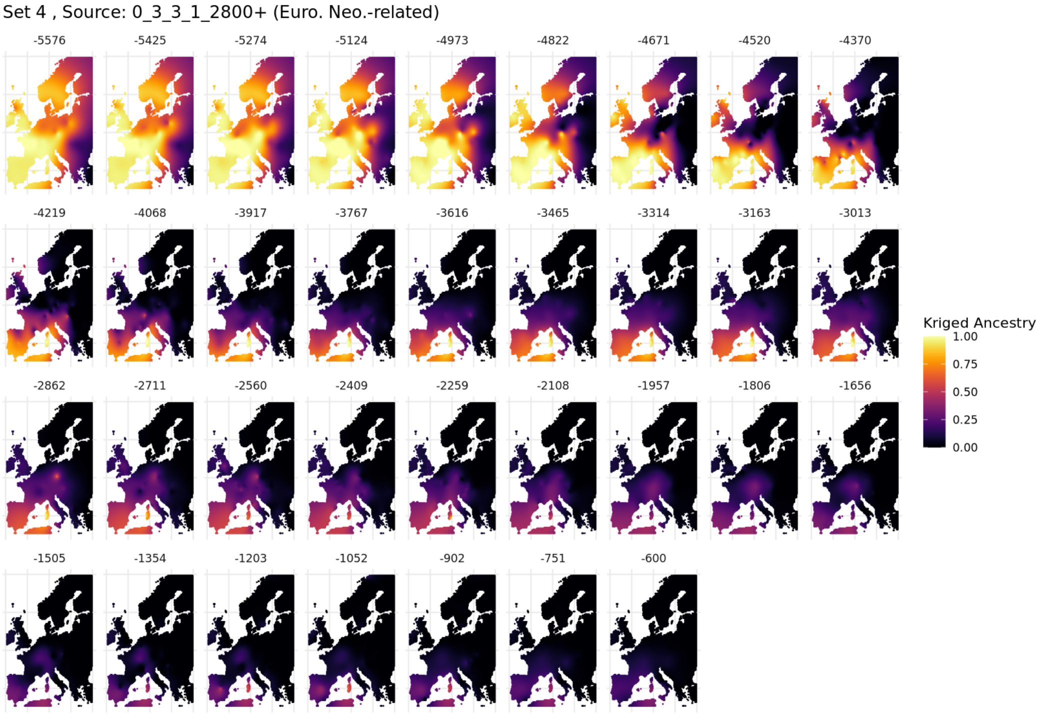


**Supplementary Fig. S1.16. Spatiotemporal kriging results for the European Farmer-related source from Set C2 using 'Europe-Wide' parameters.**

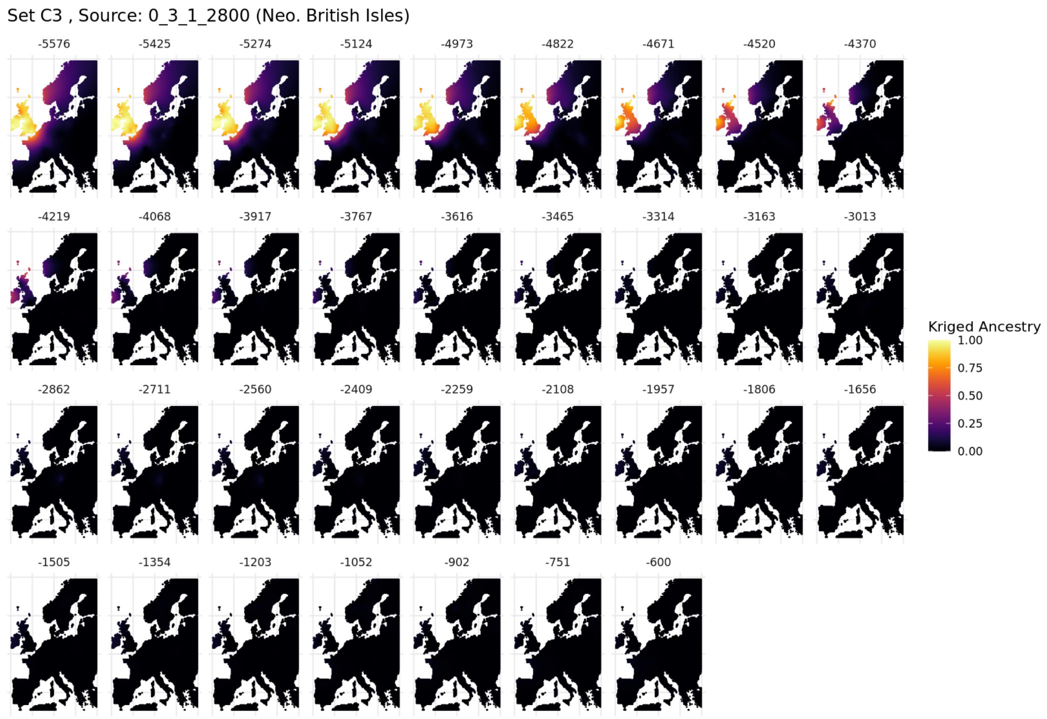


**Supplementary Fig. S1.17. Spatiotemporal kriging results for the British/Irish Neolithic Farmer-related source from Set C3 using 'Europe-Wide' parameters.**

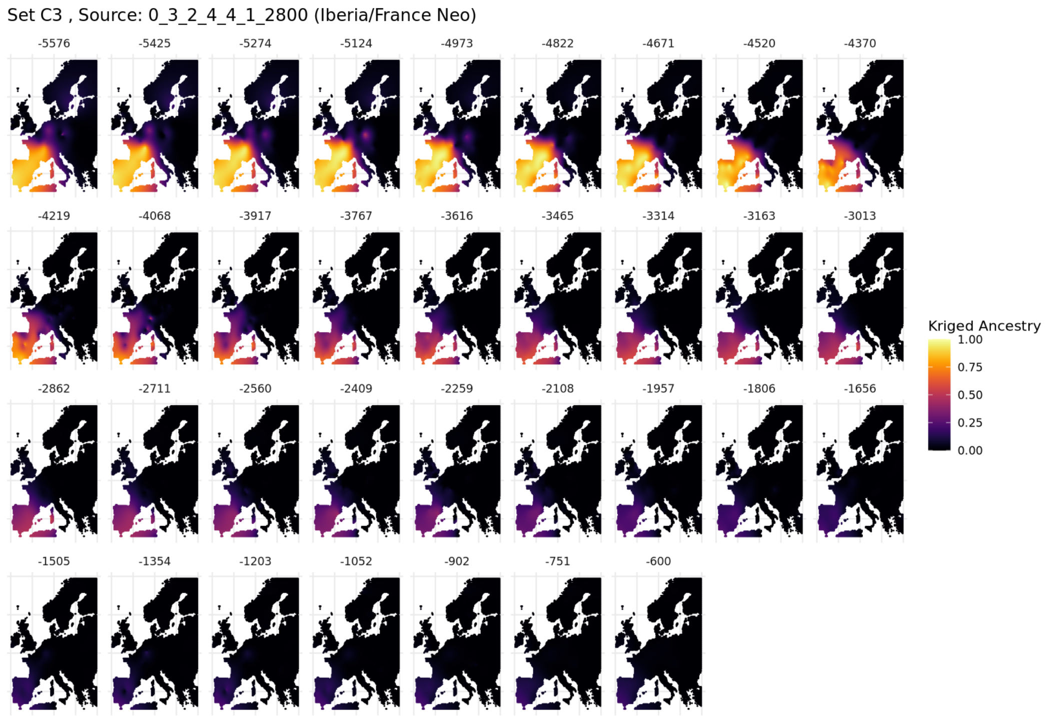


**Supplementary Fig. S1.18. Spatiotemporal kriging results for the France/Iberia Neolithic-related source from Set C3 using 'Europe-Wide' parameters.**

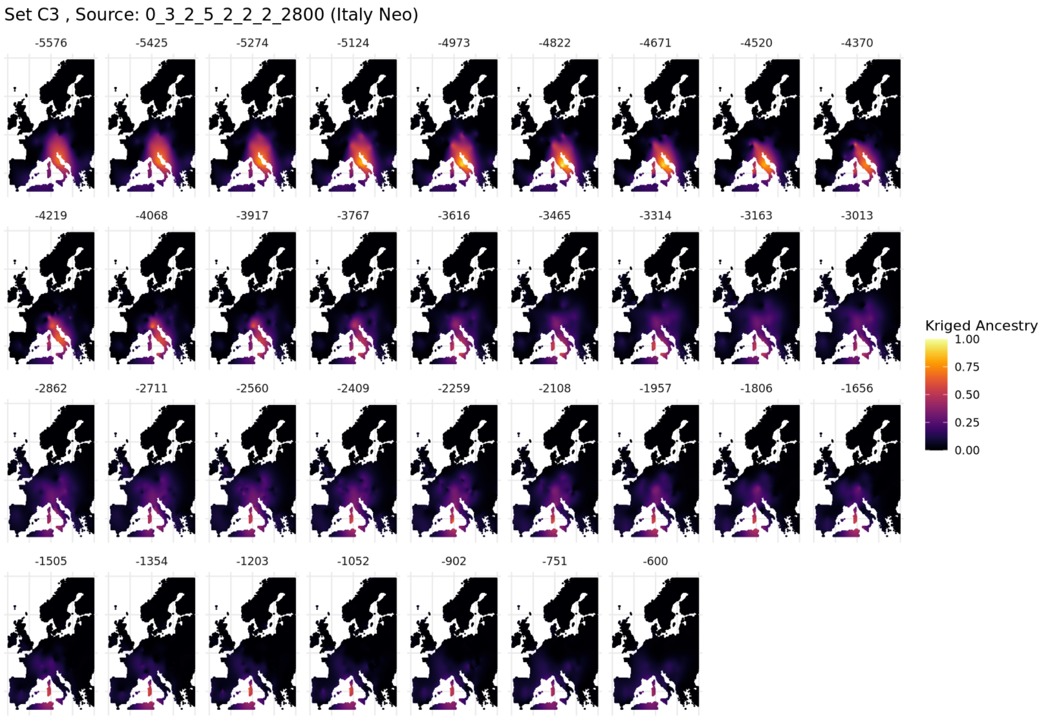


**Supplementary Fig. S1.19. Spatiotemporal kriging results for the Italian Neolithic-related source from Set C3 using 'Europe-Wide' parameters.**

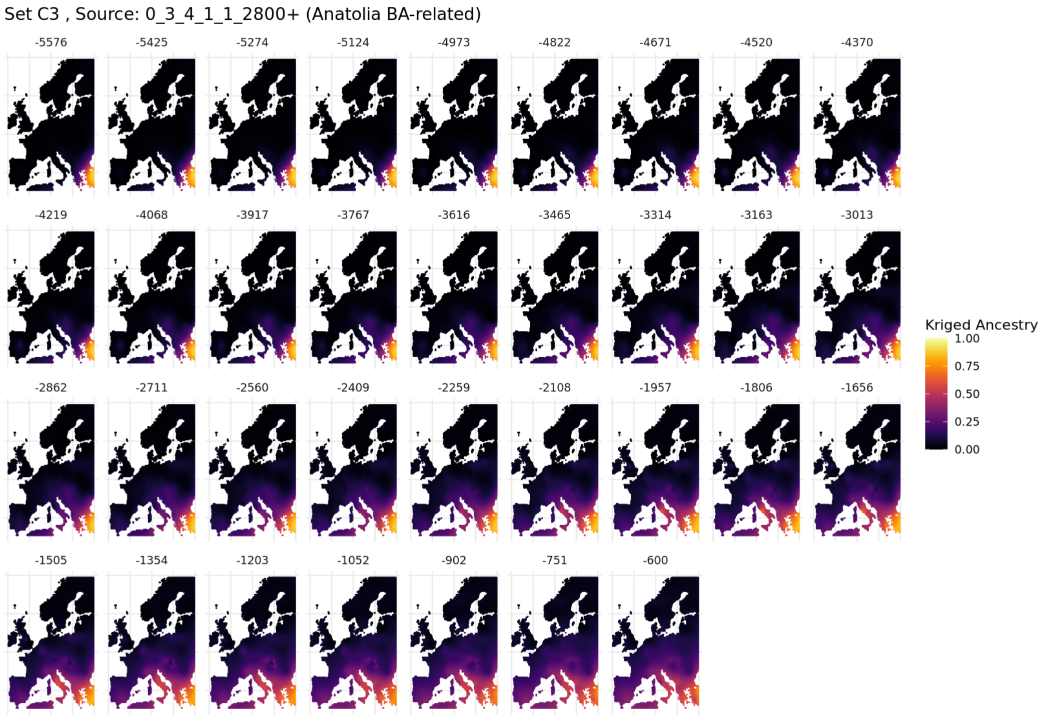


**Supplementary Fig. S1.20. Spatiotemporal kriging results for the Anatolian BA-related source from Set C3 using 'Europe-Wide' parameters.**

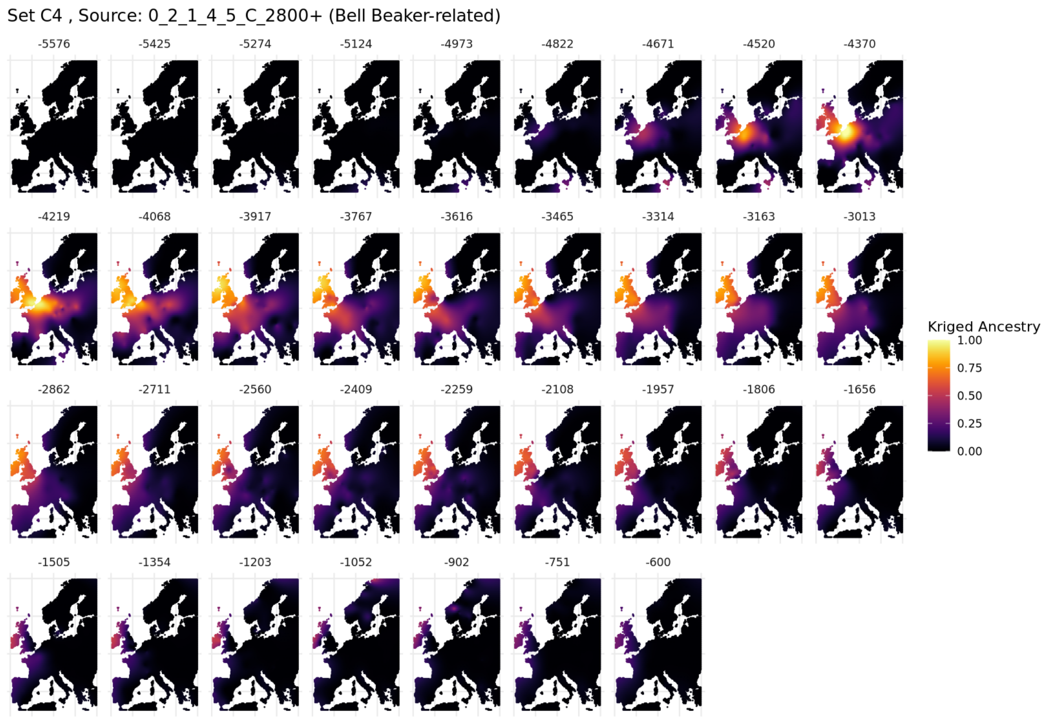


**Supplementary Fig. S1.21. Spatiotemporal kriging results for the Bell Beaker-related source from Set C4 using 'Europe-Wide' parameters.**

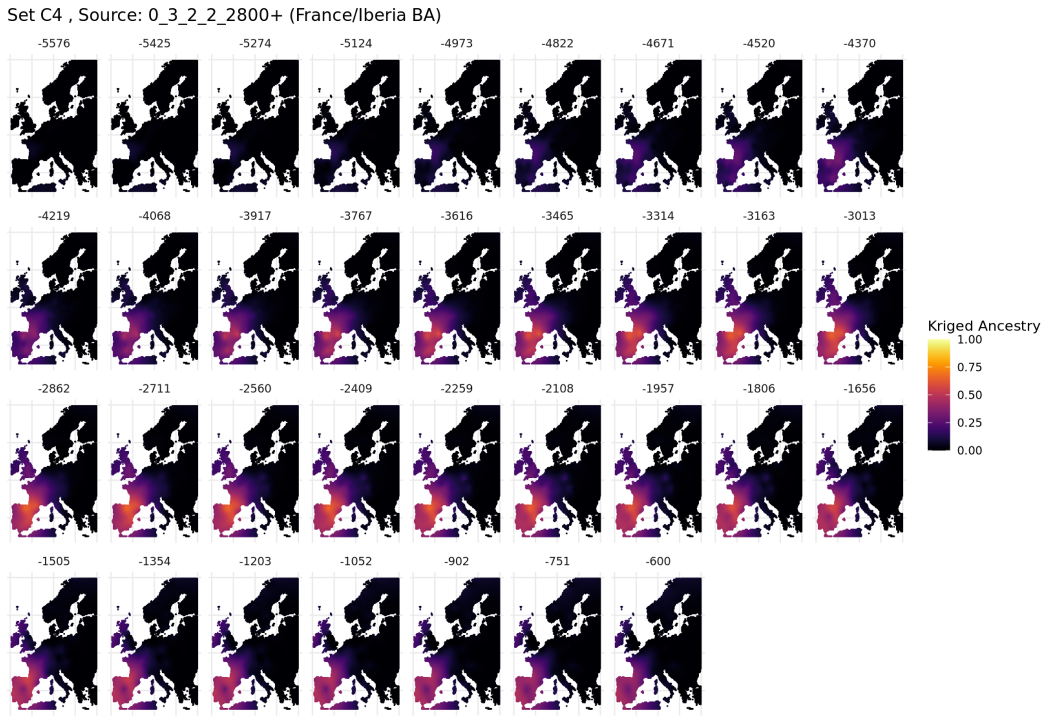


**Supplementary Fig. S1.22. Spatiotemporal kriging results for the France/Iberia BA-related source from Set C4 using 'Europe-Wide' parameters.**

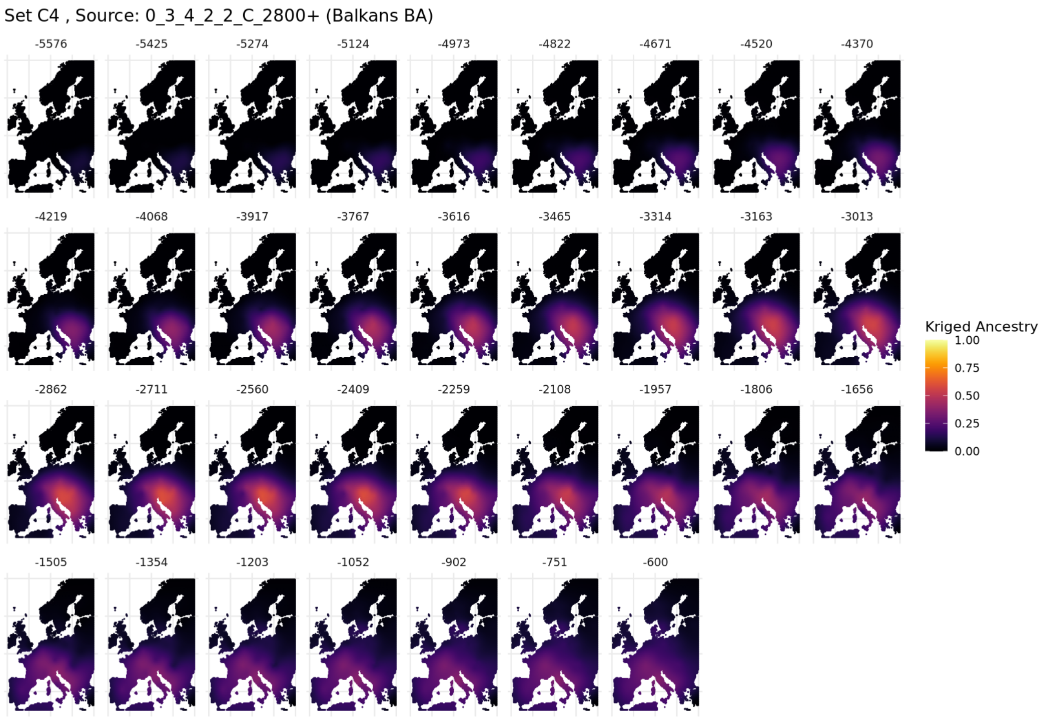


**Supplementary Fig. S1.23. Spatiotemporal kriging results for the Hungary/Serbia-related source from Set C4 using 'Europe-Wide' parameters.**

To incorporate findings of recent papers focussing on Celtic migrations, we also prepared an auxiliary dataset, including samples from [19,78,79]. We imputed, merged, and called IBD segments in the same way as the main dataset. We used the clustering from the main dataset, and to incorporate these individuals in the mixture modelling results, we treated each individual in the same way as clusters with single individuals are treated: their palette of the single individual represents of the sharing with all other clusters in the dataset, but the single-individual-clusters does not contribute to the palettes of all other individuals.

The additional individuals included in the auxiliary dataset can be found in Supplementary Table S1.4. The mixture modelling results for these individuals can be found in

Supplementary Table S1.5. Ancestry proportions for relevant individuals and source sets are plotted and discussed below.

#### Extended Discussion

*Bell Beaker and Corded Ware-related interactions*

From the IBD mixture modelling results using Set C2, we find early evidence of Bell Beaker-related ancestry being present in France by 4463 BP, the Netherlands by 4384 BP, on the British-Irish Isles by ~4300 BP (England: 4333 BP, Scotland: 4312 BP, Ireland: 3906), and reaching southern Europe by ~4300 BP (Spain 4367 BP, Portugal 4158 BP, Italy 4212 BP). Despite occurring at a similar time, the proportion of Bell Beaker-related ancestry in France, Spain and Italy is less than that of Britain and the Netherlands (Supplementary Fig. S1.9). Further east, the Bell Beaker period was preceded by the Corded Ware Culture. This transition is mirrored genetically in which the Bell Beaker-related individuals follow, or overlap with the end of, the temporal distribution of Corded Ware-related individuals. This is seen in Germany (CWC-related 4703–4358 BP, BB-related by 4439), Hungary (BB-related by 4299 BP, other CWC-related from the same time), Czech Republic (CWC-related 4739–4477 BP, BB-related by 4333), and Poland (CWC-related 4705–4239 BP, BB-related by 4250 BP). Additionally, in all regions, we found a strong temporal correlation between the arrival of Beaker-related ancestry and Y-chromosome haplogroup R-P312, confirming their association (Supplementary Note S4).

Throughout the Bronze and Iron Age, multiple eastward and westward migrations are evident at the peripheries of the region of Bell Beaker-related ancestry. This can be seen, for example, in the Czech Republic, with the expansion of Bell Beaker-related ancestry evident during the Bell Beaker period replacing Corded Ware-related ancestry, and the subsequent reduction during the Únětice period with the reappearance of Corded Ware-related ancestry from the east (Extended Data Fig. 1).

*Urnfield and Celtic-related ancestries in Scandinavia*

Using Set C6 provides the resolution to distinguish between ancestry within these Celtic speaking regions and is also informative in relation to Migration Period movements. Some of the Migration Period individuals from Britain carry high proportions of Knovíz-related ancestry and little to no British-Irish Bronze Age-related, suggestive of migrations from or admixture on the continent (Supplementary Fig. S1.6). Migration Period migrations into Denmark and Sweden detected elsewhere revealed that people carrying some continental ancestry[2,80], but primarily of Scandinavian ancestry[41], arrived in Denmark and Southern Sweden by the Viking Period; here we provide further insight into the source of the continental ancestry: the influx of small proportions of continental ancestry is modelled as Knovíz and Hungary/Serbia Bronze Age-related ancestry, generally lacking British-Irish Bronze Age and French/Iberian Bronze Age. As such, we can exclude the Netherlands, France, Britain and Ireland as a source of this continental ancestry, and infer a source region further east. This stands in direct contrast to Norway, where high proportions of the British-Irish Bronze Age-related ancestry are detected in most individuals with non-local ancestry, consistent with previous studies[80].

*The Linguistic Landscape of Iron Age Iberia*

The finding of the multiple Bell-Beaker-related migrations impacting Iberia further provides a framework for understanding the linguistic landscape of Iron Age Iberia. During the Iron Age, Lusitanian, an Indo-European language related to Italic and Celtic (Supplementary Note

S5), was found alongside the Celtic Celtiberian language. It remains debated whether these two languages diverged over thousands of years within Iberia, or whether the Lusitanian arrived first and Celtiberian later[81–83]. Our results reveal that two Steppe-derived migrations impacted Iberia, the first occurring between 4400–4000 BP and consistent with the arrival of Lusitanian, and the latter between 3200–2500 BP, introducing Celtic and the Urnfield Culture. Thus, in the face of the Celtic expansions, Lusitanian can be understood as a vestige of the first wave of Indo-Europeanization that persisted in Iberia until the second wave introducing Celtic.

*Comparison to previous studies*

To gain additional resolution to distinguish the British Isles from continental populations, we included a Bronze Age source from Britain and Ireland (Set C6), corresponding to the fixed 'left' population described in the qpAdm analyses below. In England from the Late Bronze Age until the Migration Period, individuals tend to be modelled primarily by the British-Irish Bronze Age ancestry, with some Knovíz or French/Iberian Bronze Age-related ancestry (Supplementary Fig. S1.12). In France, Iron Age individuals tend to be modelled primarily by the Knovíz and French/Iberian Bronze Age-related ancestries, but with little to no British-Irish Bronze Age-related ancestry (Supplementary Fig. S1.24). Notable outliers with high British-Irish Bronze Age ancestry in France include two individuals from the Urville-Nacqueville necropolis in Normandy, a site archaeologically linked with Southern England[49], one of whom carries a typical British-Irish Beaker paternal lineage (R1b-L21).

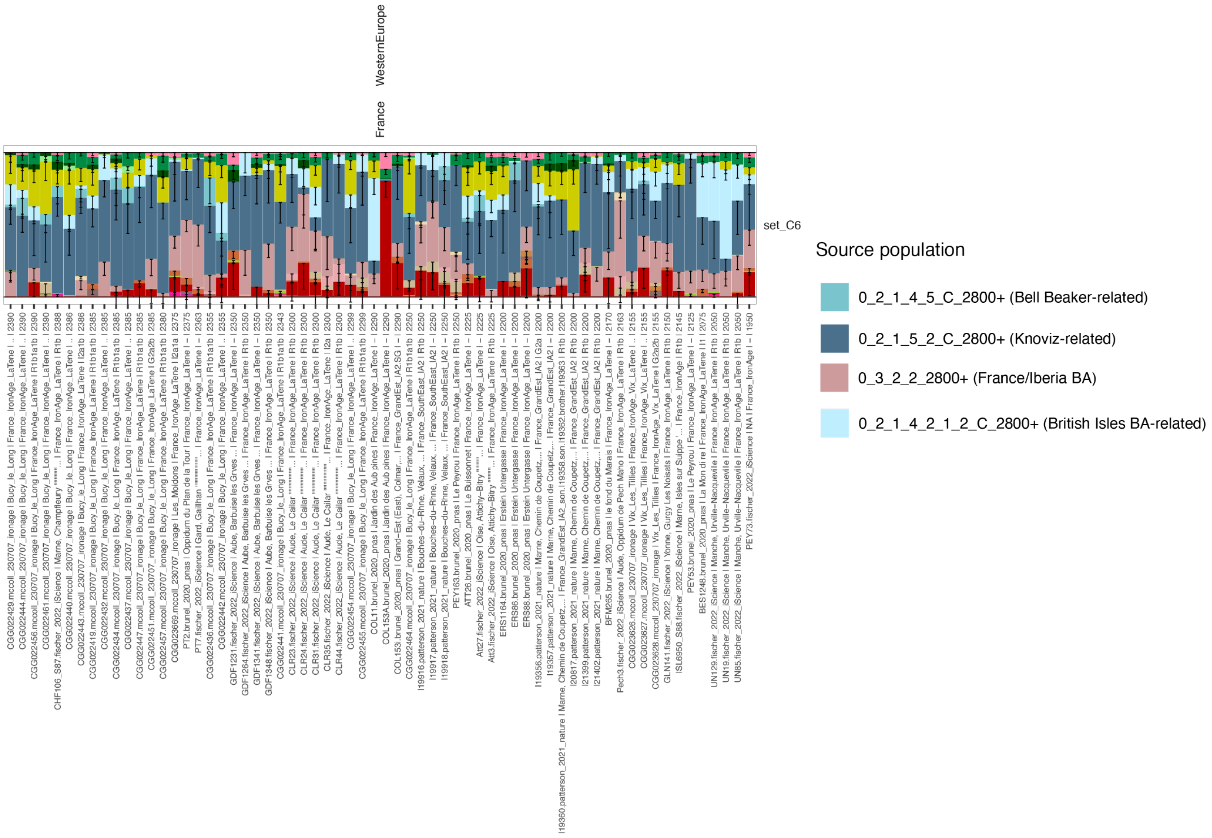


**Supplementary Fig. S1.24. mixture modelling results for French individuals between 2400-1900 BP, for mixture modelling sets C6.** Relevant sources are shown in the legend.

Our results stand in contrast to recent studies on Iron Age France[49] and Southwest Germany[19], where qpAdm analyses do not reject models in which early Iron Age populations are modelled by local Bronze Age populations. This different outcome likely results in part from the application of less fine-scale methods and inclusion of fewer ancient samples. The benefit of IBD mixture modelling in providing resolution at the individual level is also apparent from the ability to detect diversity within a population, when compared to qpAdm analyses which are typically run at a population level to achieve statistical significance. Notably, the identification of the outlier individuals from Iron Age Normandy, France with elevated British-Irish Bronze Age ancestry, is consistent with archaeological evidence for this site suggesting origins in Southern England[49].

To explore further the suggested genetic continuity between Bronze and Iron Age individuals in Western Europe[19], we turn to the auxiliary dataset, which included the Bronze Age Lech Valley individuals used as a potential source in [19]. Fitting with expectations for Celtic individuals from the main dataset, we find the Celtic elite-related individuals included from [19] to be modelled primarily as Knoviz related. Consistent with the spatial inference on the origin of individuals from that study, we find in addition to Knoviz-related ancestry, MBG017 is modelled with Bronze Age Anatolian and Balkans Bronze Age ancestry, and MBG009 to be modelled with Scandinavian ancestry.

For many of these individuals, the MBA Lech Valley individuals could not be excluded as a source using qpAdm [19]. We find individuals from the Lech Valley in southern Germany dated between ~3950-3700 BP to be distinct from earlier Lech Valley individuals (4300-3950 BP) (Fig S1.x1). This is apparent from the large proportion of Italian/Switzerland Neolithic Farmer ancestry (mixture modelling Set C4), Balkans Bronze Age (mixture modelling Set C5) and Knoviz-related (mixture modelling Set C6) ancestry. We also note the presence of Italian Neolithic ancestry, rather than Iberian Neolithic ancestry, similar to Knoviz-related individuals. Combined, these results point to these later Lech Valley Bronze Age elite individuals being linked with a more southern population, consistent with the Knoviz-related ancestry forming between 4000 and 3300 BP. We note that the proportion of the Balkans Bronze Age and Knoviz-related ancestry is less in these individuals than in the later Celtic Elites (Fig S1.25), and most Hallstatt individuals from France, Germany and Austria (Extended Data Fig. 3), suggesting the Lech Valley individuals are related to, but not the direct source of the later Urnfield expansions.

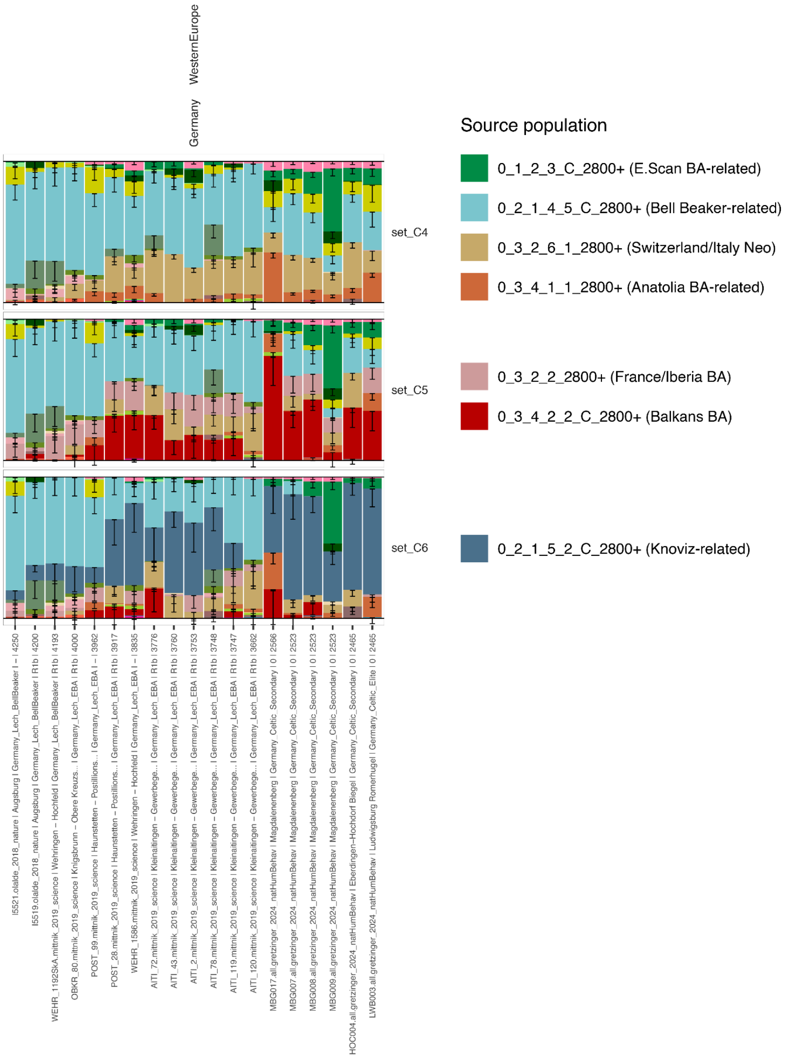


**Fig S1.25. IBD mixture modelling results for Bronze Age Lech Valley individuals and Iron Age Celtic Elite-related individuals for mixture modelling sets C4-C6.** Relevant sources are shown in the legend.

In general, we find the British Iron Age individuals from [78] to be modelled similarly to other individuals from the British Isles (Fig S1.26). Of particular note is WBK02, who was marked as an genetic outlier, suggested to be from continental Europe[78]. Consistent with this suggestion, when using source Set C7, we find this individual to be modelled with only Knoviz and French Bronze Age ancestry, lacking British Isles ancestry, indicative of recent immigration from France. Similarly, WKB01, who was also marked as an outlier in the original study, was found to have links to Derbyshire individuals[78]. Here we find this individual is distinct from others from Winterborne Kingston, as they lack the Knoviz-related ancestry, and are instead modelled with only British Isles-related ancestry (Set C6), similar to some individuals from Derbyshire (Supplementary Fig S1.6).

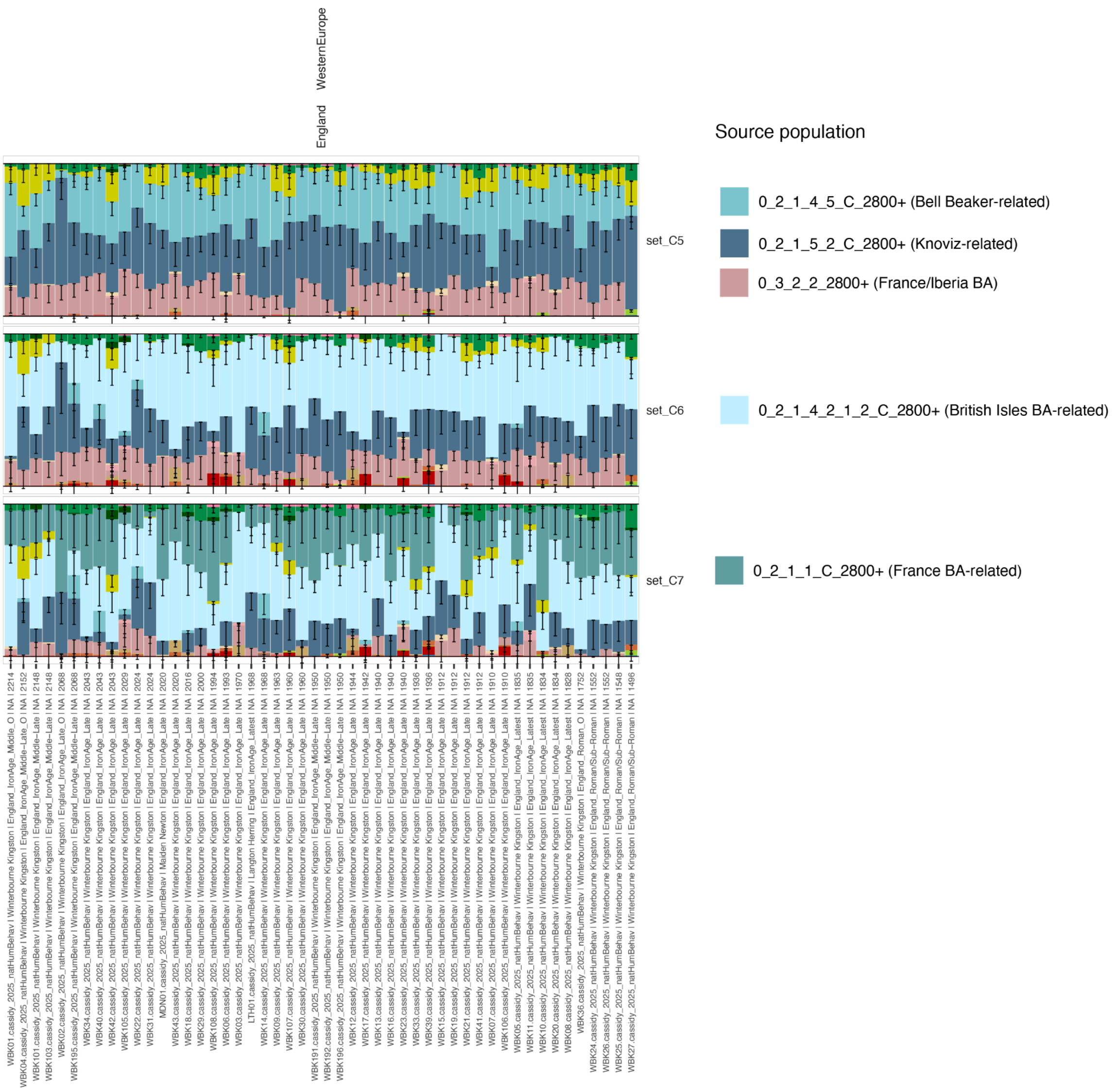


**Fig S1.26. IBD mixture modelling results for Celtic individuals from the British Isles from Cassidy et al [78] for mixture modelling sets C5-C7.** Relevant sources are shown in the legend.

### Supplementary Note S2. qpAdm Analyses

Hugh McColl[1,2] and Martin Sikora[1]

[1]Lundbeck Foundation GeoGenetics Center, Globe Institute, University of Copenhagen, Copenhagen, Denmark

[2]Department of Historical Studies, University of Gothenburg, Gothenburg, Sweden

While qpAdm may not have the resolution to detect the fine scale demographic shifts (for example the inability to resolve a specific source population or to detect discontinuity resulting from the influx of a closely related population), a result that directly contradicts the IBD mixture modelling would need to be accounted for. We therefore set out to test the major finding of this paper, that three distinct migrations occurring throughout the Bronze Age occurred associated with Bell Beaker-related populations. The first, being the initial southward spread of Steppe ancestry, the second being the spread of admixed individuals from nearby France/Iberia to the British-Irish Isles, and the last being the spread from Central Europe.

As targets, we binned individuals from England, where dense sampling across the entire Bronze and Iron Age is present, and from France, where a large number of new samples improves resolution. Bronze and Iron Age individuals were binned into four groups, 4800–4000 BP, 4000–3200 BP, 3200–2800 BP, 2800–2470 BP and 2470–2200 BP. Outliers with high Farmer ancestry were excluded. The full list of individuals in each target group can be found in Supplementary Table S2.4.

For all tests, we fixed 6 individuals from the British-Irish Isles (0_2_1_4_2_1_2_C_2800+, 4150–3585 BP). We then assembled a series of potential Neolithic and early Bronze Age sources from across Europe, including populations primarily modelled with Neolithic Farmer ancestry from the British-Irish Isles (0_3_1_2800+), France/Iberia (0_3_2_4_4_1_2800+), Denmark (0_3_3_2_2_2_C_2800+), Italy (early: 0_3_4_3_5_4_C_2800+, and late: 0_3_2_5_2_2_2_2800+), Czech Republic (0_3_3_1_2800+), Poland (Globular Amphora Culture: 0_3_3_3_C_2800+) and Bronze Age Anatolia (0_3_4_1_1_2800+). We also included individuals from Bronze Age France/Iberia (0_3_2_2_2800+) and from Hungary/Serbia (0_3_4_2_2_C_2800+) who are modelled with small proportions of Bell Beaker-related ancestry.

For all tests, we set a series of outgroups populations: Corded Ware-related (0_2_2_2_1_1_C_2800+), Yamnaya-related (0_2_2_3_2800+), Iran Neolithic (0_3_4_1_2_1_C_2800+), Caucasus Hunter-Gatherer related (0_3_4_1_7_2800+), Early Anatolia Farmer-related (0_3_4_3_3_2_2_2800+), East Asia (0_4_2_1_4_2800+), Scandinavian HG-related (0_5_2_2_1_1_2800+), Ukraine EHG-realted (0_5_2_2_1_2_1_2800+), Russia EHG-related (0_5_2_3_2_2800+),Italy WHG-related (0_5_3_1_3_2800+), France HG-related (0_5_3_2_1_1_2800+), Iberia HG-related (0_5_3_2_2_1_2800+), Lithuanian/Latvian HG-related (0_5_3_2_3_2800+), Danish HG-related (0_5_3_3_1_2800+) and Africa Neolthic-related (0_6_3_2800-).

For 1pop models, the British-Irish Isles Bronze Age source was set as the ‘left’ group, and all the rotation groups and outgroups were set as the ‘right’ group. For 2pop models, the British-Irish Isles Bronze Age source was fixed in the ‘left’ group, and the outgroups were fixed in the ‘right’ group. We then cycled through each of the rotation groups, placing one in the ‘left’ group and all others in the 'right'. The 3pop models had the same setup as the 2pop models, but including all possible combinations of two populations from the 'rotate' group, together with the British-Irish Isles Bronze Age source as the 'left' group, and all others in the 'right' group.

The full set of qpAdm results can be found in Supplementary Tables S2.1-2.3. The full list of individuals can be found in Supplementary Table S2.4. P-values for the various qpAdm tests are also shown in Supplementary Fig. S2.1.

*Discussion*

From the IBD mixture modelling results, the occurrence of three distinct migrations is apparent on the British-Irish Isles. When using qpAdm, only in the earliest period (4800–4000 BP) do we see individuals being consistent under the 1 pop model, with the local Bell Beaker source (Supplementary Fig. S2.1). In contrast, in the Middle Bronze and Late Bronze Age, only 2 population models are not rejected, with multiple possible Neolithic sources, or Bronze Age France/Iberia, but not Bronze Age Serbia/Hungary. In the Iron Age transects, no 1 or 2 population models are consistent with the data. In the first (2800–2470 BP), the only models not rejected include Bronze Age Serbia/Hungary. In the second (2470–2200 BP), the only non-rejected model (with marginal significance) includes Bronze Age Serbia/Hungary.

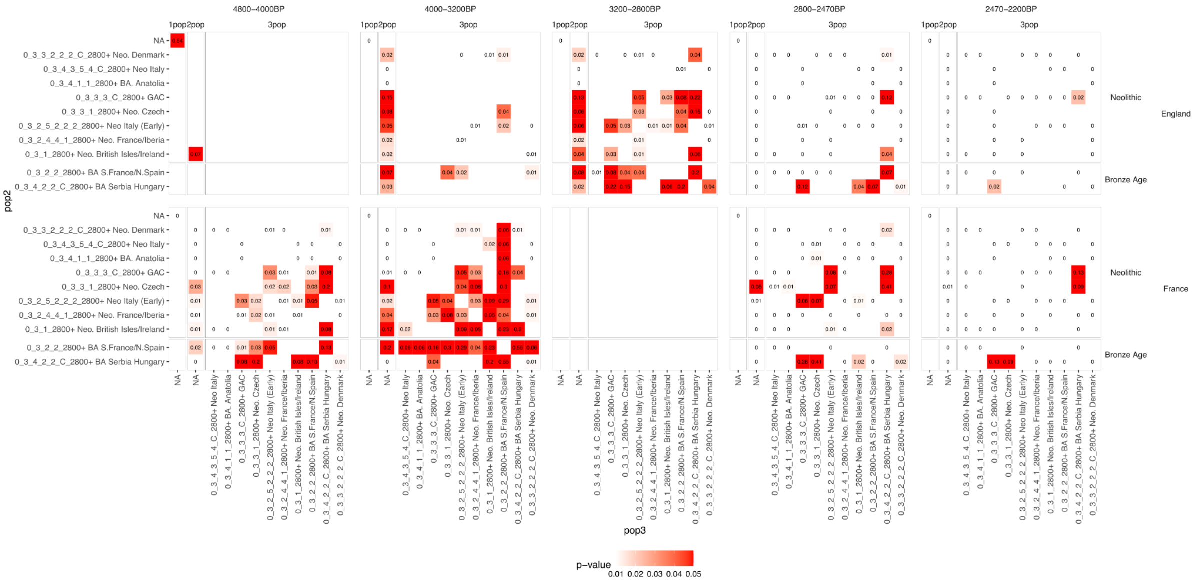


**Supplementary Fig. S2.1. P-values for rotating qpAdm models for the five time transects from England and France.** For all models, Pop1 is fixed as the English Bell Beaker-related cluster. For the 2 population model, the second pop is listed on the Y axis. For the three population mode, the third population is listed on the X axis. P-values for feasible models are shown for each model.

In France, no 1 population model is consistent with the data (Supplementary Fig. S2.1). For the 4800–4000 BP bin, a Neolithic Farmer group or the local Bronze Age source are weakly significant. For the 4000–3200 BP bin, multiple Neolithic Farmer groups or the local Bronze Age group are not rejected, but not the Bronze Age Hungary/Serbia group. During the Iron Age a single Neolithic Farmer is feasible in a 2 population model for the earlier bin, but not the later. In both, multiple 3 population models including Bronze Age Hungary/Serbia but not France/Spain fail to be rejected.

### Supplementary Note S3. Chromopainting results

William Barrie[1] and Hugh McColl[2,3]

[1]Department of Genetics, University of Cambridge, Cambridge, United Kingdom

[2]Lundbeck Foundation GeoGenetics Center, Globe Institute, University of Copenhagen, Copenhagen, Denmark

[3]Department of Historical Studies, University of Gothenburg, Gothenburg, Sweden

Chromosome painting (or 'chromopainting') has often been used to infer both genome-wide ancestry proportions and local ancestry, including using grouped ancient samples as 'donor' or 'source' populations[84,85] due to its ability to detect fine-scale population structure based on patterns of haplotype sharing. However, more recently, IBD mixture modelling has been favoured for detecting very fine-scale structure, relying on shared genome tracts which are identical-by-descent rather than just genealogically nearest neighbours. Here, we tested whether the main results of this paper could be replicated using the more traditional chromosome painting method.

We used sparsepainter[86], a fast replacement of the previous chromopainter algorithm, to paint ancient samples with reference panels of other ancient samples. We then used Sourcefind[87] to infer admixture proportions. We ran five iterations of chromosome painting, with each iteration (or set) having a greater number of reference populations. Full lists of the reference populations used and their composite samples can be found in Supplementary Table S3.1. The spatial and temporal distributions can be visualised in Fig. S3.1, Fig. S3.2, Fig. S3.3 and Fig. S3.4. Three sets were chosen based on the key findings of the paper. (1) A set including representatives of the Bell Beaker and Corded Ware Cultures (Set CWC_BB), to test whether the distinction between the two regions visible from IBD mixture modelling persisted through time, (2) A set including Bronze Age individuals from Southeast Europe and Southwest Europe (Set SEEu_SWEu) to broadly test the timing and direction of migrations that occurred by the Iron Age, and (3) a set including ancestry related to that of the Knoviz (Set Knoviz) individuals.

Source groups were manually curated, including individuals with high proportions of the corresponding ancestries. This allowed for many more individuals to be used as sources compared to the IBD mixture modelling. All other ancient samples in the dataset with their region listed as in either SouthernEurope, WesternEurope, CentralEasternEurope, or NorthernEurope were painted as 'target' samples (n=~3350 depending on the reference panel size). Finally, each sample in the reference panel was also painted using the same reference panel but excluding that sample ("leave one out").

Before painting, we filtered for autosomal SNPs with 1000G mappability mask, info score>0.5, and 1240k capture sites (n=690,211). We then ran a standard chromosome painting pipeline with the following steps:

1. Generate weight file through ref-vs-ref painting (sparsepainter -weight option). This generates weightings for sites depending on how likely they are to be painted as the reference population they have been assigned to.
2. Paint targets using weightfiles.
3. Paint reference samples using weightfiles without themselves in the reference panel (sparsepainter -loo option).

In all steps we estimated a fixedlambda value using chromosome 20 and used that for all other chromosomes.

We summed the ‘chunklengths’ per reference population across all chromosomes for both the target and reference paintings, and used this as the input to sourcefind (example code: https://github.com/will-camb/misc/blob/a05a92177c5fd70995b353bdcd4ab44640a81647/sparsepainter2sourcefind.py). We ran sourcefind with the following options: num.surrogates: 5, exp.num.surrogates: 3, num.slots: 100, num.iterations: 2000, num.burnin: 500, num.thin: 50, which we found was sufficient to converge on a stable estimate. We took the mean of the estimates across iterations as the final estimated ancestry proportion.

**Results**

Full estimates from this analysis per sample can be found in Supplementary Table S3.2.

In Supplementary Fig. S3.5, we show all individuals from England between 4,000 and 1,000 BP. Prior to 1600 BP, individuals are associated with Bell Beaker and Celtic Cultures. After 1600 BP, individuals are associated with Germanic Saxon and Viking Cultures. We see a the shift in ancestry from Bell Beaker-related to Corded Ware-related at this time, consistent with the IBD mixture modelling results.

In Supplementary Fig. S3.6, we present chromopainting results consistent with the IBD mixture modelling results shown in Supplementary Fig. S1.7. For example, in England, early individuals (4800-4000 BP) are modelled primarily as Bell Beaker-related (BBEarly_n40) or local Neolithic Farmer ancestry (neoBrIs_n29). Individuals from the 4000-3200 and 3200-2800 show small increases in their proportions of many ancestry sources, including individuals with high proportions of the Southwest European Bronze Age ancestry (baSwEu_n19) and Neolithic Iberian (neoIb_n29) ancestry, consistent with later migrations from Ibera and Southwest France. In these two bins, some Southeast European Bronze Age (baSeEu_n27) ancestry is also present to varying degrees, but by the later periods (2800-2470 and 2470-2000 BP) it is present in more individuals and in higher proportions. We see changes in France and the Czech Republic that similarly correspond to the results from the IBD mixture modelling.

In Supplementary Fig. S3.7, we present chromopainting results consistent with the IBD mixture modelling results shown in Supplementary Fig. S1.12. As with the IBD Mixture modelling, much of the ancestry previously modelled as Southeast European Bronze Age (baSeEu_n27) is now modelled as Knoviz-related (knoviz_n47). These results and the IBD mixture modelling show similar changes through time in England, France and the Czech Republic.

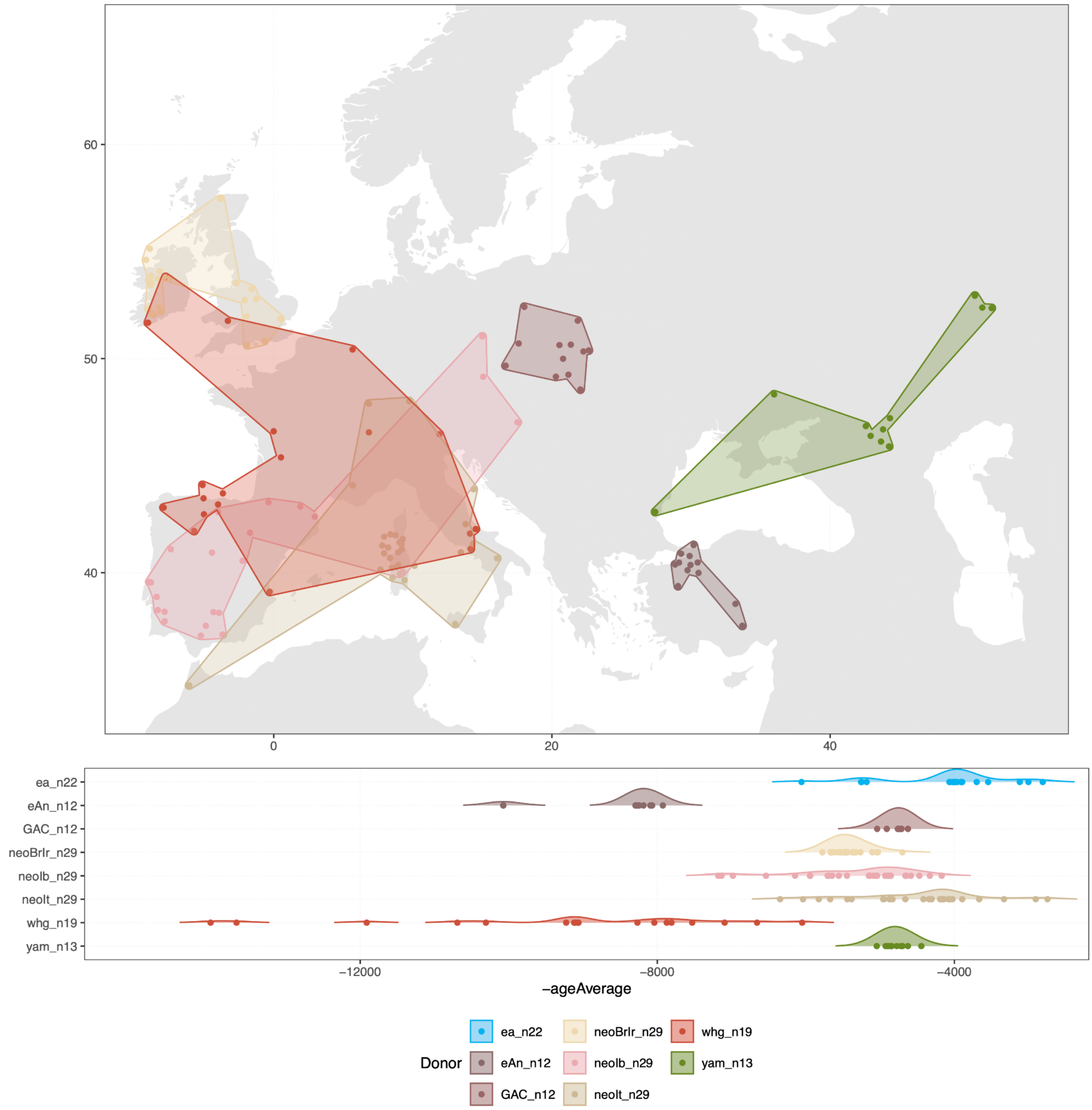


**Supplementary Fig. S3.1. Spatial and temporal distribution of the subset of samples used in all from Chromopainter Sets.** Additional samples from specific sets are shown in Fig S3.2, Fig S3.3, and Fig S3.4.. The East Asian and African sources are not shown on the map.

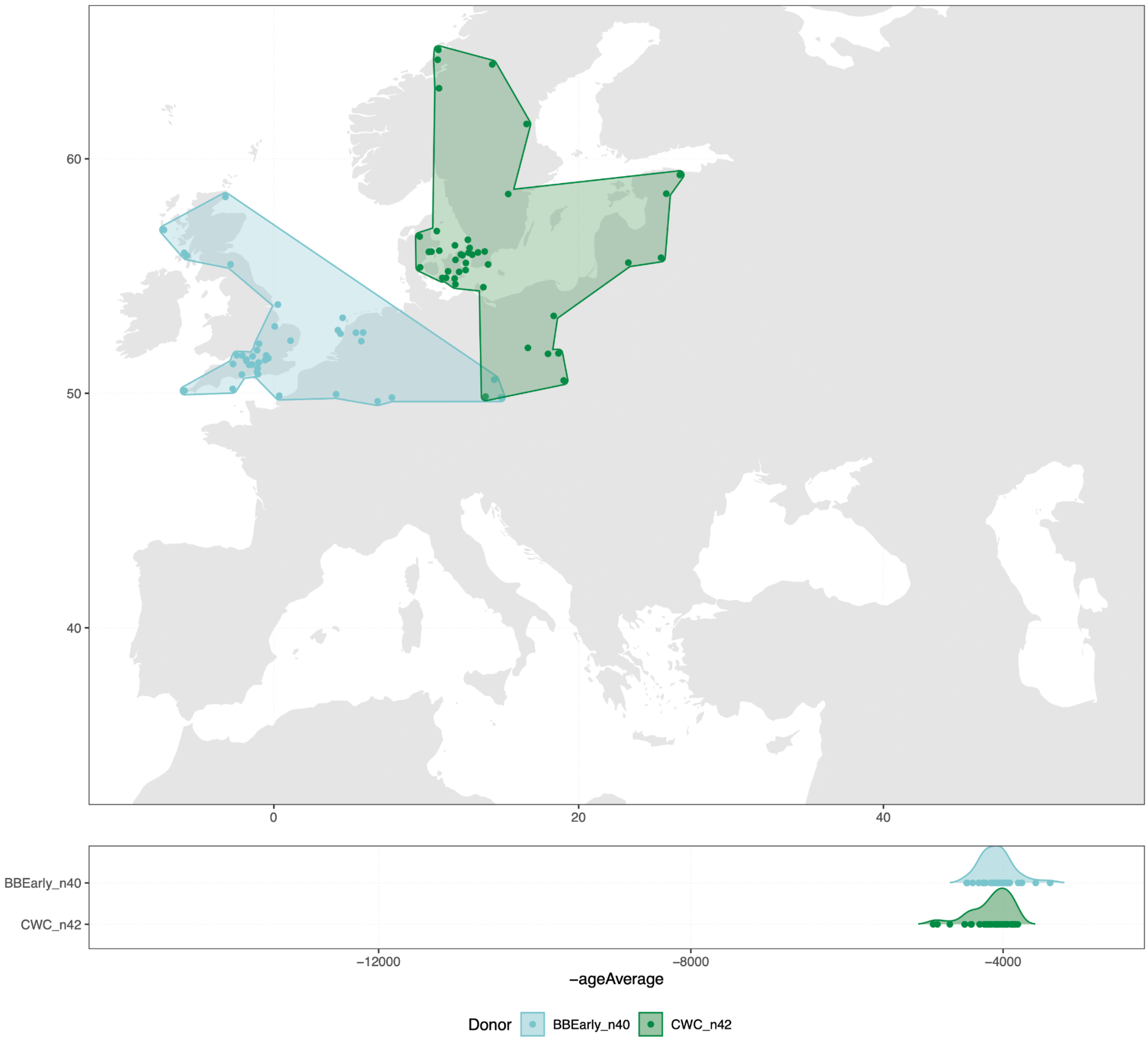


**Supplementary Fig. S3.2. Spatial and temporal distribution of the additional samples used in from Chromopainter Set CWC_BB.** Samples common to all chromopainter sets are shown in Fig. S3.1. The East Asian and African sources are not shown.

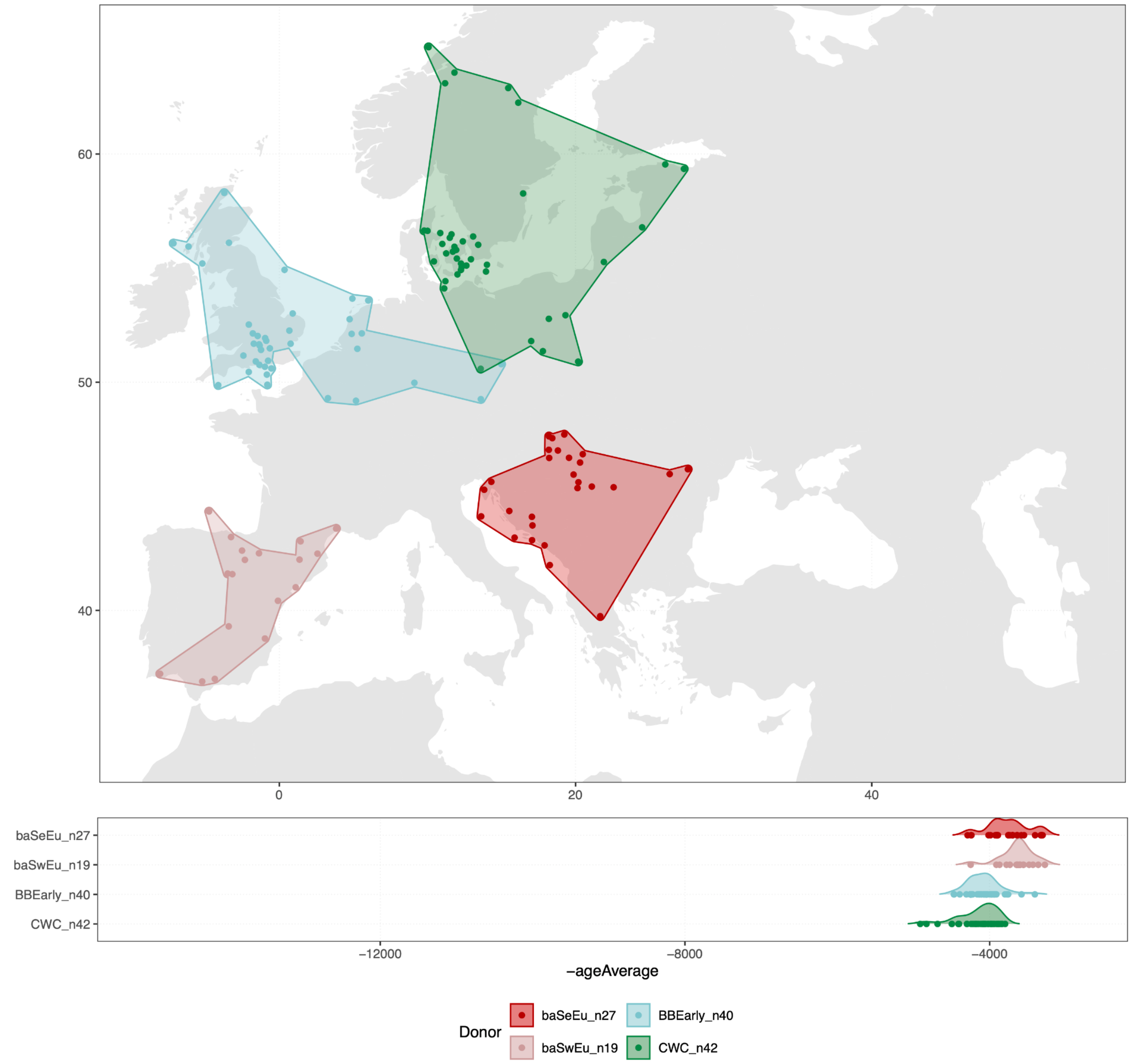


**Supplementary Fig. S3.3. Spatial and temporal distribution of the additional samples used in from Chromopainter Set SEEu_SWEu.** Samples common to all chromopainter sets are shown in Fig. S3.1. The East Asian and African sources are not shown.

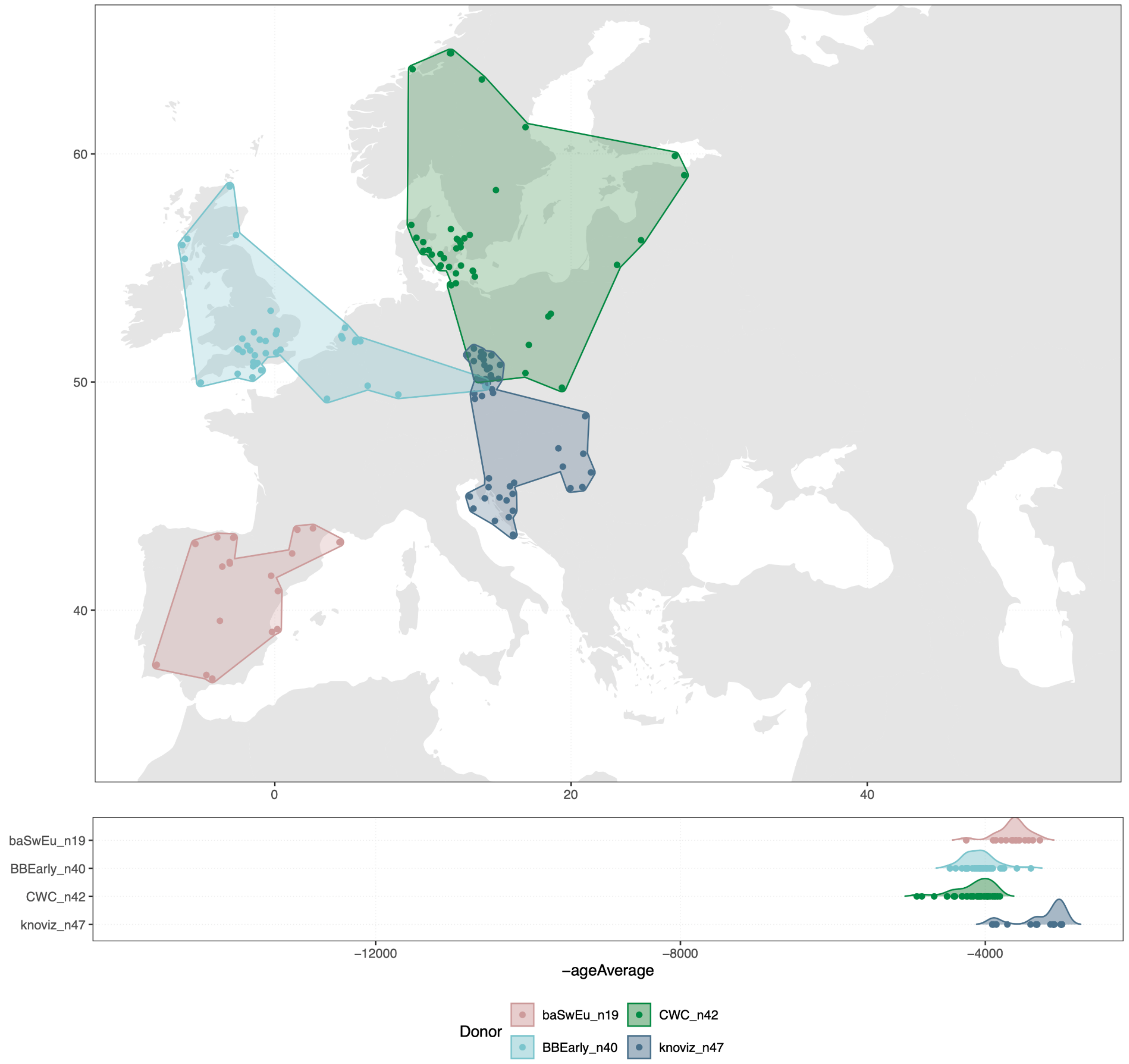


**Supplementary Fig. S3.4. Spatial and temporal distribution of the additional samples used in from Chromopainter Set 'Knoviz'.** Samples common to all chromopainter sets are shown in Fig. S3.1. The East Asian and African sources are not shown.

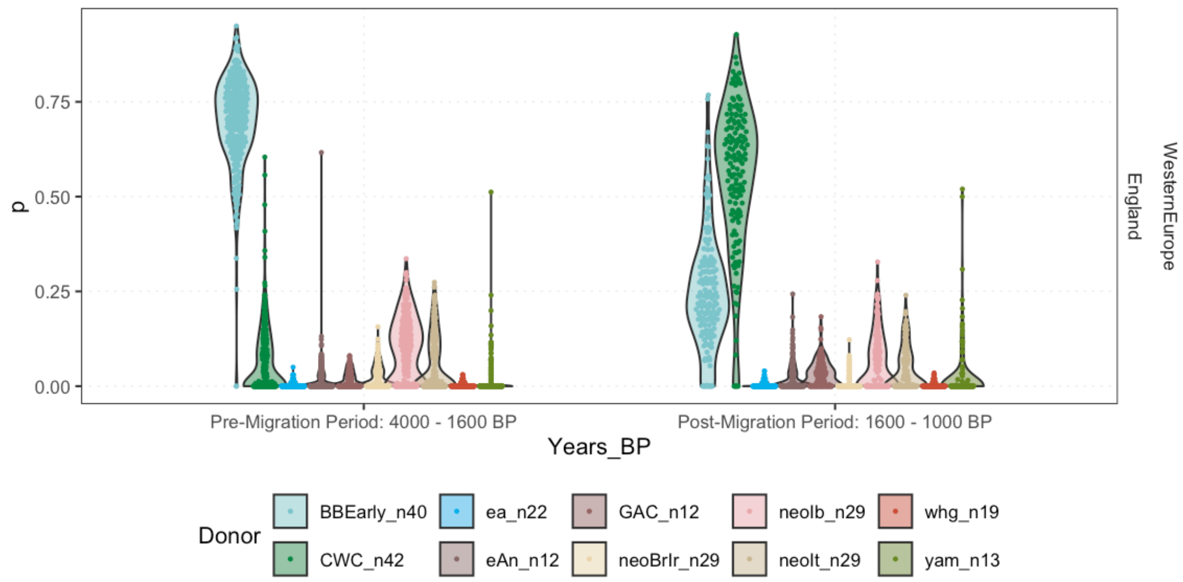


**Supplementary Fig. S3.5. Chromopainting results highlighting the distinction between Corded Ware (green) and Bell Beaker (blue) related ancestry in Celtic and Germanic-related populations respectively.** All individuals from England from 4000-1000 BP are shown, split before and after the arrival of the Saxon population.

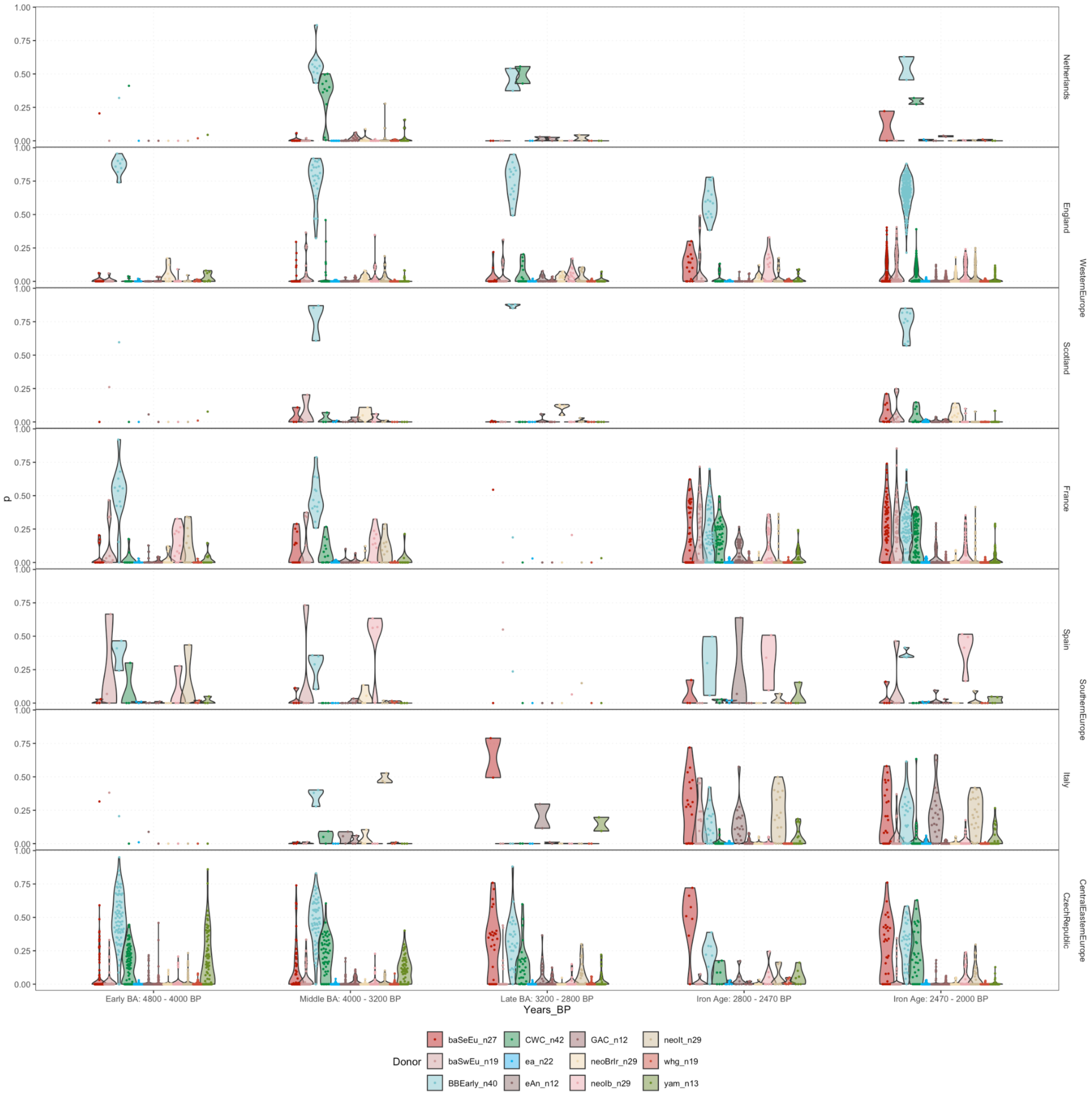


**Supplementary Fig. S3.6. Chromopainting results highlighting the variation in Bronze Age ancestry from the Balkans (Hungary/Serbia) and Southwest Europe (France/Iberia).** Only individuals with more than 2% Yamnaya ancestry modelled using Set C1 are shown. These results correspond to the IBD mixture modelling results in Supplementary Fig. S1.11.

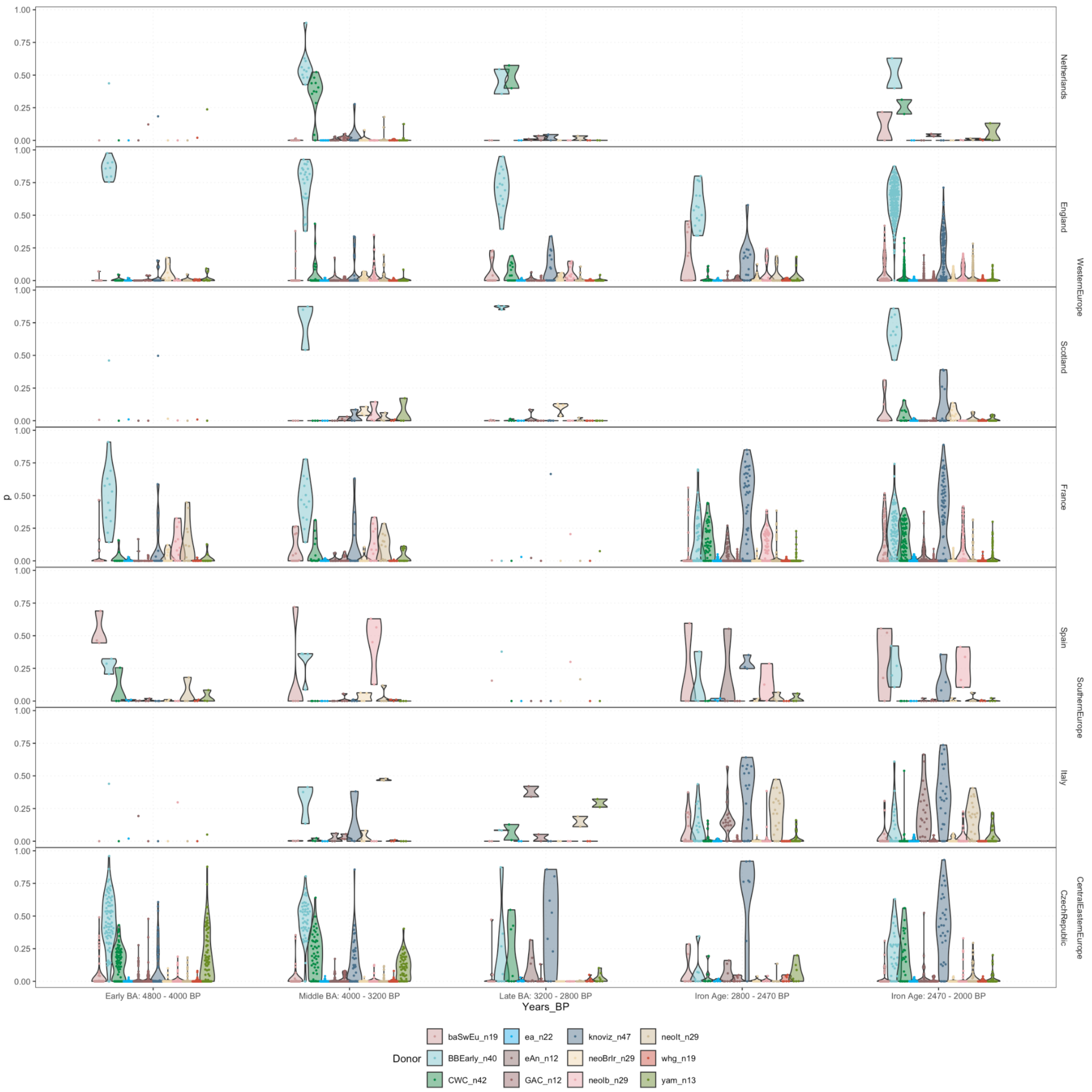


**Supplementary Fig. S3.7. IBD mixture modelling results using Set C5, highlighting the Knovíz-related ancestry.** Only individuals with more than 2% Yamnaya ancestry modelled using Set C1 are shown. These results correspond to the IBD mixture modelling results in Supplementary Fig. S1.12.

### Supplementary Note S4. Y-chromosome results

Thomaz Pinotti[1,2]

[1]Lundbeck Foundation GeoGenetics Center, Globe Institute, University of Copenhagen, Copenhagen, Denmark

[2]Laboratório de Biodiversidade e Evolução Molecular (LBEM), Universidade Federal de Minas Gerais, Belo Horizonte, Brazil

#### S3.1. Temporal and spatial co-occurrence of haplogroup R-P312 and Beaker-related ancestry

Since IBD mixture modeling provides per-individual ancestry proportions, we aimed to examine, at a fine scale, the chronological co-occurrence of distinct ancestries and Y-chromosome haplogroups. Supporting previous studies[37,88] our analysis revealed a striking correlation between the arrival of haplogroup R-P312 – downstream of R-M269 > L23 > L51 > L151 – and the spread of Beaker-related ancestry.

Across all analyzed regions, no individual carrying the R-P312 haplogroup was found to predate the earliest individual modelled with Beaker-related ancestry using IBD mixture modelling. In nearly all cases – including the Netherlands[37], Italy[37,89], Spain[18,38], England[37], Wales[17], Ireland[45], Germany[43] and Czech Republic[37] – the oldest individual with Beaker-related ancestry was also himself the oldest confirmed carrier of R-P312.

There were a few exceptions: in France[46,50], low coverage prevented precise subhaplogroup classification for the oldest individual with Beaker-related ancestry; in Scotland[37] and Poland[37,90], the earliest individuals were female; and in Portugal[37,90], the oldest male carried a

Neolithic haplogroup (I-L160). Nevertheless, in each of these cases, the next oldest individual with Beaker-related ancestry did carry R-P312, reinforcing the association between this haplogroup and the Beaker-related genetic profile.

Furthermore, in Central Europe and the Balkans, where individuals with Steppe ancestry (Corded Ware or Yamnaya-related) predate the arrival of the Bell Beaker Culture, those carrying haplogroups downstream of R-M269 belong to either R-Z2103 or rare or basal R-L51 non-R-P312 lineages[91,92] until the arrival of Beaker-related ancestry. This further supports the relationship between Beaker ancestry and R-P312. These rare paternal lineages include basal R-L51*, R-PF6538*, and R-L151*, as well as closely related haplogroups such as R-Y215377, R-BY44535, and R-Z2118.

The co-occurrence of dominant R-P312 and R-U106 and these rare related haplogroups suggests that this region may have been the area of R-L51 diversification. While these related paternal lineages never reach high frequencies, they appear more commonly in later periods within the archaeogenetic record of Central Europe, the Balkans and the Italic Peninsula, including in non-Indo-European speaking Etruscans[17,93,94]. This pattern likely reflects a more complex and layered population history of Steppe ancestry in these regions.

##### S3.2. Spatial distribution of subhaplogroups of R-P312 track population movements in the Middle and Late Bronze Age

Haplogroup R-P312 resolves itself in a polytomy, with downstream lineages showing some level of geographical structure, particularly among Bronze Age individuals. Subhaplogroup R-L21 (and its descendant R-DF13) is predominant in Britain and Ireland; R-DF27 (and R-Z195) in Iberia and France; R-U152 (and R-L2) in Central Europe and Northern Italy; and R-

DF19 and R-Z30597 potentially associated with the Netherlands and Scotland, respectively, though sample sizes for these two are very limited. This regional distribution of R-P312 subhaplogroups provides a framework to explore population movements during the Bronze and Iron Ages. Additionally, the European Farmer-related haplogroup G-L497 also seems to be a geographically structured lineage, being largely restricted to Central Europe and Italy during the Bronze Age, and could also be used to track increased connectivity between regions in the archaeogenetic record.

However, it must be noted that, consistent with autosomal DNA evidence, Western Europe exhibits a strong paternal lineage continuity from the Bronze Age to the present, particularly when considering R-P312 subhaplogroups. In most regions, modern males predominantly descend from the same subhaplogroups that were already common in the Bronze Age. Notably, in Ireland and the Basque Country, R-L21 and R-DF27, respectively, account for over 80% of all present-day male lineages[95–100].

Allele-frequency-based [17] and haplotype-based methods detect an arrival of continental ancestry in Britain during the Middle Bronze Age (3,200-4,000 BP). Consistent with this, we detect the R-DF27 sub haplogroup – associated with France and Iberia – among a few individuals during and after the Middle Bronze Age [17,37], which could be evidence of this signal of gene flow from these regions. Interestingly, one of these individuals from the Middle Bronze Age, from Cliffs End Farm, Kent, was shown to be a genetic and isotopic outlier[17].

IBD mixture modelling results have demonstrated gene flow between Central Europe (specifically, Urnfield-associated Knovíz Culture) and Britain and Iberia, who share an ancestry source which is also present in Hallstatt, La Tène and other contexts in France and Germany. Supporting the autosome evidence, we document multiple occurrences of typically Central European lineages R-U152 (and its downstream lineage R-L2) and G-L497 in Britain by ~2,300 BP[17] and France by ~2,600 BP[17,49,50]. On the continent, those individuals are mainly from Hallstatt, La Tène/Gaulish contexts. Though significantly later, we also find haplogroup R-U152 in Medieval Belgium, where Celtic languages had been spoken in the Iron Age[101].

In Iberia, IBD mixture modelling suggests gene flow with Central Europe to start by 2,500 BP, but very few male individuals have been previously analyzed from Spain or Portugal between the crucial time period between 1,800 and 2,800 BP [17,18] most of them at low coverage. Their paternal lineages are dominated by local R-DF27 and its subhaplogroups [102], but one individual from Northeastern Spain belongs to haplogroup R-U152 (L2), which can be tentatively linked with the Urnfield presence in the region [103]. Additionally, three present-day Northeastern Spanish individuals from the Iberians from Spain (IBS) 1000 Genomes population also carry R-U152 (L2) haplogroups.

We are not able to evaluate the paternal lineage transect in Ireland due to the absence of sampled individuals from Ireland in the period between 3500 and 1200 BP.

### Supplementary Note S5. Linguistic Supplementary

John Koch[1] and Guus Kroonen[2,3]

[1] University of Wales Centre for Advanced Welsh and Celtic Studies

[2] Department of Nordic Studies and Linguistics, University of Copenhagen

[3] Leiden University Center for Linguistics, Leiden University

#### 1 Background: The early Celtic languages

Celtic is a prominent branch of the Indo-European languages family. By the Iron Age, Celtic language varieties were spoken across much of Europe. All of these diverged from a common linguistic ancestor, Proto-Celtic, or at least from a Common Celtic dialect continuum[32], often estimated to have existed during the late 4th millennium BP[104], cf. c. 3200–2900 BP[30], c. 3205 (2515–3963 BP)[31] or at any rate after the start of the 4th millennium BP[105] and before c. 2600 BP[106]. On a deeper level, the Celtic branch is widely grouped with the closely related Italic branch, ancestral to e.g. Latin, under a Italo-Celtic subclade that arguably also included Lusitanian[14,107,108], but this subgrouping is not universally accepted[109].

Six pre-medieval languages are widely recognized: Celtiberian, Lepontic, Gaulish, Galatian, Brittonic, and Goidelic.

Celtiberian was spoken in Pre-Roman Iberia. It is predominantly recorded in inscriptions found in Northeastern Spain from the 2nd century BCE to the 1st century CE[110–112]. Some Celtiberian inscriptions are written in Roman script. The bulk of the earlier materials is in the native Celtiberian script, which derives from the Northeastern Iberian semi-syllabary used for the unclassified Iberian language, documented between the 6th and 1st centuries BCE. Extending beyond the part of east-central Spain inhabited by groups identified as *Celtiberes*, there is evidence of Ancient Celtic personal, place-, and group names further north and west, reaching Galicia, southwest Spain, and all regions of Portugal, such as the widespread Celtic place-name element *-briga* 'hillfort, town'. As there is evident dialect variation, the inclusive term 'Hispano-Celtic' may be used to distinguish the Ancient Celtic of east-central Spain from this more widespread evidence. The Tartessian language, known from inscriptions from Southwestern Iberia between the 8th and 5th centuries BCE, has sometimes also been classified as a Celtic language[113–115], but is alternatively taken as Indo-European though not certainly Celtic[112], certainly not Celtic[15], or a linguistic isolate[116–118].

Lepontic is attested from the 6th to 1st century BCE in inscriptions from northern Italy written in alphabetic scripts closely related to those used to write the neighboring non-Indo-European Etruscan language[119,120].

Ancient Celtic varieties usually called Gaulish are widely attested from the 5th century BCE across France and eastward into Central Europe around the Rhine, Alps, and upper Danube[121,122]. Gaulish is attested in Greek and Roman scripts. There are Celtic names and inscriptions (in Roman and Etruscan-related scripts) from northern Italy that are often classified as Cisalpine Gaulish. However, there is debate as to whether this evidence reflects, with Lepontic, a distinct Cisalpine Celtic language rather than grouping more closely with Gaulish north of the Alps[120,123].

Galatian appears in Anatolia following documented migrations that occurred after the Celtic attack on Delphi in 279/8 BCE. It is mostly written in Greek script and closely similar to Gaulish[119,124].

Brittonic names are recorded in pre-Roman times by Greek and Roman writers and occur also on the legends of native Late Iron Age coins using Roman script[125]. Ancient Brittonic is the direct ancestor of the Welsh, Breton, and Cornish languages of medieval and modern times.

Several Celtic names recorded by Greek and Roman authors and relating to Ireland can be interpreted as ancient forms of Goidelic. The language is attested in its pre-Old Irish form in c. 500 inscriptions in the indigenous ogam script, which begin by the 5th century CE[126]. Old Irish c. 600–900, sometimes called Old Gaelic, is the direct ancestor of the Irish, Scottish Gaelic, and Manx of medieval and modern times.

Lusitanian is a sparsely documented Pre-Roman Indo-European language of West-Central Iberia[82]. It is known from a small number of Lusitanian inscriptions, as well as from Latin inscriptions containing Lusitanian elements[81]. Due to its fragmentary nature, it is difficult to establish the exact linguistic affiliation of the language within the Italo-Celtic group[81,127]. It has been classified as an archaic form of Celtic[128–130], but is alternatively held to be non-Celtic and potentially closer to Italic than to Celtic[83,131]. At any rate, Lusitanian clearly did not share in the diagnostically Celtic innovation of the weakening and, most often, loss of **p*, as shown by Lusitanian *porcom* ‘pig’. Therefore, loss of IE **p* would have to be removed from the definition of Celtic to include Lusitanian in the family.

**2. Models for a Celtic homeland and dispersal**

The case for linking the initial spread of Yamnaya-related ancestry with the spread of Indo-European is relatively straightforward[42,43]. The genetic makeup of the incoming Yamnaya pastoralists and settled Early European Farmers had been long separate and was so different as to effectively rule out that they originally spoke the same language, mutually intelligible dialects, or languages that were demonstrably related at all. On the other hand, the subsequent emergence of Proto-Celtic (PC), as an Indo-European branch, is more complicated. This process occurred at least a millennium after the arrival of Steppe ancestry and requires more subtle methods of detection. We must therefore consider a range of Bronze Age prehistoric populations all of whom carried high levels of Steppe-related and variable proportions of Neolithic farmer ancestry. It is likely that many of the ancestry groups under consideration spoke early Indo-European languages, although Europe at this time still likely also hosted a range of extinct non-Indo-European languages. During this period, the prehistoric ancestors of the attested Celtic languages were linguistically closer to PIE and therefore more similar to one another as well as to the ancestors of the other branches.

The archaeogenetic evidence for the expansion of Steppe ancestry c. 5000 BP can be seen as ruling out models in which the Indo-European speech that became Celtic arrived in the historical Celtic territories c. 8000-6000 BP with the First Farmers then evolved into Celtic *in situ* there, either in all of these territories[132] or first the Atlantic façade then spreading from there[27].

Potentially relevant to the Iron Age geographic distribution of Celtic are three later, large-scale migrations across Western, Southern and Central Europe, detected in this study. The first is the arrival of (Bell Beaker-related) Steppe ancestry between 4300 and 4000 BP, with origins around the Netherlands and Northern France. Two others occurred in the Bronze Age[cf. 17,133]. Of these, the second involves migrations to the British Isles between 4000-2800 BP, with origins in mainland Western Europe. The final occurred between 3200-2800 BP, with origins in Central Europe and associated with spread of the Urnfield and Hallstatt Culture across Western Europe, reaching the British Isles by at least 2800 BP, and Iberia by at least 2500 BP. As such, the suitability of each as a potential vector for Celtic must be considered.

When considering the early Bell Beaker migrations as the vector of Celtic, we find that the geographical range of the migrations is roughly consistent with that of the distribution of Celtic in the Iron Age. Beaker groups have accordingly been seen as plausible Celtic forerunners[36,134–136]. However, it is doubtful that all or even most of the PIE > PC sound changes[137,138] could have been completed by 4300-4000 BP (see above). For that time it would be more accurate to speak of the post-PIE dialects ancestral to Celtic, rather than Celtic itself[30]. Such a model would entail the early Indo-European dialects that later became Celtic first spreading with the Beaker expansion, then evolving *in situ* from post-Proto-Indo-European to Celtic[27,132,136,139].

This model requires some mechanism for linguistic innovations to be shared between cognate dialects across what had been the extent of the Beaker phenomenon and then, 1500 years later, that of the Ancient Celtic languages. Patterns of long-distance contact that possibly favoured the sharing of linguistic innovations include: the spread of standardized high-tin bronze from Britain and Ireland across mainland Europe c. 4100-3400 BP[136,140,141], elite exchange during the Early to Middle Bronze Age across the Channel/Southern North Sea ‘maritory’[142], and sea-crossing trade of copper from mining sites, such as Ross Island, Ireland

(4400-3900 BP)[143], Great Orme, Wales (3700-3400 BP)[142,144,145], and Late Bronze Age Southwest Iberian mines[146,147]. While the aforementioned interactions require only relatively small numbers of mobile specialists, the Bronze Age French/Iberian-related and Knovíz-related migrations supported by the data presented here imply more substantial population movements, which could also be integrated into this model. The shifts in proportions of Bell Beaker-related ancestry vs. Neolithic farmer-related ancestry between 4300 and 2500 BP, tending towards north-south equalization (fig. 2 above), suggest that the region that had been Bell Beaker territory and became Celtic formed a socio-cultural area[136,139,141] in the intervening period, with significant movement of population across it in both directions. In this model, the archaism of Celtiberian and non-Celticity of Lusitanian can be explained as an effect of diminished contact in Iberia with Gaul and Central Europe after the Bronze-Iron transition c. 2900 BP[148]. The disparities in the levels and sources of Neolithic farmer-related ancestry[45] (and Extended Data Fig. 4) are possibly reflected in early differentiations between the Celtic dialects. With the 'Celtic from the West' model, both Lusitanian and Celtiberian formed in Southwest Europe.

To explain Celtic as a whole, the second migration detected in this study, i.e. the Bronze Age French/Iberian-related ancestry c. 4000–3200 BP, is not a good fit. Particularly the earlier part of this date range would seem too early for the completion of all the PIE > PC sound changes, though this vector could not be decisively ruled out on that basis. The more serious defect is the absence of this ancestry type in the individuals from the Czech Republic, a region for which there is Ancient Celtic linguistic evidence, as well as being a formative area for the La Tène Culture c. 2400 BP. The spread of Celtic from the Atlantic into Central Europe would have occurred in the opposite direction to the Hallstatt migrations[59]. In addition, this interpretation implies that the later Knovíz-related migrations had no significant linguistic impact on Britain, despite a significant genetic impact. The 'Celtic from the Centre' model, which situates the homeland in France, and involves migrations from there during the Bronze Age across Western and Central Europe[15], suffers from the same defects. Likewise, the hypothesis of a Celtic homeland in northern Italy, emerging there from the local Urnfield Culture of the Canegrate period (33rd c. BP)[14], is not supported by the spread of French/Iberian ancestry, nor of that of Knovíz.

The formation of the Knovíz-like ancestry by 3300 BP and its subsequent westward spread is, however, consistent with a traditional linguistic model according to which Celtic spread from Central Europe[34]. The close genetic link with Knovíz-related ancestry and the Hallstatt Culture is in agreement with proposed links between the Hallstatt Culture and early Celtic. The model in which Celtic spread across Europe from the East with a distinct Knovíz-like ancestry that formed by 3300 BP suggests that the threshold of Proto-Celtic occurred by this point in time, which is in accordance with linguistic consensus (see above). A linguistic phylogenetic basis for 'Celtic from the East' has additionally been argued on the basis of grammatical innovations shared by Celtic with Indo-Iranian, Baltic, Slavic, Greek, Tocharian and Albanian[149–151]. However, an eastern origin beyond the area of Bell Beaker-associated Bronze Age ancestries is not evidently supported by the current genetic data.

A model associating a decisive episode in the formation of Celtic Europe with the spread of Knovíz-related ancestry also potentially sheds light on the presence of the distinct linguistic structure of Iberia by 2500 BP[35]. The presence of the Lusitanian language in Iberia, alongside less ambiguously Celtic evidence, is consistent with a model of two Indo-European waves entering the Peninsula, with only the later wave having undergone the full set of PIE > PC changes. Genetically we see support for a model in which Iberia is impacted by two distinct

Bell Beaker-related migrations - the first the arrival of Steppe ancestry around 4300 BP, and the later appearance of Knovíz-like ancestry by 2500 BP. Thus if Celtiberian is a more recent intrusion, Lusitanian might have developed from the speech of local populations resulting from the initial wave[152,153].

Taken at face value, the date of c. 2500 BP for the arrival of Knovíz-related ancestry is too late to account for Celtiberian and evidence for Celtic spoken further west in the Iberian Peninsula. Going back to Bosch-Gimpera[154], the arrival of the Celtic groups in Iberia has been equated with the appearance of Urnfields in Catalonia c. 3200–2900 BP[57,58]. In the iconography of the Late Bronze Age warrior stelae of the Southwestern Iberian Peninsula, types of Urnfield weaponry have been identified dating c. 3200–3000 BP[155,156]. Celtiberian is widely recognized as archaic, the first Celtic sub-branch to split off from PC[14,137,157–161]. Secondly, there is wide acceptance for Celtic linguistic items in the Iberian Peninsula in the Early IA, i.e. before c. 2500 BP[112,115]. The spectacular Huelva hoard from SW Spain, dated to the 11th century BCE and often invoked in definitions of the Atlantic Bronze Age[27], includes metal objects in styles typical of Iberia, the British Isles and Central Europe[162]. In short, it looks as though Urnfield cultural impact reached wide areas of Iberia by c. 3200 BP, but its genetic effects first become evident only several centuries later. Due to the limited number of Bronze and Iron Age genomes from the region, the date of 2500 BP should be interpreted as a lower limit of the time of arrival, with future samples possibly helping to address this issue.

Whereas we find evidence that Iberia was impacted by two Bell Beaker-related migrations, we find that Britain and Ireland were impacted by three migrations. Unlike Iberia there is no evidence for a pre-Celtic Italo-Celtic language, limiting our ability to assess (A) whether these migrations brought an Italo-Celtic language related to Celtic and (B) whether the language of the earlier migrants had an impact on the later linguistic landscape of Insular Celtic at all. While the initial Bell Beaker-related wave would be later than Cunliffe's 'Celtic from the West' model, in which Indo-European arrived in Iberia with Neolithic farmers, the second potentially supports Koch's revised model in which the Indo-European dialects ancestral to Celtic arrived there with Bell Beaker ancestry, and from there to Britain along the Atlantic seaboard. If the third wave was seen as ancestral to all the attested Celtic languages that would contradict these two models, in that it provides hitherto unknown support for the 'Celtic from the East model'. Notably, V. Gordon Childe[34] envisaged a model in which Celtic arrived with a Late Bronze Age intrusion, taking place in the wider European context of the Urnfield Culture. However, the implied secondary spread of Celtic from Britain to Ireland, whether in the Late Bronze Age[163] or the Late Iron Age[164], was and remains archaeologically difficult to substantiate[165,166] and genomically difficult to evaluate due to an unavailability in our dataset of genomes from Ireland in the period between 3500 BP and 1200 BP.

Finally, a fourth migration, taking place in the Middle to Late Iron Age, potentially associated with Celtic-speaking groups arriving from Gaul[78], falls outside the scope of this study.

### Supplementary Note S6. Archaeological Supplementary

Kristian Kristiansen[1,2], Johan Ling[2] and John Koch[3]

[1]Lundbeck Foundation GeoGenetics Center, Globe Institute, University of Copenhagen, Copenhagen, Denmark

[2]Department of Historical Studies, University of Gothenburg, Gothenburg, Sweden

[3]University of Wales Centre for Advanced Welsh and Celtic Studies, Aberystwyth, UK

***Urnfield Cultures***

The origin and expansion of the Urnfield Culture from around 1350–1200 BC has been explained as a social and demographic expansion of populations, drawing parallels to the La Tène/Celtic migrations (see Figure 224–225 from[167]). We cite: 'The parallels to La Tène suggest a similar development – the formation of new chiefly hierarchies, followed by a major reorganization of settlement and economy, leading to a rise of strong, pioneer farming communities expanding into new habitats both locally and over long distances, supported by warrior chiefs' (page 385 from[168]). However, this expansion has also been discussed in terms of a ritual change from inhumation to cremation that expanded with a new religion but also that the pits with inhumations could represent slaves[51]. The early origin of cremation and the use of urns for the bones originated in Hungary from where it spread in subsequent centuries [169,170].

The Urnfield Culture reflects a more collective, centralized social organization based on staple finance, i.e. on surplus generated by intensive agrarian regimes, rather than wealth finance, while remaining under the strong political leadership of war chiefs[167]. This expansion and associated migrations were rooted in agrarian intensification, with the introduction of new crops such as millet and large-scale transformation of landscapes[171]. These advances supported significant population growth and facilitated the spread of the Urnfield Culture. This in turn is reflected in the formation of large, fortified settlements[171]. Also advanced metalwork of hammered cauldrons and cups for feasting and drinking flourished, with centers in Hungary and Bohemia, from where their products were exported widely across Europe[167].

Within just a few generations, this Urnfield expansion had incorporated both France and northeastern Spain into its sphere (Figure 41 in [167]). The warlike nature of the Urnfield expansions suggest they were instigators of larger geopolitical events, and social transformations, also reflected in large depositions of metal in hoards in some regions (see page 2015, Figure 8 in [172]). It has been suggested that the expanding Urnfield communities moved westward to disrupt the Tumulus Culture's control over the distribution of copper supplies originating from the Italian Alps[16,172]. An attempt was also made to move northward to settle and take control of the metal exchange. This could in turn have led to battle in the Tollense valley in Mecklenburg region that included several thousand warriors from south Scandinavia, northern Germany and central Europe[173]. However, archaeological and genetic evidence suggests that this expansion was not successful. Genetic data reveals no influx of Urnfield/Knovíz-associated signatures in Scandinavian samples, while archaeological findings indicate that the Nordic Bronze Age Culture continued its southward expansion. These geopolitical events, however, would lead to a temporary decline of copper supplies to the north from the Italian Alps[16], reflected in generation of highly worn Nordic swords[172], and probably also stimulated a search for new copper sources.

Thus the Urnfield expansion marked also a counter movement, the rise of the Atlantic trade network, which began facilitated by the movement of copper from Iberia to the British Isles and southern Scandinavia [16]. Urnfield communities across central Europe persisted and kept expanding on a smaller scale in the Iberian Peninsula, as reflected in the Urnfield burials of the northeastern Peninsula, as well as in the iconography of the Late Bronze Age warrior stelae and some metalwork in the west[155,162]. In terms of the British and Irish Isles the genetic and archaeological evidence indicates that this process had an impact mainly on southern England, the Channel Zone most strongly, but only in more limited domains in northern Britain and Ireland[78,174,175]. In Britain the intensification of characteristically Atlantic Bronze Age connections are prominent, links to Northwest France and the western Iberian Peninsula, reflected in both the distribution of artefact types and the circulation of metals[12,176,177]. Ireland seems to have expanded with a flourishing Late Bronze Age, that was connected to both Scandinavia and Iberia [178,179].

Urnfield influences were thus both widespread and long-lived in Europe, and the expansion of its various parts has therefore remained a topic for debate [180,181].

However, climatic change and new influences from the steppe in the form of Cimmerians that introduced new burials rituals, new horse and wagon technologies, and new types of warfare would soon undermine Urnfield communities, both their economy and their metalwork, as Iron became dominant (see Figure 102-103 in [51]). These changes would lead to the formation of the Hallstatt C Culture (800/750-600/550 BC), followed by Hallstatt D (600-450 BC), and La Tène from 450 BC to the beginning of the common era and the Roman march towards empire.

***Hallstatt and La Tène Cultures***

The end of the Bronze Age and the beginning of the Iron Age proper coincides with a major climatic event, the little Ice Age starting 800 BC (see Figure 12 in [51]). It would impact Urnfield farming regimes dramatically, and lead to new economic practices with a heavier stress on upland regions. Thus, the Hallstatt C Culture in western Europe is characterized by a return to the use of large tumulus burials with the deceased in wooden chambers, as well as the introduction of new horse gear, wagons and weapons, in part inspired by the influence from eastern Europe of Cimmerian warrior nomads (see pages 201-209 in [51], [182]). Also, the full-scale implementation of iron technology had a huge transformative impact. Western Hallstatt thus saw expansion along the Rhine and into southern England, Britain and Ireland[59,60]. To the east we see also the formation of large defended chiefly or royal settlements and a developed industrial production (see pages 210-249 in [51]). These developments would continue and accelerate during Hallstatt D which is characterized by a much stronger influence from the so-called Orientalizing period in the Mediterranean, represented by the Etruscans, and the expanding Greek colonies[183,184]. Connections to northern Europe at the same time faded.

The formation of the Hallstatt D royal centers, sometimes characterized as a form of proto-urbanisation[185], marked the continuation of a development towards political centralization and economic intensification that started in east Hallstatt and now took over also in the west, where former Hallstatt C centers declined or came under the political regime of the new Hallstatt D centers [184,186]. However, their flourishing came to an end when social transformations rooted in resistance to economic exploitation from royal centers in connection with demographic surplus, would lead to their downfall 500/450 BC, represented by La Tène Culture.

La Tène Culture emerged on the closer peripheries of the Hallstatt D royal centers, as a counterreaction to their power, which they crushed followed by large scale migrations during the following centuries (see Figure 156 in [51]), towards the east and south into northern Italy, the Balkans, but probably also to the west in the British Isles, even if this is debated (see Figure 176 in [51], [187]). However, their material Culture expanded widely throughout the known world from northern to central Europe, in much the same way as during the Urnfield period. As during the Urnfield period, we witness an intensification of farming during and after the expansion phase[188].

After the termination of the migrations, huge permanent proto urban settlements were formed, so-called Oppida, from around 200/150 BC (see pages 314-358 in [51]), that ended only with Roman occupation. Thus, they can be considered in part as a forced mobilization and centralization to resist Roman expansion. But they were also industrial centers whose products would be spread widely across Europe, much in the same way as during the Urnfield Culture[189].

If we use the archaeological record to make inferences from the known to the unknown, we can make inferences about migrations in prehistory. Comparing La Tène migrations that are well documented in historical records, with the Late Bronze Age Urnfield expansion, where we have no written records, we must conclude that migrations were a driving force behind the Urnfield expansion supported by agrarian and technological advances, as well as new religion.

**Conclusion**

From the Urnfield Culture to the end of La Tène we observe a recurring economic and social dynamic between periods of elite dominance (often warriors) and periods of communal ideology in burial ritual. Elites are often represented by impressive tumulus burials, with little or no recognition of ‘‘commoners”, they remain ritually invisible, whereas the communal ideology represents commoners/farmers in the form of urnfield cemeteries (Kristiansen 1998: Figure 224). Thus, European societies from the Bronze Age through the Iron Age oscillated between two dominant types of social organization – one based on sedentary farming communities with a ‘democratic’ ideology, and another based on a chiefly warrior ideology with lavish display of burial goods.

We can also describe these regularities as a recurring fight between two competing ideologies: inequality and equality. There were always forces leading to inequality and social hierarchies and opposing forces leading to decreasing inequality and the decline or reduction of hierarchies. Thus, we witness political counteractions against ‘tyranny’ in many parts of Europe in the decades around 500 BC, from Greece leading to the introduction of democracy to central Europe and the downfall of the Hallstatt D royal residences, to Scandinavia and the collapse of Bronze Age elite networks. The period 500-150 BC thus introduced new more egalitarian social formations across Europe, only to be replaced by new hierarchies from the Roman empire onwards.

111. Jordán Cólera, C. *Lengua Y Epigrafía Celtibéricas*. (Prensas de la Universidad de

Zaragoza, Zaragoza, 2019).

112. Jordán Cólera, C. *El Legado Escrito de Los Pueblos Paleohispánicos*. (Prensas de la Universidad de Zaragoza, Zaragoza, 2024).

113. Koch, J. T. Paradigm shift? Interpreting Tartessian as Celtic. in *Celtic from the West: Alternative Perspectives from archaeology, genetics, language and literature* (eds. Cunliffe, B. & Koch, J. T.) 185–302 (Oxbow Books, Oxford/Oakville, 2012).

114. Hamp, E. P. & Adams, D. Q. The expansion of the Indo-European languages: an Indo-Europeanist's evolving view. *Sino-Platonic Papers* **239**, (2013).

115. Koch, J. T. *Common Ground and Progress on the Celtic of the South-Western (S.W.) Inscriptions*. (Centre for Advanced Welsh and Celtic Studies, Aberystwyth, 2019).

116. Rodríguez Ramos, J. Las inscripciones sudlusitano-tartesiassu función, lengua y contexto socio-económico. *Complutum* **13**, 85–95 (2002).

117. Eska, J. F. Comments on John T. Koch's Tartessian-as-Celtic enterprise. *Journal of Indo-European Studies* **42**, 428–438 (2014).

118. Hoz, J. de. Method and methods. in *Palaeohispanic Languages and Epigraphies* 1–24 (Oxford University Press, 2019).

119. Eska, J. F. A salvage grammar of Galatian. *Zeitschrift für celtische Philologie* **60**, 51–64 (2013).

120. Stifter, D. *Cisalpine Celtic. Language, Writing, Epigraphy*. (Prensas de la Universidad de Zaragoza, Zaragoza, 2020).

121. Lambert, P.-Y. *La Langue Gauloise*. (Editions Errance, Paris, 1994).

122. Delamarre, X. *Dictionnaire de la langue gauloise: une approche linguistique du vieux-celtique continental*. (Editions Errance, Paris, 2003).

123. Eska, J. F. The linguistic position of Lepontic. in *Proceedings of the Twenty-Fourth Annual Meeting of the Berkeley Linguistics Society: Special Session on Indo-European*

*Subgrouping and Internal Relations* (eds. Bergen, B. K., Plauché, M. C. & Bailey, A.) 2–11 (Berkeley Linguistics Society, Berkeley, CA, 1998).

124. Freeman, P. M. *The Galatian Language: A Comprehensive Survey of the Language of the Ancient Celts of Greco-Roman Asia Minor*. (Edwin Mellen Press, Lampeter, 2001).
125. Cottam, E., de Jersey, P., Rudd, C. & Sills, J. *Ancient British Coins*. (Chris Rudd, Aylsham, Norfolk, 2010).
126. McManus, D. *A Guide to Ogam*. (An Sagart, Maynooth, 1991).
127. Wodtko, D. S. Language contact in Lusitania. *International Journal of Diachronic Linguistics and Linguistic Reconstruction* **6**, 1–48 (2009).
128. The labyrinth of continental Celtic. *Proceedings of the British Academy* **65**, 497–538 (1977).
129. Untermann, J. Lusitanisch, Keltiberisch, Keltisch. *Veleia* **2-3**, 57–76 (1986).
130. Ballester, X. 'Páramo' o del problema de la */p/ en celtoide. *Studi Celtici* **3**, 45–56 (2004).
131. Prósper, B. M. The Lusitanian oblique cases revisited: New light on the dative endings. in *Curiositas nihil recusat. Studia Isabel Moreno Ferrero dicata: estudios dedicados a Isabel Moreno Ferrero* (eds. González Iglesias, J. A., Méndez Dosuna, J. V. & Prósper, B. M.) 427–442 (Ediciones Universidad de Salamanca, Universidad de Salamanca, 2021).
132. Renfrew, C. *Archaeology and Language: The Puzzle of Indo-European Origins*. (Jonathan Cape, London, England, 1987).
133. Patterson, N. What genetics can say about Iron Age and Bronze Age Britain. in *Presenting counterpoints to the dominant terrestrial narrative of European prehistory* (eds. Koch, J. T., Fauvelle, M., Cunliffe, B. & Ling, J.) 187–190 (Oxbow Books, Oxford, 2025).

134. Abercromby, J. *A Study of the Bronze Age Pottery of Great Britain & Ireland and Its Associated Grave-Goods*. (Clarendon press, Oxford, 1912).

135. Harbison, P. The coming of the Indo-Europeans to Ireland: an archaeological viewpoint. *The Journal of Indo-European Studies* **3**, 101–119 (1975).

136. Koch, J. T. Convergence *in situ*: the formation of the Indo-European branches and the Bronze–Iron transition. in *Presenting counterpoints to the dominant terrestrial narrative of European prehistory. Maritime Encounters 1* (eds. Koch, J. T., Fauvelle, M., Cunliffe, B. & Ling, J.) 203–220 (Oxbow Books, Oxford, 2025).

137. McCone, K. R. *Towards a Relative Chronology of Ancient and Medieval Celtic Sound Change*. (Department of Old and Middle Irish, St Patrick's College, Maynooth, 1996).

138. Isaac, G. R. *Studies in Celtic Sound Changes and Their Chronology*. (Innsbrucker Beiträge zur Sprachwissenschaft, Innsbruck, 2007).

139. Garrett, A. Convergence in the formation of Indo-European subgroups: phylogeny and chronology. in *Phylogenetic methods and the prehistory of languages* (eds. Forster, P. & Renfrew, C.) 139–151 (McDonald Institute for Archaeological Research, Cambridge, 2006).

140. Pare, C. F. E. Bronze and the Bronze Age. in *Metals make the world go round: supply and circulation of metals in Bronze Age Europe* (ed. Pare, C. F. E.) 1–37 (Oxbow Books, Oxford, 2000).

141. Koch, J. T. Out of the flow and ebb of the European Bronze Age: heroes, Tartessos, and Celtic. in *Celtic from the West 2: rethinking the Bronze Age and the arrival of Indo-European in Atlantic Europe* (eds. Koch, J. T. & Cunliffe, B.) 101–146 (Oxbow Books, Oxford, 2013).

142. Needham, S. P. Encompassing the sea: 'maritories' and Bronze Age maritime interaction. in *Bronze Age Connections: Cultural Contact in Prehistoric Europe* (ed.

Clark, P.) 12–37 (Oxbow Books, Oxford, 2009).

143. Burlot, A. Chalcolithic and Bronze Age Atlantic connections c. 2500–800 BC. in *Presenting counterpoints to the dominant terrestrial narrative of European prehistory, Maritime Encounters 1* (eds. Koch, J. T., Fauvelle, J. T., Cunliffe, B. & Ling, J.) 31–48 (Oxbow Books, Oxford, 2025).

144. Williams, A. & Le Carlier de Veslud, C. Boom and bust in Bronze Age Britain: major copper production from the Great Orme mine and European trade, *c.* 1600–1400 BC. *Antiquity* **93**, 1178–1196 (2019).

145. Williams, R. A. *Boom and Bust in Bronze Age Britain: The Great Orme Copper Mine and European Trade*. (Archaeopress, Oxford, 2023).

146. Stos-Gale, Z. A. & Ling, J. Archaeology and science: impact of lead isotope analyses on the archaeological discourse of metal trade for the Scandinavian and British communities in the 3rd–1st millennia BC. in *Presenting counterpoints to the dominant terrestrial narrative of European prehistory, Maritime Encounters 1* (eds. Koch, J. T., Fauvelle, M., Cunliffe, B. & Ling, J.) 147–166 (Oxbow Books, Oxford, 2025).

147. Hunt-Ortiz, M. A. *et al.* Late Bronze Age copper mining in southern Iberia: preliminary results of fieldwork at Las Minillas (Granja de Torrehermosa, Badajoz, Spain). in *Presenting counterpoints to the dominant terrestrial narrative of European prehistory, Maritime Encounters 1* (eds. Koch, J. T., Fauvelle, M., Cunliffe, B. & Ling, J.) 167–186 (Oxbow Books, Oxford, 2025).

148. Koch, J. T. Phoenicians in the west and break-up of the Atlantic Bronze Age. in *Celtic from the West 3. Atlantic Europe in the metal ages. Questions of shared language* (eds. Koch, J. T. & Cleary, K.) 431–476 (Oxbow Books, Oxford, 2016).

149. Schmidt, K. H. *Celtic: A Western Indo-European Language?* (Innsbrucker Beiträge zur Sprachwissenschaft, Innsbruck, 1996).

150. Schmidt, K. H. Armenian and Celtic. Towards a New Classification of Early Indo-European Dialects. *Bulletin of the Georgian Academy of Sciences* **175**, 199–203 (2007).
151. Isaac, G. R. The origins of the Celtic languages: language spread from east to west. in *Celtic from the west. Alternative perspectives from archaeology, genetics, language, and literature* (eds. Cunliffe, B. & Koch, J. T.) 153–167 (Oxbow Books, Oxford, 2010).
152. Prósper, B. M. The inscription of Cabeço das Fráguas revisited. Lusitanian and Alteuropäisch populations in the West of the Iberian Peninsula. *Trans. Philol. Soc.* **97**, 151–184 (1999).
153. Luján, E. R. Language and writing among the Lusitanians. in *Paleohispanic languages and epigraphies* (eds. Sinner, A. G. & Velaza, J.) 304–334 (Oxford University Press, Oxford, 2019).
154. Bosch-Gimpera, P. The two Celtic waves in Spain. *Proceedings of the British Academy* **26**, 1–126 (1942).
155. Brandherm, D. Westward ho? Sword-bearers and all the rest of it. in *Celtic from the West 2: Rethinking the Bronze Age and the arrival of Indo-European in Atlantic Europe* (eds. Koch, J. T. & Cunliffe, B.) 147–156 (Oxbow Books, Oxford, 2013).
156. *Bronze Age Rock Art in Iberia and Scandinavia: Words, Warriors, and Long-Distance Metal Trade*. (Oxbow Books, Oxford, 2024).
157. McCone, K. R. *The Celtic Question: Modern Constructs and Ancient Realities*. (Dublin Institute for Advanced Studies, Dublin, 2008).
158. Watkins, C. W. Two Celtic notes. Studia Celtica et Indogermanica. in *Festschrift für Wolfgang Meid* (eds. Anreiter, P. & Jerem, E.) 539–543 (Archeolingua, Budapest, 1999).
159. Isaac, G. R. Insular Celtic vs Gallo-Brittonic: an empirical and methodological question. in *Celtic connections: Papers from the Tenth International Congress of Celtic Studies,*

*Edinburgh, 1995, 2. Archaeology, numismatics, historical linguistics* (eds. Gillies, W. & Harding, D. W.) 190–202 (University of Edinburgh, Edinburgh, 2005).

160. De Bernardo Stempel, P. Language and historiography of the Celtic peoples. in *Celtes et Gaulois, l'archéologie face à l'histoire: Celtes et Gaulois dans l'histoire, l'histographie et l'idéologie moderne* (ed. Rieckhoff, S.) 33–56 (Bibracte, Glux-en-Glenne, 2006).
161. Matasović, R. *Etymological Dictionary of Proto-Celtic*. (Brill, Leiden, Netherlands, 2009).
162. Montero Ruiz, I., Hunt Ortiz, M. A. & Santos Zalduegui, J. F. El depósito de la Ría de Huelva: procedencia del metal a través de los resultados de análisis de isótopos de plomo. in *El hallazgo leones de Valdevimbre y los depósitos del Bronce Final Atlántico en la Península Ibérica* (eds. Celis, J., Delibes de Castro, G., Fernández Manzano, J. & Grau Lobo, L.) 194–209 (Estudios y Catálogos Museos de Castilla y León, Léon, 2007).
163. Mallory, J. P. From the steppe to Ireland: The impact of aDNA research. in *The Indo-European Puzzle Revisited* 129–145 (Cambridge University Press, 2023).
164. Schrijver, P. *Language Contact and the Origins of the Germanic Languages*. (Routledge, London, England, 2014).
165. Mallory, J. P. *The Origins of the Irish*. (Thames & Hudson, 2013).
166. O'Brien, W. Language shift and political context in Bronze Age Ireland: Some implications of hill fort chronology. in *Celtic from the West 3: Atlantic Europe in the metal ages: Questions of shared language (Celtic Studies Publications 19)* (eds. Koch, J. T. & Cunliffe, B.) 219–246 (Oxbow Books, Oxford, 2016).
167. Sørensen, M. L. S. & Rebay-Salisbury, K. *Death and the Body in Bronze Age Europe: From Inhumation to Cremation*. (2023).
168. Unger, J. Obříství, a Late Bronze Age Port of Trade in Central Bohemia. *Studia Hercynia* **20**, 7–28 (2016).

174. Gerloff, S. *Atlantic Cauldrons and Buckets of the Late Bronze and Early Iron Ages in Western Europe: With a Review of Comparable Vessels from Central Europe and Italy*. (Franz Steiner Verlag Wiesbaden gmbh, 2010).

175. Eogan, G. Ideas, People and Things: Ireland and the External World during the Later Bronze Age. in *Ireland in the Bronze Age, Proceedings of the Dublin Conference, April 1995* 128–135 (Stationery Office, Dublin, 1995).

176. Berger, D. *et al.* The Salcombe metal cargoes: New light on the provenance and circulation of tin and copper in Later Bronze Age Europe provided by trace elements

and isotopes. *J. Archaeol. Sci.* **138**, 105543 (2022).

177. Needham, S. & Bowman, S. Flesh-hooks, technological complexity and the Atlantic Bronze Age feasting complex. *European Journal of Archaeology* **8**, 93–136 (2005).
178. Almagro-Gorbea, M. Ireland and Spain in the Bronze Age. in *Ireland in the Bronze Age: Proceedings of the Dublin Conference, April 1995* 136–148 (Office of Public Works, Dublin, 1995).
179. Waddell, J. *The Prehistoric Archaeology of Ireland*. (1998).
180. Dzięgielewski, K., Gawlik, A. & Przybyła, M. S. *Migration in Bronze and Early Iron Age Europe*. (Archeobooks, 2010).
181. Falcheti Peixoto, R. & Iacono, F. Urnfield Bronze Connections: Rethinking Late Bronze Age Mobility. *OCNUS* **30**, 149–172 (2022).
182. Metzner-Nebelsick, C. *Der 'Thrako-Kimmerische' Formenkreis aus der Sicht der Urnenfelder- und Hallstattzeit im südöstlichen Pannonien*. (Verlag Marie Leidorf, 2002).
183. Egg, M. Die Halltstattkulturen und Italien während der älteren Eisenzeit. **127**, 18–61 (2021).
184. Krausse, D. & Beilharz, D. *'Fürstensitze' und Zentralorte der frühen Kelten: Abschlusskolloquium des DFG-Schwerpunktprogramms 1171 in Stuttgart, 12.-15. Oktober 2009*. (2010).
185. Fernandez-Götz, M. & Krausse, D. Rethinking Early Iron Age urbanisation in Central Europe: the Heuneburg site and its archaeological environment. *Antiquity* **87**, 473–487 (2013).
186. Fernández-Götz, M., Wendling, H. & Winger, K. *Paths to Complexity: Centralisation and Urbanisation in Iron Age Europe*. (2014).
187. Vannier, E. The Funerary Architecture of the La Tène Period in North-western Gaul and

Southern Britain. *Proceedings of the Prehistoric Society* **86**, 285–304 (2020).

188. Hornung, S. *Produktion . Distribution - Ökonomie: Siedlungs- und Wirtschaftsmuster der Latènezeit; Akten des internationalen Kolloquiums in Otzenhausen, 28. - 30. Oktober 2011*. (2014).

189. Fernández-Götz, M. Urbanization in Iron Age Europe: Trajectories, Patterns, and Social Dynamics. *Journal of Archaeological Research* **26**, 117–162 (2017).
